# A novel phospholipid mimetic targeting LRH-1 ameliorates colitis

**DOI:** 10.1101/2020.09.01.278291

**Authors:** Suzanne G. Mays, Emma H. D’Agostino, Autumn R. Flynn, Xiangsheng Huang, Guohui Wang, Xu Liu, Elizabeth J. Millings, C. Denise Okafor, Anamika Patel, Michael L. Cato, Jeffery L. Cornelison, Dianna Melchers, René Houtman, David D. Moore, John W. Calvert, Nathan T. Jui, Eric A. Ortlund

## Abstract

Phospholipids are ligands for nuclear hormone receptors (NRs) and regulate transcriptional programs relevant to normal physiology and disease. Here, we demonstrate that mimicking phospholipid-NR interactions greatly improves agonists of liver receptor homolog-1 (LRH-1), a promising therapeutic target for diabetes and colitis. Conventional LRH-1 modulators partially occupy the binding pocket, leaving vacant a region important for phospholipid binding and allostery. Therefore, we constructed a set of hybrid molecules with elements of natural phospholipids appended to a synthetic LRH-1 agonist. The phospholipid-mimicking group improves binding affinity, increases LRH-1 transcriptional activity, promotes coregulator recruitment, and interacts with the targeted LRH-1 residues in crystal structures. The best new agonist markedly improves colonic histopathology and disease-related weight loss in a humanized LRH-1 murine T-cell transfer model of colitis. This is the first evidence of *in vivo* efficacy for an LRH-1 modulator in colitis, a leap forward in agonist development.

## Introduction

Phospholipids (PLs) play many roles in physiology, such as forming lipid bilayers, signaling for apoptosis, and activating G-protein coupled receptors. PL are labile and are synthesized and catabolized in response to stimuli such as fed or fasting states, stress, and circadian rhythms,^1,2^ yielding metabolites that perform specialized functions. Phosphatidylcholines (PCs) are a subtype of PL that are vital for maintaining integrity and curvature of eukaryotic cellular membranes^3^ and also play important roles as cell signaling mediators. Catabolism of PCs produces bioactive molecules, such as arachidonic acid,^4^ and generates labile methyl groups used for DNA methylation and synthesis of nucleotides and amino acids.^5^ Additionally, certain PCs exert broad effects on metabolic homeostasis by acting as ligands for nuclear hormone receptors (NRs).^1,6–9^ As NR ligands, PCs represent a novel, understudied class of hormone.

Particularly strong evidence supports a role for PCs as ligands of liver receptor homolog-1 (LRH-1). LRH-1 is an orphan NR that regulates cholesterol homeostasis, metabolism, proliferation, and intestinal inflammation, garnering attention as a novel therapeutic target for diseases such as diabetes, cancer, and inflammatory bowel diseases.^9–15^ LRH-1 binds a wide range of PLs *in vitro*, and it is activated exogenously by medium-chained, saturated PCs.^9,16^ Moreover, several models have shown a relationship between endogenous PC levels and LRH-1 activity. Diet-induced depletion of PCs in mice induces an “antagonistic” pattern of gene expression in the liver similar to LRH-1 liver-specific knockout mice.^17^ Similarly, diet-induced depletion of PCs in *C. elegans* inhibits the worm LRH-1 ortholog.^18^ In human hepatocytes, LRH-1 senses PCs generated via methyl metabolism to control beta-oxidation of fatty acids and mitochondrial biogenesis.^19^ Finally, a PC agonist of LRH-1 (dilauroylphosphatidylcholine, DLPC) induces LRH-1-dependent anti-diabetic effects in mouse models of insulin resistance.^9^ The anti-diabetic effects occur via repression of *de novo* lipogenesis in the liver.

The sensitivity of LRH-1 to PC levels suggests a regulatory circuit connecting PC availability to LRH-1-controlled gene expression. However, this circuit has been difficult to elucidate. Natural PCs are insoluble, rapidly metabolized, and unlikely to be LRH-1-selective, which makes them difficult to work with in the laboratory. These poor pharmacological properties also limit the use of PCs as therapeutics. We therefore designed a set of “PL-mimics” by fusing PL headgroups (or phosphate bioisosteres) to the synthetic LRH-1 agonist RJW100 (Figure 1a-b).^20^ These hybrid molecules contain the hexahydropentalene (6HP) core from RJW100, appended with a polar moiety that mimics a PL headgroup at position R^4^ (Figure 1a-b).^20^ Based on structural analyses, we hypothesized that the modified R^4^ groups would mimic PL interactions in the LRH-1 binding pocket and stimulate receptor activity. Indeed, the agonists activate LRH-1 robustly *in vitro* and promote distinctive patterns of coregulator protein recruitment. We present new crystal structures showing that the agonists are anchored deeply in the pocket via the 6HP core and that the added polar groups make PL-like contacts at the mouth of the pocket as designed. The best of this class is active *in vivo*, suppressing lipogenic genes in the liver and inducing steroidogenic genes in the gut in separate mouse studies. In a murine model of chronic colitis, the best new agonist profoundly reduces inflammation and disease-related weight loss. This work demonstrates that strategic targeting of the PC binding site improves activity, elucidates the mechanism of action of these compounds, and provides the first *in vivo* evidence of efficacy in colitis for a synthetic LRH-1 modulator.

**Figure 1.**
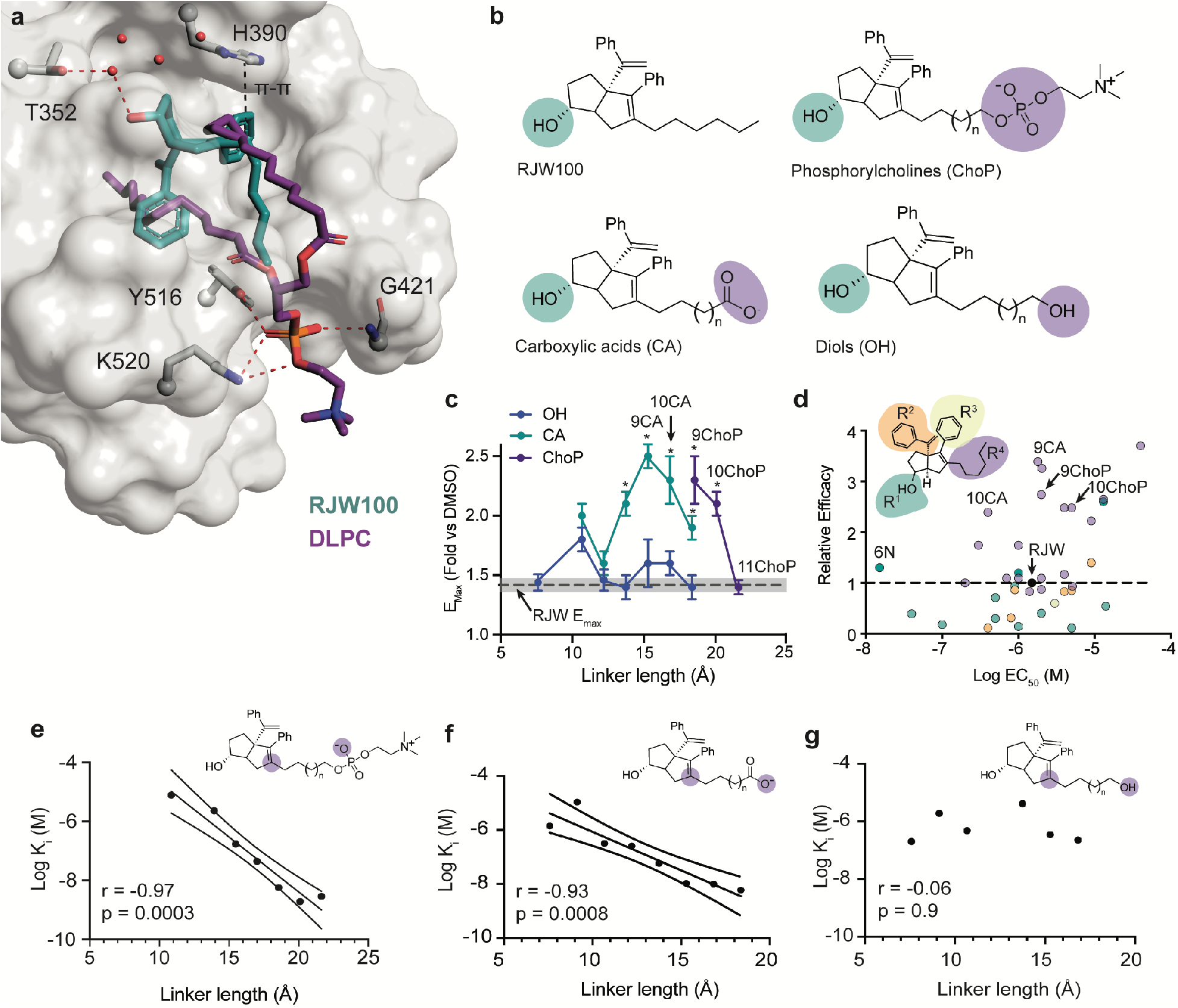
Design, binding, and activity of PL-mimetics. **a**, LRH-1 ligand binding pocket (LBP) from PDB 5L11, showing the binding modes of synthetic agonist RJW100 (teal sticks) and phospholipid agonist DLPC (PDB 4DOS, purple sticks). Key interactions made by each agonist are highlighted. **b**, Phosphorylcholines or carboxylic isosteres were conjugated to the RJW100 core via alkyl linkers of 4-11 carbons.^20^ Diols with alkyl linkers of 4-11 carbons were synthesized for comparison with PL mimics. **c**, Plot of E_max_ values from luciferase reporter assays as a function of linker length. Each point represents the mean +/-SEM from 2-3 experiments. Compounds that did not activate sufficiently for E_max_ calculation are omitted. *Dotted line* is the mean E_max_ of RJW100; grey shading indicates SEM. *, p< 0.05 *versus* RJW100 by two-way ANOVA followed by Sidak’s multiple comparisons test. **d**, Activity profiles of the PL-mimics *versus* related LRH-1 agonists. *Inset*, RJW100 chemical structure, showing the color scheme for the dots in the plot (colored according to the site of modification). Relative efficacy is E_max_ relative to RJW100, calculated as described in Methods. **e-g**, Binding affinity (K_i_) plotted as a function of linker length for ChoPs *(**e**)*, CAs *(**f**)*, and diols *(**g**)*. *Lower insets*, Pearson correlation coefficients (r) and p-values. *Upper insets*, illustration of the portion of R^4^ used to measure linker length (distance between the two purple spheres).

## Results

### Structure-guided design of PL-mimics

We exploited information from LRH-1 crystal structures to design a set synthetic PL-mimetics.^7,20–23^ PLs bind LRH-1 differently than most NR ligands: rather than being fully engulfed in the pocket, a portion of the headgroup protrudes into the solvent (Figure 1a). Lipid phosphates coordinate polar residues near the pocket mouth, while the acyl chains line the pocket interior, making numerous hydrophobic contacts (Figure 1a). In contrast, the synthetic LRH-1 agonist RJW100 binds deep in the pocket (Figure 1a). We previously focused on strengthening deep pocket interactions made by RJW100, resulting in the discovery of the agonists 6N and 6Na (Figure S1).^24,25^ For the PL-mimics, we used an alternative approach, seeking to promote interactions near the mouth of the pocket. Notably, the alkyl “tail” of RJW100 (position R^4^) overlaps with the PL fatty acyl chains and follows a trajectory toward the pocket mouth (Figure 1a). Based on this observation, we synthesized sets of compounds modified at R^4^ with alkyl tails of varied lengths that terminate in polar, PL-mimicking groups (Figure 1b, S2-S4).^20^

### Modifications improve binding affinity and activity

The addition of a phosphorylcholine (ChoP) or carboxylic acid (CA) to position R^4^ on the RJW100 scaffold dramatically increases LRH-1 transcriptional activity, depending on the length of the alkyl linker connecting the bicyclic core to the modified group.^20^ Compounds containing linkers of 9-10 carbons are the strongest activators for both classes, increasing E_max_ nearly two-fold over RJW100 (Figure 1c). The optimal linker is 18-20 Å from the bicyclic core to the terminal polar group for the ChoPs and 15-17 Å for the CAs (Figure 1c, Table S1). Compounds with short alkyl linkers (4-5 carbons) do not significantly activate the receptor above baseline at doses up to 30 μM. Diverse modifications elsewhere on the RJW100 scaffold^24^ affect potency (EC_50_) but rarely increase E_max_ (Figure 1d), demonstrating an unique ability for the R^4^ modifications to increase activity levels. The PL-mimicking groups also improve binding affinity in a linker length-dependent manner (Figure 1f-g, S2-S4). ChoPs and CAs with longer linkers (9-11 carbons) have the highest affinities, with K_i_ values between 2-11 nM (Figure 1e-g, Table S1). To test whether the improved affinity and activity are caused by the longer alkyl linker or the terminal polar group, we synthesized and tested a set of diols (terminally hydroxy-modified at R^4^) with varying linker lengths (Figure 1b). Longer linkers do not improve binding or activity of the diols compared to RJW100, suggesting that the charged polar group is the driver of the improved properties (Figure 1d, 1g, S4-S5). Together, these data suggest that PL-mimicking groups form productive interactions with LRH-1, greatly increasing transcriptional output.

### Structural basis for improved binding and activity

To understand the mechanism of action of the PL-mimics, we determined X-ray crystal structures of the LRH-1 ligand binding domain (LBD) bound to either 10CA or 9ChoP at resolutions of 2.3 and 2.55 Å, respectively (Figure 2a-b, Table 1). Both structures depict the ligand bound at a single site in the ligand binding pocket (Figure 2a-b). The peptide from the transcriptional intermediary factor 2 (Tif2) coregulator, added to stabilize the complex, is bound to the activation function surface (AFS), comprised of portions of helices 3, 4, and the activation function helix (AF-H) (Figure 2a-b).

**Figure 2.**
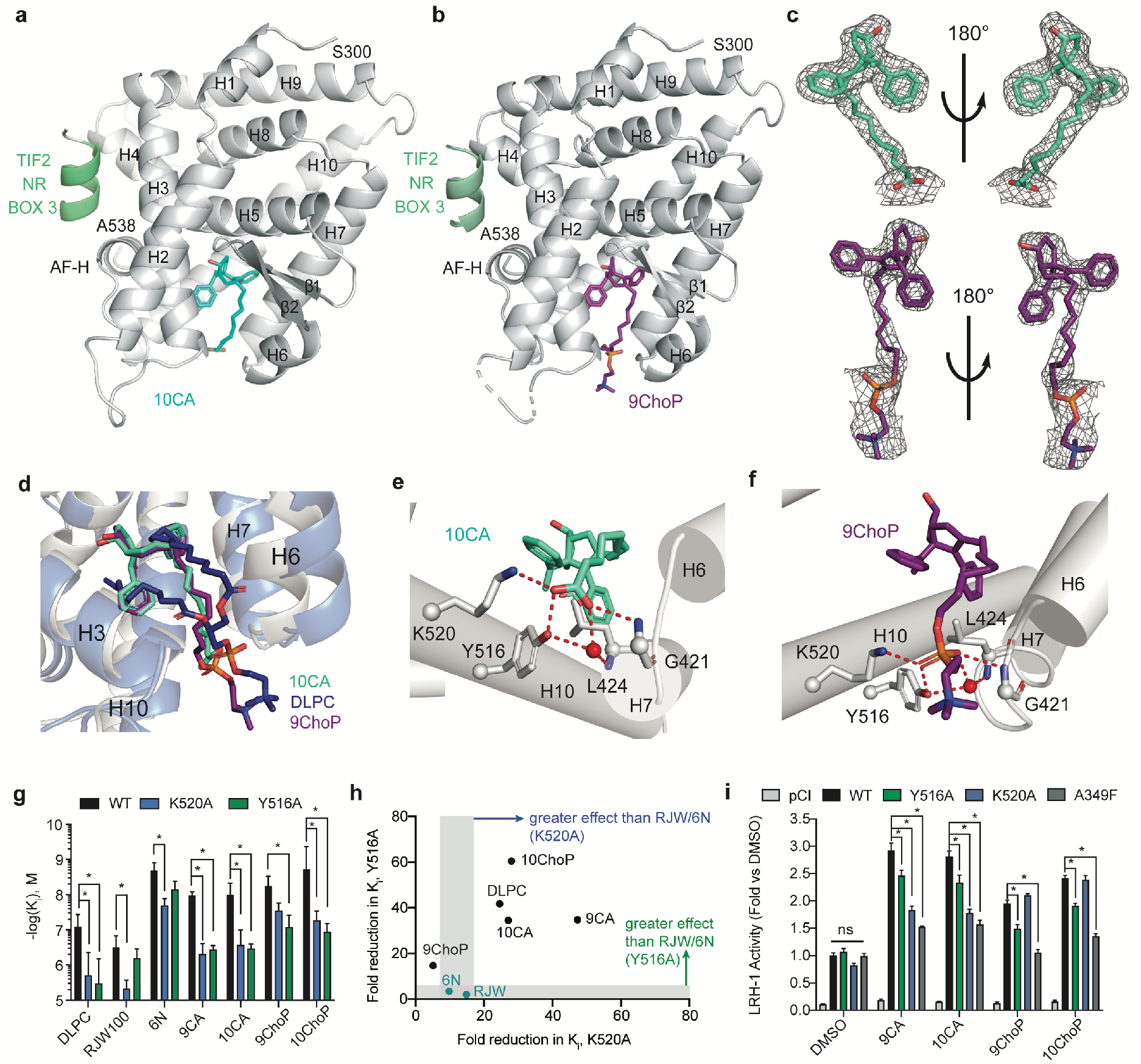
Phospholipid mimetics make PL-like interactions. **a-b**, Crystal structures of LRH-1 LBD with (***a***) 10CA (PDB: 7JYD) or (***b***) 9ChoP (PDB: 7JYE) and a fragment of the Tif2 coregulator. *H*, helix. **c**, Electron density surrounding 10CA (*top*) and 9ChoP (*bottom*). Maps are omit F_o_-F_c_, contoured at 2σ. **d**, Superposition of three ligands from LRH-1 structures: 10CA (cyan sticks), 9ChoP (purple sticks), or DLPC (dark blue sticks, from PBD 4DOS^7^). 10CA and DLPC protein backbones are shown as grey and light blue cartoons, respectively. **e-f**, Close-up of the interactions made by the terminal polar groups of 10CA (***e***) and 9ChoP (***f***). Hydrogen bonds are shown as red dotted lines; water molecule as a red sphere. **g**, Effects of K520A or Y516A mutations on binding affinity. Each bar represents the mean +/-95% CI from two experiments conducted in quadruplicate. *, p < 0.05 by two-way ANOVA followed by Dunnett’s test for multiple comparisons. (h) Fold changes in K_i_ for mutant *versus* WT LRH-1. Grey shading indicates effects of the mutations on RJW100 and 6N binding, which do not contact the mutated residues. **i**, Luciferase reporter assays with LRH-1 containing K520A or Y516A mutations. The A349F mutation occludes the pocket and was a negative control. Cells were treated with 30 μM of each compound for 24 hours prior to measurement of luciferase signal. *, p < 0.05 *versus* WT LRH-1 treated with each agonist; (two-way ANOVA followed by Dunnett’s test for multiple comparisons).

**Table 1:** X-ray data collection and refinement statistics.

| **Data collection** | LRH-1 + 10CA + TIF2 | LRH-1 + 9ChoP + TIF2 |
| --- | --- | --- |
| Space group | P3<sub>2</sub>21 | P3<sub>2</sub>21 |
| Cell dimensions |  |  |
| $a, b, c$ (Å) | 88.4, 88.4, 105.7 | 89.2, 89.2, 106.5 |
| $\alpha, \beta, \gamma$ (°) | 90, 90, 120 | 90, 90, 90 |
| Resolution (Å) | 38.27 - 2.30 (2.38 - 2.30) | 34.21 – 2.55 (2.64-2.55) |
| $R_{\text{pim}}$ | 0.045 (0.305) | 0.074 (0.542) |
| $I / \sigma I$ | 22.7 (1.79) | 13.8 (1.31) |
| CC<sub>1/2</sub> | 97.0 (77.6) | 98.7 (57.1) |
| Completeness (%) | 99.8 (98.7) | 99.9 (100.0) |
| Redundancy | 15.8 (15.3) | 13.8 (13.8) |
| **Refinement** |  |  |
| Resolution (Å) | 2.30 | 2.55 |
| No. reflections | 21639 | 16426 |
| $R_{\text{work}} / R_{\text{free}}$ (%) | 21.3 / 23.3 | 17.6 / 20.1 |
| No. atoms |  |  |
| Protein | 2037 | 2061 |
| Water | 38 | 18 |
| Twin Law | n/a | -h, -k, l |
| B-factors |  |  |
| Protein | 66.3 | 63.7 |
| Ligand | 52.6 | 65.0 |
| Water | 60.2 | 49.7 |
| R.m.s. deviations |  |  |
| Bond lengths (Å) | 0.002 | 0.001 |
| Bond angles (°) | 0.48 | 0.33 |
| Ramachandran favored (%) | 97.2 | 97.6 |
| Ramachandran outliers (%) | 0 | 0 |
| PDB accession code | 7JYD | 7JYE |

Values in parentheses indicate highest resolution shell.

The electron density surrounding the 10CA and 9ChoP agonists unambiguously indicates their positions in the ligand binding pocket, including the extended alkyl linkers and terminal polar groups (Figure 2c). The 6HP cores, styrene substituents and hydroxyl groups adopt nearly identical positions in the deep pocket to RJW100 (Figure S6a). Remarkably, the alkyl tails of both 10CA and 9ChoP extend toward the mouth of the pocket, with the terminal polar groups close to the DLPC phosphate (seen by superposition with PDB 4DOS,^26^ Figure 2d). The contacts made by the CA and ChoP groups near the mouth of the pocket are very similar to those made by DLPC, including hydrogen bonds with the sidechains of residues Y516 and K520 and with the α-amino group of G421 (Figure 2e-f). The agonists also make a water-mediated hydrogen bond with the α-amino group of L424 via the CA and phosphate groups (also made by DLPC), strengthening their associations with the pocket mouth. Molecular dynamics simulations (MDS) with the 10CA structure show that the interactions with Y516 and K520 persist for 50.2% and 37.6% of a 1 μs simulation, respectively. This is similar to DLPC, which interacts for 49.4% of the time with Y516 and 33.2% with K520. The interaction with the backbone amide of G421 is less stable for 10CA than DLPC (27.0% of the simulation *versus* 48.1% for DLPC).

We were particularly interested in how the PL-mimics affect receptor conformation near the mouth of the pocket, since flexibility there is important for allosteric signaling by DLPC.^16^ Specifically, the helix 6/β-sheet region is used to sense ligands and communicate with the coregulator binding surface to recruit appropriate coregulator proteins.^16^ PL binding expands the pocket mouth relative to apo LRH-1 via a ^~^3 Å displacement of helix 6 by the sn1 acyl chain.^7,23^ 10CA and 9ChoP do not shift helix 6; instead, they displace the C-terminus of helix 10, near the mouth of the pocket, by 2.0-2.3 Å relative to apo LRH-1 (from superposition with PDB 4PLD)^16^ and 3.5-3.9 Å relative to the LRH-1-DLPC structure, Figure S6b). To form hydrogen bonds with Y516 and K520, the R^4^ polar groups of the PL-mimics move toward the C-terminus of helix 10 relative to the DLPC phosphate.

To interrogate the function of the PL-like interactions made in the structures, we mutated key LRH-1 residues and measured the impact on binding affinity and transcriptional activity. Introduction of either a Y516A or K520A mutation decreases binding affinities (increasing K_i_) of longer-tailed CAs and ChoPs by 14-60-fold *versus* WT LRH-1, with the exception of 9ChoP for the K520A mutation (Figure 2g, S6). The mutations also impair DLPC binding, decreasing its affinity by 24-40-fold (Figure 2g, S6). Compounds with the 6HP scaffold that do not contact these residues (RJW100 and 6N)^23,24^ are insensitive or less sensitive to these mutations (Figure 2g-h). In luciferase reporter assays, mutations to Y516A or K520A do not affect basal activity of LRH-1; however, the Y516A mutation reduces activation by all four agonists by 24-33% relative to WT protein (Figure 2i). Compounds 9CA and 10CA are very poor activators of the K520A mutant, reducing its activation by around 60% *versus* WT LRH-1, while 9ChoP and 10ChoP activate it equally well as WT protein. These results demonstrate that one or more of the interactions near the mouth of the pocket contribute to the increased affinity and efficacy conferred by the extended polar group at R^4^.

### Effects on LRH-1 conformation and coregulator binding

To understand how the PL-mimics switch LRH-1 into the active state, we investigated how activating *versus* non-activating CAs affect LRH-1 conformation in solution using hydrogen-deuterium exchange mass spectrometry (HDX-MS). We used four CAs for HDX-MS, two that bind LRH-1 but are inactive (4CA and 5CA) and the two most active agonists in this series (9CA and 10CA). Relative to the inactive compounds, 9CA and 10CA stabilize the LRH-1 activation function surface (AFS), a dynamic region in the LBD that serves as a binding site for coregulator proteins (Figure 3a, Figure S7). Stabilized regions in the AFS include helix 3 and the loop preceding the activation function helix (pre-AF-H loop) (Figure 3a-c). 10CA also stabilizes the AF-H and part of helix 4 relative to 4CA (Figure 3a-b).

**Figure 3.**
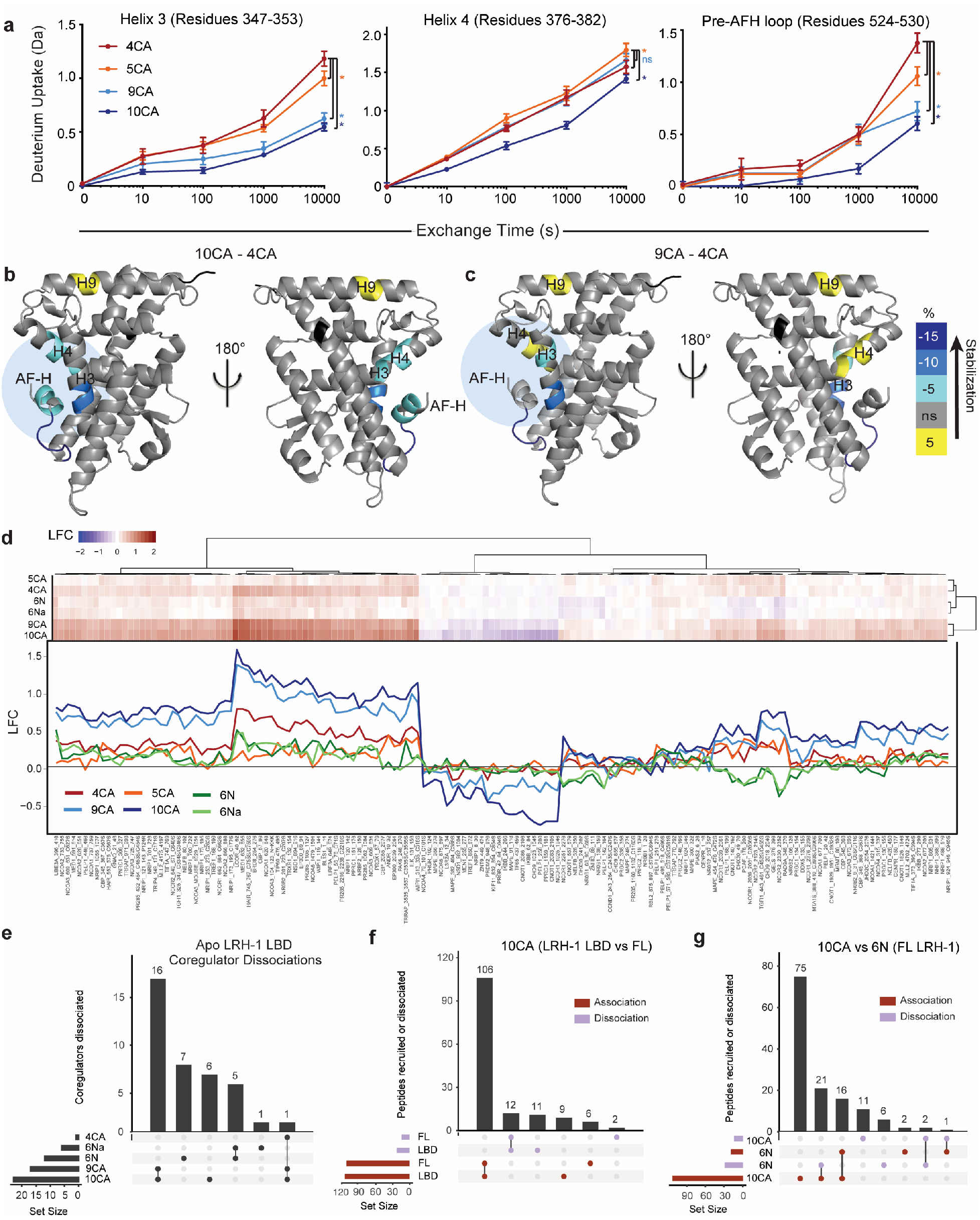
Longer-tailed PL mimetics stabilize the AFS and alter coregulator binding. **a**, Plots of deuterium uptake over time from HDX-MS experiments at sites in the LRH-1 AFS. Each point represents the mean +/-SD for three replicates. Slower deuterium exchange indicates greater stability. *, p < 0.05. *versus* 4CA by two-way ANOVA followed by Holm-Sidak’s test for multiple comparisons. **b-c**, Differences in deuterium uptake for 10CA-LRH-1 (***b***) and LRH-1-9CA (***c***) *versus* LRH-1-4CA mapped to the structure. *Blue circles*, AFS. *Scale bar* indicates the degree of stabilization (teal and blue) or destabilization (yellow) by 9CA or 10CA. *Black*, unmapped regions. *H*, helix. **d**, Heatmap and corresponding raw traces showing log fold change (LFC) coregulator binding relative to apo-LRH-1-LBD. **e-g**, Upset plots showing overlaps in coregulator binding events affected by at least 1.5-fold. Horizontal bars represent the number of events in each set, and vertical bars indicate numbers of events that overlap in between the groups that have filled-in circles below them. For example, the first vertical bar in ***e*** shows that a set of 16 coregulator peptides dissociate from LRH-1 in the presence of 9CA and 10CA that are not affected by 6N and 6Na. **g**, Upset plot comparing overlap in coregulator recruitment events by 10CA bound to full-length LRH-1 (FL) or LRH-1 LBD. Red bars indicate coregulators that associated to the 10CA-LRH-1 complex, and purple bars indicate coregulator dissociation. **h**, Upset plot highlighting an opposite recruitment pattern by compounds 10CA and 6N bound to FL-LRH-1.

The stabilization of the AFS observed with HDX-MS suggests that 9CA and 10CA may affect coregulator associations. To test this, we conducted coregulator profiling using the Microarray Assay for Real-time Coregulator-Nuclear Receptor Interactions (MARCoNI), a microarray that quantifies associations of 64 coregulators from a library of 154 peptides containing NR interaction motifs.^27^ We first investigated how long-*versus* short-tailed compounds affect coregulator binding to purified LRH-1 LBD, using the same CAs as the HDX-MS experiments. Since LRH-1 copurifies with PL from *E. coli* in the pocket, thought to act as weak agonists,^26,28,29^ we generated apo LRH-1 LBD by stripping the *E*. coli PL and refolding the receptor^26^ (Figure S9). Apo LRH-1 LBD preferentially binds peptides from known LRH-1 corepressors (*i.e*. nuclear receptor corepressor 1 (NCOR1), nuclear receptor subfamily 0 group B member 1 (NR0B1, also called DAX-1), and nuclear receptor subfamily 0 group B member 2 (NR0B2, also called SHP) (Figure S9). The strongest interactor is NFKB inhibitor beta (IκBβ), a direct repressor of the retinoid X receptor^30^ that has not been previously reported to bind LRH-1 (Figure S9). Both 9CA and 10CA disrupt many of the strongest coregulator interactions with apo-LRH-1 (*i.e*. IκBβ, NCOR1, and transcriptional cofactor of *c-fos* (TREF1)), while strongly recruiting a specific complement of coregulators (*e.g*. ligand dependent nuclear receptor corepressor (LCOR), nuclear receptor coactivator A1 (NCOA1) and nuclear receptor coactivator A3 (NCOA3), Figure 4a). In contrast, 4CA and 5CA have much weaker effects on coregulator associations (Figure 4a). These results suggest that the longer-tailed agonists promote conformational changes to the LBD that affect coregulator recognition of the AFS, consistent with observations from HDX-MS.

**Figure 4.**
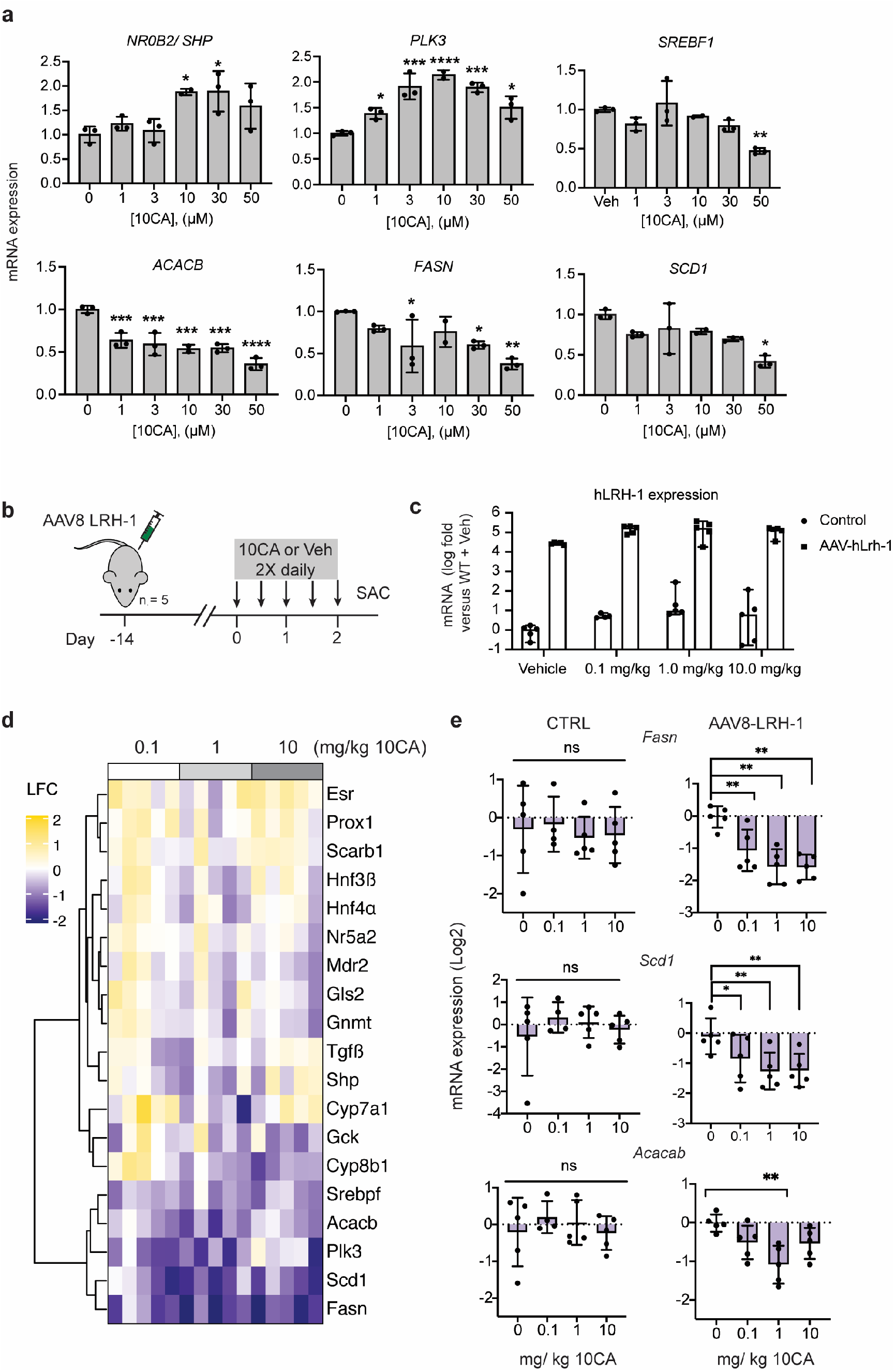
10CA activates LRH-1 in hepatocytes and in the liver. **a**, qRT-PCR analysis of LRH-1 targets in Huh7 hepatocytes. Cells were treated for 24 hours with 10CA concentrations indicated in the x-axes. Each bar represents the mean fold change *versus* DMSO-treated cells, with individual data points shown. Error bars are standard deviation for 2-3 replicates. Data are normalized to TBP. *, p < 0.05; **, p < 0.01; ***, p < 0.001; ****, p < 0.0001 by one-way ANOVA followed by Dunnett’s multiple comparisons test. **b**, Schematic of the experimental design for the mouse study. Mice were injected with AAV8-hLRH-1 intravenously to induce human LRH-1 expression, then after two weeks were treated with five doses of 10CA over three days. **c**, Liver tissue from mice was isolated and analyzed by qRT-PCR to measure levels of human LRH-1. **d**, Heatmap of Nanostring results in mice expressing AAV8-hLRH-1 and treated with vehicle, 0.1, 1.0, or 10.0 mg/ kg 10CA. The log_2_ fold changes (LFC) *versus* vehicle-treated mice are shown. **e**, Individual bar graphs from Nanostring showing downregulation of lipogenic genes in mice receiving AAV8-hLRH-1 but not in control (CTRL) mice. *, p < 0.05, **p, 0.01, by two-way ANOVA followed by Benjamini-Yekutieli False Discovery Rate (FDR) method, using an FDR threshold of 0.05.

In addition to demarking differences between active and inactive compounds, the coregulator recruitment pattern of 9CA and 10CA also differs from that of 6N and 6Na, LRH-1 agonists that share the 6HP scaffold but are modified elsewhere on the molecule (Figure S1).^24,25^ Unsupervised hierarchical clustering of the data shows a striking separation by ligand class: the most structurally similar ligands cluster together (Figure 4b). The close relationship between 9CA and 10CA in the clustering analysis appears to be due to common effects on coregulator dissociations. This distinguishes them from 4CA, which exhibits an otherwise similar (albeit weaker) pattern of coregulator associations to the longer-tailed compounds (Figure 4b). Coregulator recruitment by 6N and 6Na is also weak relative to the longer-tailed CAs, but it also displays a different pattern. For example, 9CA and 10CA promote dissociation of 16 coregulator peptides in common, while 6N and 6Na affect a distinct set of five coregulator peptides (Figure 4c).

Many of the observations from the experiment with isolated LBD are recapitulated in a MARCoNI study with FL-LRH-1. This is not a direct correlate to the LBD experiment, since apo-FL protein was unstable, and FL-LRH-1 in this experiment was bound to copurifying *E. coli* PL. Interestingly, recombinant FL-LRH-1 exhibits a nearly identical coregulator binding pattern to untreated apo-LBD (Figure S9). The FL protein also responds very similarly to 10CA, strongly recruiting peptides from LCOR, NCOA1, and SHP and strongly dissociating IκBβ, NCoR1, and TREF (Figure S9). A total of 85% of the peptides affected by at least 1.5-fold over baseline by 10CA in the LBD experiment are also altered by more than 1.5-fold with FL-LRH-1 (Figure 4d). As in the LBD experiment, the effects of 6N are relatively weak and mainly involve coregulator dissociation, including IκBβ and several canonical LRH-1 corepressors (*e.g*. NCOR1, PROX1, SHP, and DAX-1, Figure S9). Differences between 10CA and 6N are more apparent with FL-LRH-1 than LBD: 22 peptides are recruited by one compound and displaced by the other (Figure 4e). Differential coregulator recruitment by 10CA *versus* 6N are indicative of distinct ligand-induced conformational changes at the LRH-1 AFS, demonstrating that receptor allostery is tunable through ligand modifications. These studies also reveal the importance of the PL-mimicking group in driving coregulator recruitment.

### 10CA reduces lipogenic gene expression in the liver

LRH-1 is highly expressed in the liver, where it regulates bile acid biosynthesis, ER-stress resolution, and *de novo* lipogenesis.^9,10,31^ We tested whether 10CA, one of the most active of the new compounds in luciferase assays, could activate endogenous LRH-1 in Huh7 hepatocytes. A 24-hour treatment of 10CA dose-dependently upregulates known LRH-1 transcriptional targets SHP (*NR0B2*) and polo-like kinase 3 (*PLK3*). 10CA also downregulates targets involved in *de novo* lipogenesis, including sterol regulatory element binding transcription factor 1 (*SREBF1*, also called *SREBP1-c*), acetyl-CoA carboxylase 2 (*ACACB*), fatty acid synthase (*FASN*), and stearoyl-CoA desaturase-1 (*SCD1*) (Figure 5a). Interestingly, downregulation of these lipogenic genes in the liver is associated with LRH-1-dependent antidiabetic effects by DLPC.^9^

**Figure 5.**
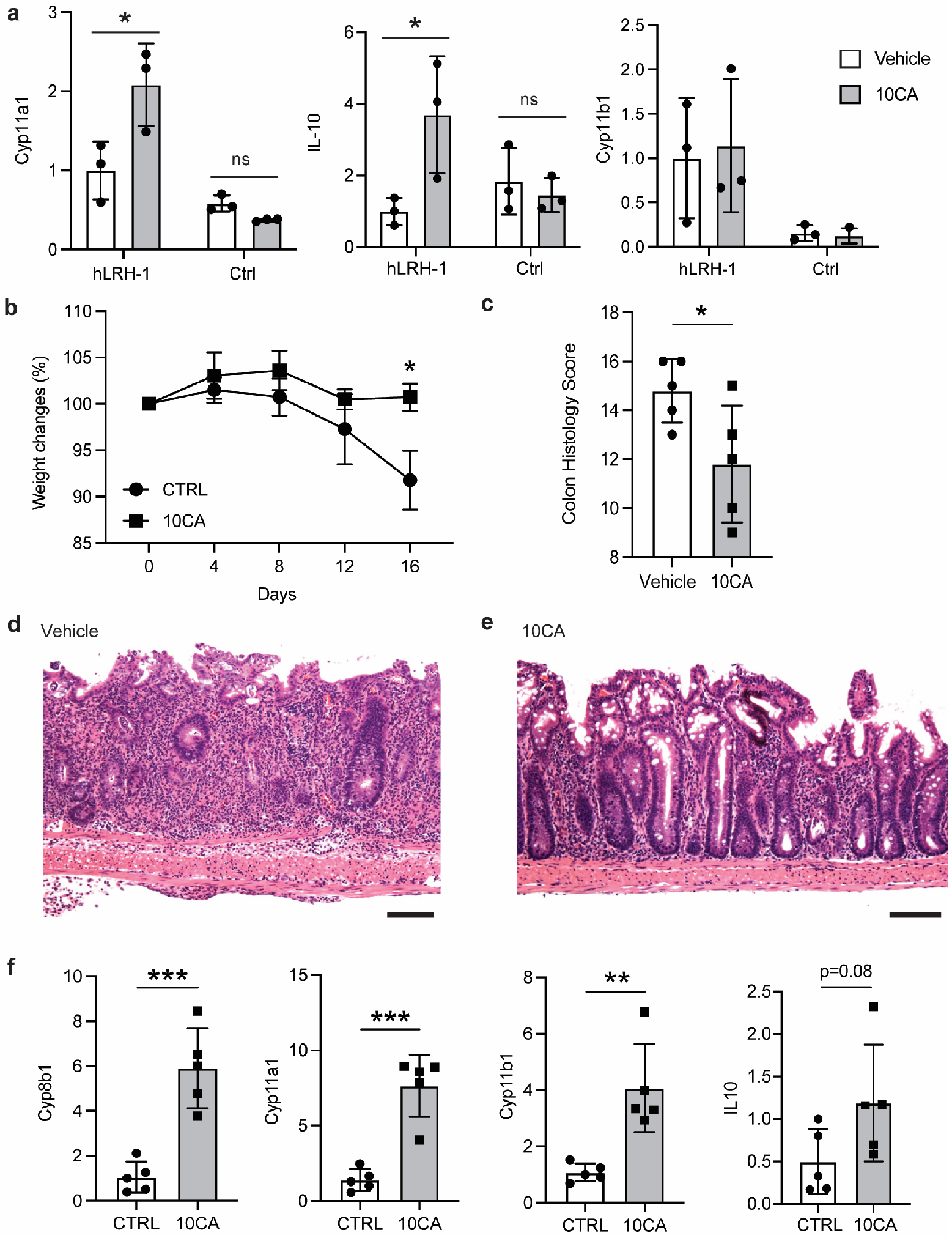
Efficacy of 10CA in models of intestinal inflammation and colitis. **a**, Gene expression changes in enteroids treated with 10CA, measured by qRT-PCR (n = 3). Bars represent the mean, error bars are standard deviation, and individual points are shown from three replicates. Bars labeled “hLRH-1” denote enteroids expressing human LRH-1 in animals where mouse Lrh-1 was conditionally deleted in the intestinal epithelium. Bars labeled “Ctrl” denote enteroids from epithelial-specific Lrh-1 knockout mice *, p < 0.05 by two-way ANOVA followed by Sidak’s multiple comparisons test. **b**, Plot of weight changes of mice over the course of treatment in TCT experiment. Mice (n = 5) were treated six days a week with vehicle (CTRL) or 10CA (20 mg/ kg). Each point represents the mean +/-SEM from five mice per group. *, p< 0.05 (two-way ANOVA followed by the Bonferroni test for multiple comparisons). **c**, Histology scores were calculated as described in the methods section and Table S3. Each bar represents the mean, error bars are SD, and individual values are shown as points. **d-e**, Representative colon sections stained with H&E show reduced intestinal damage with 10CA treatment (***e***) compared to vehicle (***d***) Scale bars are 100 nm. ***f***, qRT-PCR from of LRH-1 transcriptional targets in colonic crypts of mice from the TCT experiment. Each bar represents the mean, error bars are SD, and individual values are shown as points. *, p < 0.05, **, p < 0.01, ***, p < 0.001 relative to vehicle-treated organoids or mice by two-tailed, unpaired Student’s t-test.

To determine whether 10CA can activate LRH-1 in the liver *in vivo*, we overexpressed human LRH-1 (hLRH-1) in C57BL/6J mice using AAV8 adenovirus. The overexpression strategy was chosen because 10CA and analogs are poor activators of mouse LRH-1 in luciferase reporter assays. Two weeks after viral infection, mice were treated with 10CA at 0.1, 1, or 10 mg/kg in a series of five intraperitoneal injections spanning three days (Figure 5b). Expression of hLRH-1 was verified by qRT-PCR (Figure 5c), and agonist-induced gene expression changes in the liver were assessed by Nanostring.^32^ The most prominent effect of 10CA treatment is the downregulation of lipogenic genes (Figure 5d-e, Table S4). The affected genes are the same as those downregulated in Huh7 cells (*FASN, ACACB*, and *SCD1*). *SREBF1* also trends lower in 10CA-treated mice but does not reach statistical significance (p = 0.09 *versus* vehicle-treated controls, Table S4). Expression of hLRH-1 is required for downregulation of the lipogenic genes, since their expression is unaffected in control mice (given AAV8-GFP, Figure 5e).

### Efficacy of 10CA in organoid and murine models of colitis

To explore the efficacy of 10CA in a disease context, we tested it in models of intestinal inflammation and of ulcerative colitis (UC), a chronic disease characterized by inflammation of the colon and decreased integrity of the intestinal epithelium.^33^ LRH-1 is novel therapeutic target for UC: it regulates local glucocorticoid production in the intestine, which is important for maintenance of the intestinal epithelial barrier.^11,15,34,35^ Loss of LRH-1 function in mice decreases glucocorticoid production in the intestinal epithelium and decreases expression of *Cyp11a1* and *Cyp11b1*, genes that regulate corticosterone production.^11^ These defects are rescued by overexpression of hLRH-1.^15^

To evaluate the efficacy of 10CA *in vitro*, we used an organoids derived from intestinal crypts of mice in which hLRH-1 is expressed in the context of conditional deletion of endogenous LRH-1 in the intestinal epithelium. These intestinal organoids (enteroids) mimic intestinal inflammation when stimulated with tumor necrosis factor alpha.^15,36,37^ Overnight treatment with 10CA induces expression of *Cyp11a1* and the anti-inflammatory cytokine *Il-10* (Figure 5a). These changes in gene expression require the expression of hLRH-1, since they are not observed in epithelium-specific LRH-1-deficient enteroids. Expression of *Cyp11b1* was unchanged in these experiments (Figure 5a).

We used a naïve CD4^+^ T cell transfer (TCT) model of UC to test 10CA *in vivo*. In this model, immuno-compromised mice are implanted with naive T cells (CD4^+^CD45RB^hi^) to induce intestinal inflammation and destruction of the intestinal epithelium in a gradual manner that resembles chronic colitis^38^. Mice used in this experiment were previously generated by crossing recombinase activating gene-2-deficient (Rag2^-/-^) mice with mice expressing hLRH-1 but no endogenous Lrh-1 in the intestinal epithelium^38^. Three weeks after T cell transfer, mice (n = 5) were treated with 10CA (20 mg/ kg) or vehicle (CTRL, 0.09% normal saline) for 16 days via intraperitoneal injections. Vehicle-treated animals lost an average of 8.6% initial body weight over the course of treatment, while mice treated with 10CA were protected from weight loss and were within 1% of mean pre-treatment body weights at the end of the study (Figure 5b). 10CA-treated mice also had markedly improved colon histology scores, which quantify severity of inflammation, tissue damage, and crypt loss in the colon (Figure 5c). Representative colonic tissue sections illustrate the striking differences between 10CA-treated and control mice (Figure 5d-e). Colonic crypts isolated from 10CA-treated mice showed strong upregulation of LRH-1 transcriptional targets, including *Cyp8b1, Cyp11a1*, and *Cyp11b1* (Figure 5f). This is the first LRH-1 small molecule agonist to show *in vivo* efficacy in a model of UC, demonstrating the potential of PL-mimicking LRH-1 agonists for treatment of inflammatory bowel diseases.

## Discussion

The discovery that PCs act as NR ligands has illuminated the potential for novel signaling axes connecting PC metabolism with gene expression. Compelling evidence from gain- and loss-of-function studies connects PC availability to LRH-1-controlled transcriptional programs involving lipid and methyl metabolism. The response of LRH-1 to dietary PCs suggests that LRH-1 senses PCs as a proxy for nutrient availability, but this has been difficult to study. Likewise, PCs have anti-inflammatory effects in the intestinal mucosa that are not fully explained by their physical role in this tissue.^39–42^ The agonists described herein provide tools to probe a potential LRH-1-PC regulatory circuit, which is challenging to elucidate with natural PCs due to their lability and promiscuity.^43^

Crystallographic data reveals very different binding modes of natural PL and the synthetic LRH-1 modulator RJW100, with the former making polar interactions near the mouth of the pocket and the latter being anchored in the deep pocket.^7,23^ The hybrid PL-mimics were designed with the rationale that appending a polar moiety to an extended R^4^ alkyl tail would allow the agonists to interact with both regions of the pocket. With two new crystal structures, we demonstrate that this strategy was successful: the 6HP cores nearly perfectly superpose with RJW100, while the polar groups in the extended tails form hydrogen bonds with several residues engaged by the DLPC phosphate group (Figure 2). Disrupting these hydrogen bonds via mutagenesis reduces binding affinity and decreases activity of the longer-tailed agonists (Figure 2). Compounds with shorter tails or lacking a charged group at the R^4^ terminus are not more active than RJW100, further demonstrating that the PL-mimicking interactions are responsible for the improved binding and activity profiles (Figure 1). HDX-MS experiments show that the longer-tailed CAs stabilize the LRH-1 AFS relative to shorter-tailed CAs (Figure 3). The AFS stabilization is associated with much stronger effects on coregulator recruitment than seen with shorter-tailed CAs, including dissociation of many factors bound by apo-LRH-1 (Figure 3). Together, these data suggest a model in which specific interactions made by the PL-mimicking group promote conformational changes at the AFS that activate the receptor by modulating coregulator associations. Notably, the allostery observed for the PL-mimics is different from that of DLPC, which relies upon flexibility and motion of the region near helix 6.^16^ The PL-mimics do not displace helix 6 in the crystal structures (Figure S6b), and the flexibility of helix 6 is unchanged in HDX-MS experiments with 9CA and 10CA *versus* shorter-tailed CAs (Figure 3). The agonists therefore mimic PC interactions but act more as hybrids between PCs and synthetic agonists with respect to effects on receptor conformation and allostery. Nevertheless, the PL-mimics recapitulate many gene expression changes made by exogenous DLPC^9^, including downregulation of lipogenic genes in hepatocytes and mouse liver (Figure 4). Downregulation of the same lipogenic genes and consequent suppression of *de novo* lipogenesis is thought to be the main mechanism through which DLPC exerts anti-diabetic effects, including reducing liver fat accumulation and improving insulin resistance.^9^ This suggests a potential for the new agonists in diabetes and metabolic disorders associated with obesity, which is an ongoing area of research in our laboratories. A limitation of the mouse study with AAV8-hLRH-1 is that it is an overexpression model. Expression of the human receptor was needed due to amino acid differences in the binding pocket of the mouse homolog that reduce sensitivity to ligands^44^. However, hLRH-1 overexpression could increase basal activity, potentially blunting the effects of agonists. This is a possible reason that relatively few genes were altered by 10CA in the NanoString study. Future studies will address this limitation by utilizing a newly-generated humanized mouse.

LRH-1 is a novel therapeutic target for UC as an important regulator of intestinal cell renewal and local inflammation in the gut.^15,34^ One of the most exciting findings in these studies is the strong efficacy in UC *in vivo*. While gain and loss of function studies have demonstrated the potential of targeting LRH-1 in colitis,^15^ and the agonist 6N activates steroidogenic genes in enteroids,^24^ this is the first demonstration of a synthetic molecule to exhibit *in vivo* efficacy. In the TCT model of UC, LRH-1 activation by 10CA decreases disease markers, inhibits colonic crypt loss, and ameliorates disease-associated weight loss, with concomitant changes in expression of LRH-1 controlled steroidogenic genes (Figure 5). Targeting LRH-1 in UC is a particularly intriguing strategy, as its activation suppresses inflammation locally in the gut^34,45^. UC is commonly treated with corticosteroids, anti-TNFa therapies, or other systemic immunosuppressants that often cause adverse effects associated with immunosuppression in tissues other than the colon^46^. Thus, LRH-1 agonists may offer a more precise approach to UC treatment, reducing the potential for on-target, adverse effects in other tissues.

Together, this work has demonstrated an exciting potential for incorporating PL-mimicking interactions into LRH-1 agonists. The PL-mimics are valuable tools to study LRH-1 activation by ligands and to broaden our understanding of PL-regulated transcriptional programs. They also have potential as therapeutics for diseases associated with aberrant LRH-1 activity.

## Methods

### Statistics

Unless otherwise noted, all curve-fitting and statistics were conducted with GraphPad Prism software, v9.

### Chemical synthesis

Synthesis of the phosphorylcholine derivatives (4ChoP-11ChoP, also called 12a-12h) and carboxylic acid derivatives (4CA-11CA, also called 13a-13h) has been previously reported.^20^ Synthetic methods for RJW100 and analogs (compounds 1N-8N; 1X-8X; 9-23, used to generate Figure 2B) and compound 6Na have also been published previously.^25,36^ Synthesis of diols (4OH-11OH) are described in detail in the supplemental materials.

### Cell Culture

Hela cells were purchased from Atlantic Type Culture Collection (ATCC) and cultured in MEMα medium supplemented with 10% charcoal-stripped fetal bovine serum (FBS). Huh7 cells were provided by the Paul Dawson laboratory (Emory University) and cultured in DMEM/F12 medium supplemented with 10% FBS. Cell lines were authenticated by ATCC and tested negative for mycoplasma. Cells were maintained under standard culture conditions.

### Reporter gene assays

Hela cells were seeded into white-walled, 96-well culture plates at 7500 cells/ well. The next day, cells were transfected with reporter plasmids and either pCI empty vector or full-length LRH-1-pCI (5 ng/ well). Reporter plasmids consisted of (1) the pGL3 basic vector containing a portion of the SHP promoter containing the LRH-1 response element cloned upstream of firefly luciferase (50 ng/ well) and (2) a constitutively active vector encoding *Renilla* luciferase used for normalization (1 ng/ well). Transfections utilized the FugeneHD transfection reagent at a ratio of 5 μl Fugene: 2 μg DNA. Cells were treated with agonists 24 hours after transfection at concentrations indicated in the figure legends. Luminescence was quantified 24 hours after treatment using the DualGlo kit (Promega) on a BioTek Neo plate reader. EC_50_ and E_max_ were calculated by fitting data to a three-parameter dose-response curve.

### Calculation of Relative Efficacy (RE)

RE was calculated to compare data from luciferase reporter assays conducted on different dates by normalizing to data from RJW100 assessed in parallel experiments. We used the formula (Max_cpd_ – Min_cpd_) / (Max_Rjw100_ – Min_Rjw100_). “Cpd” denotes the test compound, and “Max” and “Min” denote the dose response curve maximum and minimum, respectively. A RE of 0 indicates a completely inactive compound, a value of 1 indicates equal activity to RJW100, and values above 1 indicate greater activity.

### Protein purification

#### LRH-1 LBD

BL21(pLysS) *E. coli* were transformed with LRH-1 LBD (residues 299-541) in the pMSC7 vector. Cultures were grown at 37 °C in liquid broth (LB) to an OD_600_ of 0.6 prior to induction of protein expression with 1 mM isopropyl-1-thio-D-galactopyranoside (IPTG). Cultures were grown for 4 hours at 30 °C following induction. Cells were harvested by centrifugation and resuspended into lysis buffer containing 20 mM Tris (pH 7.5), 150 mM NaCl, 5% glycerol, and 0.1 mM phenylmethylsulphoyl fluoride (PMSF). Protein was purified by nickel affinity chromatography with Buffer A of 20 mM Tris HCl (pH 7.4), 150 mM NaCl, 5% glycerol, and 25 mM imidazole and Buffer B of 150 mM Tris HCl (Buffer A containing 500 mM imidazole). Protein was incubated with DLPC (4-fold molar excess) overnight at 4 °C to displace copurifying *E. coli* PL. The protein was then re-purified by size exclusion into assay buffer (20 mM Tris HCl (pH 7.4), 150 mM NaCl, 5% glycerol), concentrated to ^~^3 mg/ ml, and stored at −80 °C. The same purification strategy was used for protein containing the point mutations described in the text.

#### Full-length LRH-1

Full-length human LRH-1 (FL-LRH-1; residues 1-495, isoform 1, codon optimized for *E. coli*) was cloned into a PET28b vector modified with a His-SUMO tag and transformed into BL21 (DE3) *E. coli* cells. Cells were grown in LB media at 37 °C to an OD_600_ of 0.5. Protein expression was induced with 0.2 mM IPTG and grown overnight at 16 °C. Cells were harvested by centrifugation and resuspended into lysis buffer containing 20 mM Tris (pH 7.5), 500 mM NaCl, 5% glycerol, 25 mM imidazole, 0.5 mM *tris*(2-carboxyethyl) phosphine (TCEP), and 0.1 mM PMSF. Cells were lysed by sonication for 8 minutes and clarified by centrifugation. Protein was purified by nickel affinity chromatography with elution buffer of 20 mM Tris HCl (pH 7.5), 500 mM NaCl, 5% glycerol, 0.5 mM TCEP and 125 mM imidazole. Ion-exchange chromatography was used to remove DNA with tandem Hitrap Q-SP columns (GE Healthcare). Protein was flowed through the Q column onto the SP column, from which it eluted as a single peak using a linear gradient of NaCl from 0.35 M to 1 M. Finally, the protein was purified to homogeneity by size exclusion chromatography. Functionality of the purified protein was determined by testing its ability to bind DNA containing an LRH-1 response element, and its ability to bind ligand. DNA binding was measured using fluorescence polarization with a 6-carboxyfluorescein (cFAM)-labeled double-stranded DNA probe containing the LRH-1 binding element from the CYP7A1 promoter. FL-LRH-1 was titrated against 10 nM FAM-CYP7A1 in buffer containing 20 mM Tris (pH 7.5), 150 mM NaCl, 5% glycerol and 0.5 mM TCEP, with a final volume of 50 μl, and incubated at room temperature for 30 min. Polarization was monitored on a Synergy 4 microplate reader (Biotek) at an excitation/emission wavelength of 485/528 nm. Curves were fit using a one-site specific binding equation with baseline-subtraction of wells containing protein but no probe. One biological replicate was conducted with two technical replicates. Ligand binding ability was verified by fluorescence polarization as described below.

### Generation of apo LRH-1

To extract lipids from LRH-1 LBD, 1 ml of purified protein (3 mg) was added to 3.75 ml of chloroform-methanol solution (1:2 v/v) and vortexed briefly. An additional 2.5 ml chloroform:water solution (1:1 v/v) was added and the mixture was vortexed again. The stripped and unfolded protein was pelleted by centrifugation at 1000 rpm for 10 minutes. The resulting protein pellet was dissolved into 0.5 ml of buffer containing 50 mM Tris (pH 8.0), 6 M guanidine hydrochloride and 2 mM DTT. Protein was refolded by fast dilution at 4 °C into 20 mM Tris (pH 8.5), 1.7 M urea, 4% glycerol and 2 mM DTT (25 ml). The final urea concentration was adjusted to 2 M, and the protein was concentrated to ^~^ 1.5 ml, followed by overnight dialysis against PBS containing 2 mM DTT at 4 °C. Refolded protein was purified by size exclusion chromatography to remove aggregates and remaining unfolded protein. Ligand binding ability was verified by fluorescence polarization as described below.

### Mutagenesis

Mutations were introduced to LRH-1 in the pMSC7 and pCI vectors using the Quikchange Lightning Site-Directed Mutagenesis Kit as directed by the manufacturer (Agilent). Constructs were sequenced prior to use.

### Ligand binding assay

Assays were conducted in black, polystyrene, non-binding surface 384-well plates (Corning) with 30 μl volumes in assay buffer (150 mM NaCl, 20 mM Tris-HCl pH 7.4, 5% glycerol). Plates were incubated overnight at 4 °C and centrifuged briefly before polarization measurement. Polarization was monitored on a Neo plate reader (Biotek) at excitation/emission wavelengths of 485/528 nm. We first measured the binding affinity of the fluorescein amidite (FAM)-labeled probe (6N-FAM) for WT and mutant LRH-1 by titrating the purified protein in the presence of 10 nM 6N-FAM, as previously described.^47^ Probe affinity and B_max_ for WT and mutant LRH-1 LBD was used to determine probe and protein concentrations for competition experiments (which were 10 nM 6N-FAM plus 5, 10, and 15 nM WT, K520A, or Y516A LRH-1, respectively). These assay conditions were validated using unlabeled 6N ligand, which was found to completely outcompete the probe for both mutant and WT LRH-1. Competition experiments used unlabeled agonists at a concentration range of 200 pM-200 μM. Experiments were conducted twice in quadruplicate, and compiled data was fit to a one-site, fit K_i_ curve. Statistical significance between K_i_ values was determined by two-way ANOVA followed by Dunnett’s test for multiple comparisons. For visual comparison of curves from WT or mutant protein (Figure S7), we normalized the data to competition curves generated from unlabeled 6N. To do this, FP data were baseline-subtracted, using the FP measurement from wells with the lowest concentration of competitor as the baseline. Then, we calculated the percent probe displacement at each concentration point relative to probe displacement by unlabeled 6N. This did not change the K_i_, but it put the curves on the same scale and provided a visual representation of the extent of probe displacement by each competitor.

### Crystallography

LRH-1 LBD was purified by nickel affinity chromatography as described above. The His tag was cleaved overnight at 4 °C using tobacco etch virus (TEV) protease with dialysis against buffer containing 150 mM NaCl, 5% glycerol, and 20 mM Tris HCl (pH 7.4). Cleaved protein was isolated using nickel affinity chromatography. To generate protein-ligand complexes, purified protein was incubated overnight at 4 °C with 4-5-fold molar excess of ligand. Complexes were purified by size exclusion chromatography into crystallization buffer consisting of 150 mM NaCl, 100 mM ammonium acetate (pH 7.4), 1 mM DTT, and 1 mM EDTA, 2 mM 3-[(3-cholamidopropyl)dimethylammonio]-1propanesulfonic acid (CHAPS) and then incubated with 4-fold molar excess of Tif2 peptide for two hours at room temperature (Tif2 peptide sequence was H_3_N-KENALLRYLLDK-CO_2_^-^). Crystallization conditions for each complex were as follows:

#### LRH-1-10CA-Tif2

The LRH-1-10CA-TIF2 complex was concentrated to 5 mg/mL and screened using the Classics screen (Qiagen) and a Phoenix Liquid Handler (Art Robbins Instruments) in 96-well sitting drop plates. Crystals were generated at room temperature in 0.4 M N/K tartrate.

#### LRH-1-9ChoP-Tif2

Crystals were generated by soaking 9ChoP into LRH-1-10CA-Tif2 crystals. First, we generated LRH-1-10CA-Tif2 crystals in larger drops by microseeding, using LRH-1-10CA-Tif2 crystals described above as seed stocks. Fresh LRH-1-10CA-Tif2 complex was concentrated to 5.2 mg/mL, and the seed stock was added to the protein at a dilution of 1:100. The complex was then crystallized by sitting drop vapor diffusion at 4 °C, in drops containing 1 μl protein/seed stock plus 2 μl crystallant. The crystallant was 9-19% tert-butanol, 0-6 % glycerol, 0.1 M tri-sodium citrate pH 4.6. The 9ChoP ligand was soaked into the crystal drops at 1 μM (^~^1% DMSO) for two days, and 1 μM 9ChoP was also added to the cryoprotectant.

### Structure Determination

Crystals were flash-frozen in liquid nitrogen using a cryoprotectant of 30% glycerol in crystallant. Data were collected remotely from Argonne National Laboratory (South East Regional Collaborative Access Team, 22ID beamline). Data were processed and scaled using HKL2000^48^ and phased by molecular replacement using Phaser-MR in Phenix^49^ with PDB 5L11 as the search model. For the 9ChoP structure, the 10CA structure was used as the search model. Coot^50^ and Phenix.refine^49^ were used for model building and refinement, respectively. Figures were constructed using Pymol.^51^ Measurements to determine displacement of helix 10 were between C-alphas of residue N523 for each protein.

### Hydrogen-deuterium exchange mass spectrometry (HDX-MS)

Following affinity purification and removal of the His-tag as described above, LRH-1 LBD protein was re-purified by size exclusion chromatography into an assay buffer of phosphate-buffered saline, pH 7.5, plus 5% glycerol. Protein aliquots (2 mg/ml) were incubated with carboxylic acid derivatives 4CA, 5CA, 9CA, or 10CA at 5-fold molar excess, overnight at 4 °C. Samples were diluted 1:7 (v/v) in labeling buffer to initialize exchange reactions (same as the protein buffer but containing 99.9 % D_2_O instead of H_2_O). The reactions were quenched after 0, 10, 100, 1000 and 10000 seconds by adding equal volume of precooled quenching buffer (100 mM phosphate, 0.5 M tris(2-carboxyethyl) phosphine, 0.8% formic acid, and 2% acetonitrile, pH 2.5). Reactions for each sample at each time point were performed in triplicate. Quenched samples were passed through an Enzymate BEH pepsin column (Waters). Fragmented peptides were separated by an C18 UPLC column and analyzed by a Q-Tof Premier mass spectrometer. ProteinLynx Global SERVER™ (PLGS) was used to identify peptides by database searching of LRH-1 LBD sequence. The HDX-MS data were further processed using DynamX (v3.0) and the differential HDX between ligand-bound states was calculated by comparing the relative fractional uptake for each residue at given time.

### NR-coregulator recruitment by MARCoNI

Assay mixes consisted of 25 nM ALEXA488-conjugated penta-His antibody (Qiagen # 35310), 50 μM DTT, 20mM Tris (pH 7.4), 250mM NaCl, 0.5 mM TCEP and purified protein: either (1) 50 nM His-LRH-1 LBD (in the apo state or pre-loaded with compounds), or (2) 50 nM His-SUMO-hLRH-1-FL (bound to copurifying PL from *E. coli* or preloaded with compound during purification). Compounds (dissolved in DMSO) were added to the assay mixes at final concentrations of 10 μM. DMSO was 2% v/v in all mixes. Coregulator profiling was conducted by Microarray Assay for Real-time Coregulator Nuclear Receptor Interaction (MARCoNI), using PamChip #88101.^52^ LRH-1-coregulator binding was measured in triplicate and quantified using BioNavigator software (PamGene). The compound-induced log-fold change (LFC) of LRH-1 binding to each coregulator peptide and statistical significance (FDR threshold of 0.05 and Student’s t-test) were calculated and visualized using R software (R Core Team, 2017). Compound and interaction (dis)similarity were calculated by Hierarchical Clustering on Euclidean Distance and Ward’s agglomeration. Upset plots were generated using the R package UpSetR v1.4.0.^53^

### Molecular dynamics simulations

Two crystal structures of LRH-1 LBD in complex with Tif2 were used for model construction (1) PDB 7JYD (10CA ligand) and (2) PDB 4DOS (DLPC). Structures were modified at the N and C termini so that they included residues 300-540 of LRH-1 and residues 742-751 of the Tif2 peptide. Missing residues and sidechains within this protein sequence were added. All complexes were solvated in an octahedral box of TIP3P water with a 10 Å buffer around the complex. Na^+^ and Cl^-^ ions were added to neutralize the complex and achieve physiological conditions (150 mM sodium chloride). All systems were generated with the Xleap module of AmberTools^54^, using the ff14SB force field.^55^ Parameters for all ligands were obtained using the Antechamber module^56^ of AmberTools. All minimizations and simulations were performed with Amber17^54^. Using a minimization protocol of 5000 steps of steepest descent followed by 5000 steps of conjugate gradient, systems were minimized in an iterative manner: i) with 500 kcal/mol.Å^2^ restraints on all atoms; ii) with 500 kcal/ mol.Å^2^ restraints on restraints retained on ligand and peptide atoms; iii) with restraints removed from all atoms. Following minimization, systems were heated from 0 to 300 K using a 100-ps run with constant volume periodic boundaries and 5 kcal/mol.Å^2^ restraints on all protein and ligand atoms. Equilibration was performed using a 12-ns MD run with 10 kcal/mol.Å^2^ restraints on protein and ligand atoms using the NPT ensemble. Restraints were reduced to 1 kcal/mol.Å^2^ for an additional 10 ns of equilibration. All restraints were removed, and production runs were performed for 1 μs in the NPT ensemble. A 2-fs time step was used for all simulations. All bonds between heavy atoms and hydrogens were fixed with the SHAKE algorithm.^57^ A 10-Å cutoff was used for nonbonded interactions. Long range electrostatics were evaluated using Particle Mesh Ewald (PME). For analysis, frames were obtained at 20 ps intervals from each simulation. The CPPTRAJ module^58^ of AmberTools was used for structural averaging of MD trajectories and distance calculations. The stability of hydrogen bonds between ligands and protein was calculated by measuring the distances between heavy atoms of interacting residues (for example, C of carboxylate groups, N of amino groups, O of hydroxyl groups) over the course of the simulations and quantifying the percentage of time they were within 3.5 Å of each other.

### Agonist treatment of Huh7 cells

Huh7 cells were plated in 6-well plates at 5×10^5^ cells/well. After 24 hours, cells were treated with vehicle (0.1% DMSO in media) or indicated concentrations of 10CA. Cells were collected for RNA extraction after 24 hours of treatment. RNA extraction was performed using the RNAeasy Mini RNA extraction kit (Qiagen).

### Animals

Experimental protocols for the AAV8-LRH-1 overexpression study were approved by the Institute for Animal Care and Use Committee at T3 Laboratories and Emory University and conformed to the Guide for the Care and Use of Laboratory Animals, published by the National Institutes of Health (NIH Publication No. 86-23, revised 1996), and with federal and state regulations. C57BL/6J mice (male; 10-12 weeks of age) were purchased from Jackson Labs (Bar Harbor, ME, USA). Mice were housed at room temperature in a 12-hr light/12-hr dark vivarium, with *ad libitum* access to food and water.

For the experiments involving enteroids and TCT, the study protocol was approved by the Animal Care and Use Committee of the Baylor College of Medicine and was in accordance with the Guide for the Care and Use of Laboratory Animals [DHHS publication no. (NIH) 85-23, revised 1985, Office of Science and Health Reports, DRR/NIH, Bethesda, MD 20205]. Animals were housed and bred in a specific pathogen free (SPF) facility. An inducible intestinal epithelial cell (IEC) knockout line was created by crossing animals harboring CreERT2 under control of the villin promoter with LRH-1 floxed (LRH-1^f/f^) animals and bred to homozygosity (LRH-1 KO, LRH-1^f/f^;Villin-Cre+). The humanized LRH-1 line (LRH-1^f/f^;hLRH-1^ΔΔ^;Villin-Cre+) was generated with Rosa26-LoxP-STOP-LoxP-hLRH-1 animals, crossed into our LRH-1 KO line, which conditionally overexpresses hLRH-1 in the intestinal epithelium with the knockout of endogenous mLRH-1.

### Viral overexpression and drug treatment

Adeno-associated virus serotype 8 (AAV8) expressing N-terminal Avi-tagged human LRH-1 and AAV8-GFP (used as a negative control) were obtained from Vector Biolabs (Malvern, PA, USA). C57BL/6J mice were administered 1×10^12^ genome copies (GC)/mL via femoral vein injection. Two weeks after virus treatment, mice (n = 5) were administered 0.1, 1.0, or 10 mg/ kg of 10CA or vehicle (5% ethanol in corn oil) via intraperitoneal injection five times over a 3-day period. Mice were treated each morning and evening for two days and on the morning of day 3. Mice were euthanized on day 3, approximately four hours after the final drug treatment. Mouse liver tissue was collected, flash-frozen in liquid nitrogen, and stored at −80 °C. RNA was extracted from the liver samples using the miRNAeasy Mini kit (Qiagen), following cryo-pulverization of the tissues in liquid nitrogen and homogenization in Trizol (Invitrogen). A portion of the RNA was retained for NanoString analysis, and a portion was subjected to qRT-PCR to verify overexpression of hLRH-1. Reverse transcription was performed using the High Capacity cDNA Reverse Transcription Kit (Applied Biosystems). qRT-PCR used the *Power* SYBR^TM^ Green PCR Master Mix (Applied Biosystems) and run on a StepOnePlus™ Real-Time PCR instrument (Applied Biosystems). Data were analyzed via the ΔΔCt method, with *Tata binding protein* (*Tbp*) as the reference gene. Primer sequences are shown in Table S2.

### NanoString Gene Expression Analysis

RNA concentrations were measured on a Nanodrop 1000, and quality was assessed on a 2100 Bioanalyzer Instrument (Agilent) using the RNA 6000 Nano (25ng/ul – 1,000ng/ul) or Pico assay (0.5ng/ul – 20ng/ul). All RNA Integrity (RIN) scores were >9.0.25ng of total RNA was hybridized with biotin-labeled capture probes and fluorescently-labeled reporter probes (NanoString Technologies, Inc.) for 18 hours at 65 °C. Following hybridization, samples were injected into a NanoString SPRINT cartridge and loaded onto the nCounter SPRINT Profiler instrument, where hybridized mRNAs were immobilized for imaging and excess probes were removed. Following image acquisition, mRNA counts were extracted from raw RCC files using nSolver™ data analysis software (v4.0, NanoString Technologies, Inc.). Significance was assessed by two-way ANOVA, followed by Benjamini-Yekutieli False Discovery Rate (FDR) method, using an FDR threshold of 0.05, as recommended by Nanostring gene expression guidelines. Hierarchical clustering of the data on Euclidean distance and Ward’s agglomeration was done using the R package ComplexHeatmap.

### Humanized LRH-1 Mouse Intestinal Enteroid Culture

Intestinal crypt cultures (enteroids) were derived from LRH-1 KO (LRH-1^f/f^;Villin-Cre+), and hLRH-1 (LRH-1^f/f^;hLRH-1^ΔΔ^;Villin-Cre+) male mice (6–8 weeks old)^15^. Briefly, the small intestine was isolated and flushed with ice-cold PBS, opened longitudinally, then cut into 1–2 mm pieces. Intestinal fragments were incubated in an EDTA-containing solution (4 mM) at 4 °C for 60 min on a tube rocker. The intestinal fragment suspension was fractionated by vertical shaking manually and crypt-containing fractions passed through a 70 μm cell strainer for plating in Matrigel. The crypt-Matrigel suspension was allowed to polymerize at 37 °C for 15 min. Intestinal organoids were grown in base culture media (Advanced DMEM/F12 media, HEPES, GlutaMax, penicillin, and streptomycin) supplemented with growth factors (EGF, Noggin, R-spondin, R&D Systems), B27 (Life Technologies), N2 (Life Technologies), and N-acetyl cysteine (Sigma). Intestinal enteroids were passaged every 3 days. Established hLRH-1 or LRH-1 KO enteroids were treated with mouse TNF-α overnight to provoke inflammatory changes, then treated with vehicle (DMSO) or compound 10CA (1 μM) overnight. Following the treatment, enteroid tissues were harvested for real-time PCR.

### RNA Isolation and qRT-PCR of enteroids

Enteroids were washed in ice cold PBS and suspended in Trizol solution (Sigma). RNA was isolated with RNeasy spin columns (Qiagen). DNAse-treated total RNA was used to generate cDNA using Superscript II (Quanta). SYBR green-based qPCR (Kapa Biosystems) was performed on a Roche LightCycler 480 II. The ΔΔC_t_ method was used for calculating gene expression fold changes using Rplp0 (ribosomal protein, large, P0, known as 36B4) as the reference. Primer sequences are shown in Table S2.

### T Cell transfer model of colitis

Animals were housed and bred in SPF facility at Baylor College of Medicine. The *hLRH1*^IEC^ (*Rag2^-/-^ hLRH1^IEC-TG^* mLRH*1*^IEC-KO^) mouse line was generated by crossing *Flox-Stop-Flox hLRH-1;VilCre* animals with *Rag2*^-/-^ *mLRH1^flox/flox^*;*VilCre* animals *(hLRH1^IEC^)* animals, in which human LRH-1 replaces mouse LRH-1 in enterocytes. For T cell transfer experiments, *hLRH1^IEC^* mice were used to induce the chronic enterocolitis as described^38^. Briefly, wild-type splenic CD4^+^CD45RB^high^ cells were isolated by MACS separation and flow cytometry cell sorting, and then transferred by intraperitoneal injection to recipient mice (0.5 × 10^6^ cells/mice). All recipients were weighed immediately prior to T cell transfer to determine baseline weight, and then weighed twice weekly after T cell transfers for the duration of the experiment. 10CA treatment started from week 4 after the T cell transfer, 20 mg/kg 10CA in sterile saline was injected intraperitoneally (i.p.) 6 days per week. Normal saline was used as control treatment. Transferred recipient mice were euthanized upon losing 20% of pre-transfer baseline weight. All recipient mice receiving different treatment were co-housed throughout to normalize microflora exposure.

At the end of treatment, disease activity scores were assessed according to Table S3. For colon histological analysis, the colon was divided into three segments (proximal, middle, and distal). Each segment was embedded in paraffin, sectioned at 5 μm, and stained with hematoxylin and eosin. Histological analysis was performed in the Cellular and Molecular Morphology Core of the Digestive Disease Center at Baylor College of Medicine. The sections were blindly scored using a standard histologic colitis score. Three independent parameters were measured: severity of inflammation (0–3: none, slight, moderate, severe), depth of injury (0–3: none, mucosal, mucosal and submucosal, transmural), and crypt damage (0–4: none, basal one-third damaged, basal two-thirds damaged, only surface epithelium intact, entire crypt and epithelium lost). The score of each parameter was multiplied by a factor reflecting the percentage of tissue involvement (×1, 0–25%; ×2, 26–50%; ×3, 51–75%; ×4, 76–100%) averaged per colon.

## Acknowledgements

The authors thank the HDX-MS core in School of Medicine, Emory University for their technical assistance in data collection and analysis. This study was supported in part by the Emory Integrated Genomics Core (EIGC), which is subsidized by the Emory University School of Medicine and is one of the Emory Integrated Core Facilities. The authors are grateful to the beamline staff at Argonne National Laboratory, South East Regional Collaborative Team, for support during remote collection of crystal diffraction data. This work was supported in part by the National Institutes of Health under the following awards: T32GM008602 (S.G.M.), F31DK111171 (S.G.M.), T32GM008367 (E.H.D. and M.L.C.), R01DK095750 (E.A.O.), R01DK114213 (E.A.O., N.T.J., J.W.C). The work was also supported by an Emory Catalyst Award (E.A.O., N.T.J), and USDA ARS 3092-5-001-057 (D.D.M.). E.H.D was supported by the National Science Foundation Graduate Research Fellowship.

## Author Contributions

S.G.M., E.H.D., A.R.F., E.J.M., C.D.O., A.P., R.H., D.D.M., J.W.C., N.T.J., and E.A.O. participated in research design. S.G.M., E.H.D., A.R.F., X.H., G.W., X.L., E.J.M., C.D.O., A.P., M.L.C., J.L.C., and D.M. conducted experiments. S.G.M., E.H.D., A.R.F., X.H., G.W., X.L., E.J.M., C.D.O., A.P., M.L.C., J.L.C., D.M., and R.H. performed data analysis. S.G.M., E.H.D., and E.A.O. wrote the manuscript, with contributions from other authors.

## Competing Interests Statement

The authors declare no competing financial interests.

## Data availability

Authors will release the atomic coordinates and experimental data upon article publication. Data for macromolecular crystal structures are available in the Protein Data Bank (PDB). PDB IDs have been provided in figure legends and are as follows: LRH-1-10CA, 7JYD; LRH-1-9ChoP, 7JYE.

## Code availability

The R scripts used to analyze MARCoNI and Nanostring experiments are available upon request.

## Notes

### Competing Interest Statement

The authors have declared no competing interest.

### Summary of Updates

We have included new data showing ligand-drive LRH-1 activity in a T-cell model of Colitis. This replaces the prior DSS study.

## References

1. Liu, S. et al. A diurnal serum lipid integrates hepatic lipogenesis and peripheral fatty acid use. Nature 502, 550–554 (2013).

2. Wymann, M.P. & Scheiter, R. Lipid signalling in disease. Nat Rev Mol Cell Biol 9, 162–176 (2008).

3. Li, Z. et al. The ratio of phosphatidylcholine to phosphatidylethanolamine influences membrane integrity and steatohepatitis. Cell Metab. 3, 321–331 (2006).

4. Furse, S. & de Kroon, A.I. Phosphatidylcholine’s functions beyond that of a membrane brick. Mol Membr Biol 32, 117–119 (2015).

5. Walker, A.K. 1-Carbon Cycle Metabolites Methylate Their Way to Fatty Liver. Trends Endocrinol Metab 28, 63–72 (2017).

6. Musille, P.M., Kohn, J.A. & Ortlund, E.A. Phospholipid - Driven gene regulation. FEBS Letters 587, 1238–1246 (2013).

7. Musille, P.M. et al. Antidiabetic phospholipid-nuclear receptor complex reveals the mechanism for phospholipid-driven gene regulation. Nat. Struct. Mol. Biol. 19, 532–537, S531-532 (2012).

8. Chakravarthy, M.V. et al. Identification of a physiologically relevant endogenous ligand for PPARalpha in liver. Cell 138, 476–488 (2009).

9. Lee, J.M. et al. A nuclear-receptor-dependent phosphatidylcholine pathway with antidiabetic effects. Nature 474, 506–510 (2011).

10. Mamrosh, J.L. et al. Nuclear receptor LRH-1/NR5A2 is required and targetable for liver endoplasmic reticulum stress resolution. Elife 3, e01694 (2014).

11. Coste, A. et al. LRH-1-mediated glucocorticoid synthesis in enterocytes protects against inflammatory bowel disease. Proc. Natl. Acad. Sci. U. S. A. 104, 13098–13103 (2007).

12. Xu, P. et al. LRH-1-dependent programming of mitochondrial glutamine processing drives liver cancer. Genes Dev 30, 1255–1260 (2016).

13. Nadolny, C. & Dong, X. Liver receptor homolog-1 (LRH-1): a potential therapeutic target for cancer. Cancer Biol. Ther. 16, 997–1004 (2015).

14. Cobo-Vuilleumier, N. et al. LRH-1 agonism favours an immune-islet dialogue which protects against diabetes mellitus. Nat. Commun. 9, 1488 (2018).

15. Bayrer, J.R. et al. LRH-1 mitigates intestinal inflammatory disease by maintaining epithelial homeostasis and cell survival. Nat. Commun. 9, 4055 (2018).

16. Musille, P.M., Kossmann, B.R., Kohn, J.A., Ivanov, I. & Ortlund, E.A. Unexpected Allosteric Network Contributes to LRH-1 Coregulator Selectivity. J. Biol. Chem. 291, 1411–1426 (2015).

17. Wagner, M. et al. Liver receptor homolog-1 is a critical determinant of methyl-pool metabolism. Hepatology 63, 95–106 (2016).

18. Lin, C.J. & Wang, M.C. Microbial metabolites regulate host lipid metabolism through NR5A-Hedgehog signalling. Nat Cell Biol 19, 550–557 (2017).

19. Choi, S. et al. Methyl-sensing nuclear receptor Liver Receptor Homolog-1 regulates mitochondrial function in mouse hepatocytes. Hepatology 71, 1055–1069 (2019).

20. Flynn, A.R., Mays, S.G., Ortlund, E.A. & Jui, N.T. Development of Hybrid Phospholipid Mimics as Effective Agonists for Liver Receptor Homologue-1. ACS Med. Chem. Lett. 9, 1051–1056 (2018).

21. Ortlund, E.A. et al. Modulation of human nuclear receptor LRH-1 activity by phospholipids and SHP. Nat. Struct. Mol. Biol. 12, 357–363 (2005).

22. Sablin, E.P. et al. Structure of Liver Receptor Homolog-1 (NR5A2) with PIP hormone bound in the ligand binding pocket. J. Struct. Biol. 192, 342–348 (2015).

23. Mays, S.G. et al. Crystal Structures of the Nuclear Receptor, Liver Receptor Homolog 1, Bound to Synthetic Agonists. J. Biol. Chem. 291, 25281–25291 (2016).

24. Mays, S.G. et al. Development of the first low nanomolar liver receptor homolog-1 agonist through structure-guided design. J. Med. Chem. 62, 11022–11034 (2019).

25. Cornelison, J.L. et al. Development of a new class of liver receptor homolog-1 (LRH-1) agonists by photoredox conjugate addition. Bioorganic & Medicinal Chemistry Letters 30 (2020).

26. Musille, P.M. et al. Antidiabetic phospholipid-nuclear receptor complex reveals the mechanism for phospholipid-driven gene regulation. Nat Struct Mol Biol 19, 532–537, S531-532 (2012).

27. Koppen, A. et al. Nuclear Receptor-Coregulator Interaction Profiling Identifies TRIP3 as a Novel Peroxisome Proliferator-activated Receptor. Molecular and Cellular Proteomics, 2211–2226 (2009).

28. Ortlund, E.A. et al. Modulation of human nuclear receptor LRH-1 activity by phospholipids and SHP. Nat Struct Mol Biol 12, 357–363 (2005).

29. Musille, P.M., Kossmann, B.R., Kohn, J.A., Ivanov, I. & Ortlund, E.A. Unexpected Allosteric Network Contributes to LRH-1 Coregulator Selectivity. J Biol Chem (2015).

30. Na, S.-Y. et al. IkBb Interacts with the Retinoid X Receptor and Inhibits Retinoid-dependent Transactivation in Lipopolysaccharide-treated Cells. J. Biol. Chem. 273, 3212–3215 (1998).

31. Lu, T. et al. Molecular Basis for Feedback Regulation of Bile Acid Synthesis by Nuclear Receptors. Mol. Cell 6, 507–515 (2000).

32. Geiss, G.K. et al. Direct multiplexed measurement of gene expression with color-coded probe pairs. Nature Biotechnology 26, 317–325 (2008).

33. Ordás, I., Eckmann, L., Talamini, M., Baumgart, D.C. & Sandborn, W.J. Ulcerative colitis. The Lancet 380, 1606–1619 (2012).

34. Mueller, M. et al. The nuclear receptor LRH-1 critically regulates extra-adrenal glucocorticoid synthesis in the intestine. J. Exp. Med. 203, 2057–2062 (2006).

35. Bouguen, G. et al. Intestinal steroidogenesis controls PPARgamma expression in the colon and is impaired during ulcerative colitis. Gut 64, 901–910 (2015).

36. Mays, S.G. et al. Development of the First Low Nanomolar Liver Receptor Homolog-1 Agonist through Structure-guided Design. J Med Chem (2019).

37. Basak, O. et al. Induced Quiescence of Lgr5+ Stem Cells in Intestinal Organoids Enables Differentiation of Hormone-Producing Enteroendocrine Cells. Cell Stem Cell 20, 177–190 e174 (2017).

38. Ostanin, D.V. et al. T cell transfer model of chronic colitis: concepts, considerations, and tricks of the trade. Am J Physiol Gastrointest Liver Physiol 296, G135–146 (2009).

39. Treede, I. et al. Anti-inflammatory effects of phosphatidylcholine. J. Biol. Chem. 282, 27155–27164 (2007).

40. Olson, A., Diebel, L.N. & Liberati, D.M. Exogenous phosphatidylcholine supplementation improves intestinal barrier defense against Clostridium difficile toxin. J Trauma Acute Care Surg 77, 570–575; discussion 576 (2014).

41. Chen, M. et al. Phosphatidylcholine regulates NF-κB activation in attenuation of LPS-induced inflammation: evidence from in vitro study. Animal Cells and Systems 22, 7–14 (2017).

42. Schneider, H., Braun, A., Fullekrug, J., Stremmel, W. & Ehehalt, R. Lipid based therapy for ulcerative colitis-modulation of intestinal mucus membrane phospholipids as a tool to influence inflammation. Int J Mol Sci 11, 4149–4164 (2010).

43. MacDonald, J.I., Specher, H. Phospholipid Fatty Acid Remodeling in Mammalian Cells. Biochim Biophys Acta, Lipids Lipid Metab 1084, 105–121 (1991).

44. Musille, P.M., Pathak, M., Lauer, J.L., Griffin, P.R. & Ortlund, E.A. Divergent sequence tunes ligand sensitivity in phospholipid-regulated hormone receptors. J. Biol. Chem. 288, 20702–20712 (2013).

45. Fernandez-Marcos, P.J., Auwerx, J. & Schoonjans, K. Emerging actions of the nuclear receptor LRH-1 in the gut. Biochim Biophys Acta 1812, 947–955 (2011).

46. Gensler, L.S. Glucocorticoids: complications to anticipate and prevent. Neurohospitalist 3, 92–97 (2013).

47. D’Agostino, E.H. et al. Development of a Versatile and Sensitive Direct Ligand Binding Assay for Human NR5A Nuclear Receptors. ACS Med. Chem. Lett. (2019).

48. Otwinowski, Z., Minor, W. Processing of X-Ray Diffraction Data Collected in Oscillation Mode. Method. Enzymol. 276, 307–326 (1997).

49. Adams, P.D. et al. PHENIX: a comprehensive Python-based system for macromolecular structure solution. Acta. Crystallogr. D Biol. Crystallogr. 66, 213–221 (2010).

50. Emsley, P. & Cowtan, K. Coot: model-building tools for molecular graphics. Acta. Crystallogr. D Biol. Crystallogr. 60, 2126–2132 (2004).

51. Schrodinger, LLC. The PyMOL Molecular Graphics System, Version 1.3r1. (2010).

52. Aarts, J.M. et al. Robust array-based coregulator binding assay predicting ERalpha-agonist potency and generating binding profiles reflecting ligand structure. Chem Res Toxicol 26, 336–346 (2013).

53. Conway, J.R., Lex, A. & Gehlenborg, N. UpSetR: an R package for the visualization of intersecting sets and their properties. Bioinformatics 33, 2938–2940 (2017).

54. Case, D. et al. AMBER 2017. University of Californa, San Franscisco (2017).

55. Maier, J.A.M., C.; Kasavajhala, K.; Wickstrom, L.; Hauser, K. E.; Simmerling, C. ff14SB: Improving the accuracy of protein side chain and backbone parameters from ff99SB. Journal of chemical theory and computation 11, 3696–3713 (2015).

56. Wang, J.W., W.; Kollman, P. A.; Case, D. A. Antechamber: an accessory software package for molecular mechanical calculations. J Am Chem Soc 222, U403 (2001).

57. Ryckaert, J.-P.C., G.; Berendsen, H. J. Numerical integration of the cartesian equations of motion of a system with constraints: molecular dynamics of n-alkanes. Journal of Computational Physics 23, 327–341 (1977).

58. Roe, D.R. & Cheatham III, T.E. PTRAJ and CPPTRAJ: software for processing and analysis of molecular dynamics trajectory data. Journal of chemical theory and computation 9, 3084–3095 (2013).

