## Supplemental Information for "A novel phospholipid mimetic targeting LRH-1 ameliorates colitis"

Nathan T. Jui

Department of Chemistry

Emory University

1515 Dickey Drive, 456

Atlanta, GA 30322

Running title: Phospholipid mimetics as LRH-1 agonists

### Table of Contents

*I. Chemical synthesis and characterization* ... *3*

**General Synthetic Information** ... **3**

**Synthesis of PC Mimics 1–3** ... **4**

**Hybrid Precursors: Compounds S1a–S1h** ... **5**

Phosphorylcholines: Compounds 1a–h ... 8

Carboxylate Precursors: Compounds S2a–S2h ... 10

Carboxylates: Compounds 2a–2h ... 13

Diols: Compounds 3a–3h ... 16

*II. Supplemental Figures S1-S9* ... *19*

**Figure S1. Chemical structures of compounds 6N and 6Na** ... **19**

**Figure S2. Fluorescence polarization competition binding curves: phosphorylcholines.** ... **20**

**Figure S3. Fluorescence polarization competition binding curves: carboxylic acids** ... **21**

**Figure S4. Fluorescence polarization competition binding curves: diols** ... **22**

**Figure S5. Luciferase reporter assays with diols.** ... **23**

**Figure S6. Structural analyses.** ... **24**

**Figure S7. Fluorescence polarization competition binding curves: LRH-1 mutants.** ... **25**

**Figure S8. Peptide coverage and deuterium uptake in HDX-MS experiments** ... **27**

**Figure S9. Protein purification and MARCoNI.** ... **29**

*III. Supplemental Tables S1-S4* ... *29*

**Table S1. Summary of linker lengths and key biological parameters for LRH-1 agonists.** ... **29**

**Table S2. Sequences of primers used for qRT-PCR.** ... **31**

**Table S3. Modified colitis disease activity score.** ... **31**

**Table S4. Gene expression changes in the livers of mice expressing hLRH-1.** ... **32**

### I. Chemical synthesis and characterization

#### General Synthetic Information

All reactions were carried out in oven-dried glassware, equipped with a stir bar and under a nitrogen atmosphere with dry solvents under anhydrous conditions, unless otherwise noted. Solvents used in anhydrous reactions were purified by passing over activated alumina and storing under argon. Yields refer to chromatographically and spectroscopically ($^{1}H$ NMR) homogenous materials, unless otherwise stated. Reagents were purchased at the highest commercial quality and used without further purification, unless otherwise stated. n-Butyllithium (n-BuLi) was used as a 1.6 M or a 2.5 M solution in hexanes (Aldrich), was stored at 4°C and titrated prior to use. Organic solutions were concentrated under reduced pressure on a rotary evaporator using a water bath. Chromatographic purification of products was accomplished using forced-flow chromatography on 230-400 mesh silica gel. Preparative thin-layer chromatography (PTLC) separations were carried out on 1000µm SiliCycle silica gel F-254 plates. Thin-layer chromatography (TLC) was performed on 250µm SiliCycle silica gel F-254 plates. Visualization of the developed chromatogram was performed by fluorescence quenching or by staining using $KMnO_4$, p-anisaldehyde, or ninhydrin stains.

$^{1}H$ and $^{13}C$ NMR spectra were obtained from the Emory University NMR facility and recorded on a Bruker Avance III HD 600 equipped with cryo-probe (600 MHz), INOVA 600 (600 MHz), INOVA 500 (500 MHz), INOVA 400 (400 MHz), VNMR 400 (400 MHz), or Mercury 300 (300 MHz), and are internally referenced to residual protio solvent signals. Data for $^{1}H$ NMR are reported as follows: chemical shift (ppm), multiplicity (s = singlet, d = doublet, t = triplet, q = quartet, m = multiplet, dd = doublet of doublets, dt = doublet of triplets, ddd= doublet of doublet of doublets, dtd= doublet of triplet of doublets, b = broad, etc.), coupling constant (Hz), integration, and assignment, when applicable. Data for decoupled $^{13}C$ NMR are reported in terms of chemical shift and multiplicity when applicable. IR spectra were recorded on a Thermo Fisher Diamond-ATR and reported in terms of frequency of absorption ($cm^{-1}$). High Resolution mass spectra were obtained from the Emory University Mass Spectral facility. Gas Chromatography Mass Spectrometry (GC-MS) was performed on an Agilent 5977A mass spectrometer with an Agilent 7890A gas chromatography inlet. Liquid Chromatography Mass Spectrometry (LC-MS) was performed on an Agilent 6120 mass spectrometer with an Agilent 1220 Infinity liquid chromatography inlet. Preparative High Pressure Liquid chromatography (Prep-HPLC) was performed on an Agilent 1200 Infinity Series chromatograph using an Agilent Prep-C18 30 x 250 mm 10 µm column, or an Agilent Prep-C18 21.2 x 100 mm, 5 µm column.

#### Synthesis of PC Mimics 1–3

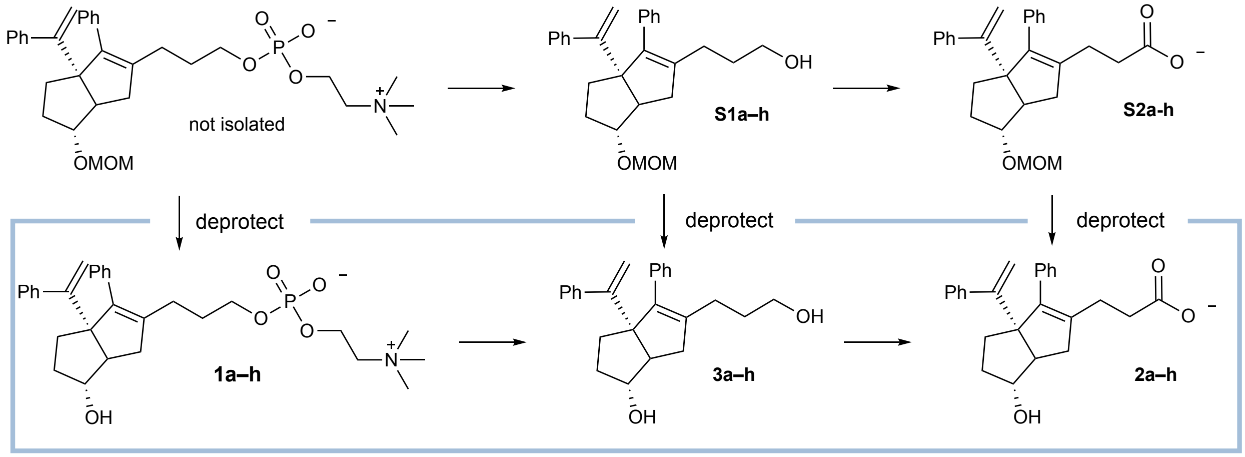

Compounds **S1a-S1h, S2a–h, 1a–h, and 2a–h** were synthesized and purified as described previously.[1]

[1] Flynn, A.R., Mays, S.G., Ortlund, E.A., and Jui, N.T.; *ACS Med. Chem. Lett.* **2018**, *9*(10), 1051–1056.

**Hybrid Precursors: Compounds S1a–S1h**

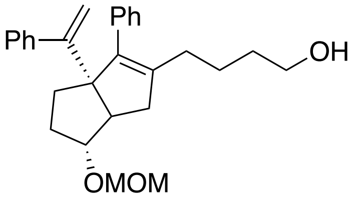

**4-(6-*exo*-(methoxymethoxy)-3-phenyl-3a-(1-phenylvinyl)-1,3a,4,5,6,6a-hexahydropentalen-2-yl)butan-1-ol** (**S1a**): **$^1$H NMR** (400 MHz, $CDCl_3$) δ 7.35 – 7.11 (m, 10H), 5.03 (d, *J* = 1.5 Hz, 1H), 4.99 (d, *J* = 1.5 Hz, 1H), 4.60 – 4.52 (m, 2H), 3.81 – 3.71 (m, 1H), 3.54 (t, *J* = 6.2 Hz, 2H), 3.29 (s, 3H), 2.47 – 2.35 (m, 1H), 2.31 – 2.26 (m, 1H), 2.09 – 1.99 (m, 4H), 1.78 – 1.54 (m, 4H), 1.52 – 1.35 (m, 3H). For the *endo* diastereomer (characteristic signals): **$^1$H NMR** (400 MHz, $CDCl_3$) δ 5.06 (d, *J* = 1.3 Hz, 1H), 4.84 (d, *J* = 1.4 Hz, 1H), 4.00 (td, *J* = 9.8, 6.1 Hz, 1H), 2.64 (dd, *J* = 17.1, 2.4 Hz, 1H), 2.53 (td, *J* = 9.0, 2.2 Hz, 1H). **$^{13}$C NMR** (126 MHz, $CDCl_3$) δ 154.4, 144.0, 140.8, 139.7, 137.3, 129.6, 127.8, 127.70, 127.65, 126.6, 114.9, 94.7, 86.7, 69.1, 62.6, 55.2, 52.8, 40.4, 32.7, 32.4, 31.5, 29.4, 24.0. **LRMS** (ESI, APCI) *m/z*: calc'd for $C_{27}H_{31}O_2$ $[M\text{-}OCH_3]^+$ 387.2, found 386.9.

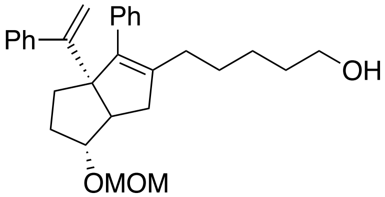

**5-(6-*exo*-(methoxymethoxy)-3-phenyl-3a-(1-phenylvinyl)-1,3a,4,5,6,6a-hexahydropentalen-2-yl)pentan-1-ol** (**S1b**): For the *exo* diastereomer: **$^1$H NMR** (600 MHz, $CDCl_3$) δ 7.41 – 7.10 (m, 10H), 5.01 (d, *J* = 1.4 Hz, 1H), 4.97 (d, *J* = 1.4 Hz, 1H), 4.60 – 4.51 (m, 2H), 3.76 (s, 1H), 3.55 (t, *J* = 6.5 Hz, 2H), 3.27 (s, 3H), 2.39 (d, *J* = 8.6 Hz, 1H), 2.29 (dd, *J* = 18.2, 9.6 Hz, 1H), 2.10 – 1.88 (m, 4H), 1.75 – 1.50 (m, 2H), 1.50 – 1.42 (m, 2H), 1.41 – 1.22 (m, 5H).

For the *endo* diastereomer (characteristic signals): **$^1$H NMR** (600 MHz, $CDCl_3$) δ 5.07 (d, *J* = 1.3 Hz, 1H), 4.84 (d, *J* = 1.4 Hz, 1H), 4.00 (td, *J* = 10.3, 5.8 Hz, 1H), 2.63 (dd, *J* = 17.3, 2.3 Hz, 1H), 2.52 (td, *J* = 9.2, 2.1 Hz, 1H). **LRMS** (ESI, APCI) *m/z*: calc'd for $C_{28}H_{33}O_2$ $[M\text{-}OCH_3]^+$ 401.2, found 401.2.

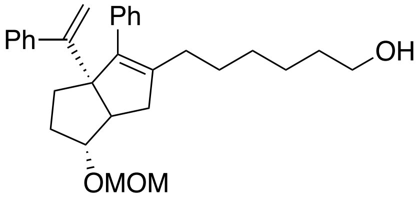

**6-(6-*exo*-(methoxymethoxy)-3-phenyl-3a-(1-phenylvinyl)-1,3a,4,5,6,6a-hexahydropentalen-2-yl)hexan-1-ol** (**S1c**): For the *exo* diastereomer: **$^1$H NMR** (600 MHz, $CDCl_3$) δ 7.36 – 7.11 (m, 10H), 5.03 (s, 1H), 4.98 (s, 1H), 4.61 – 4.51 (m, 2H), 3.77 (s, 1H), 3.57 (t, *J* = 6.7 Hz, 2H), 3.29 (s, 3H), 2.40 (d, *J* = 9.1, 1.8 Hz, 1H), 2.30 (dd, *J* = 16.9, 9.3 Hz, 1H), 2.10 – 1.96 (m, 4H), 1.79 – 1.53 (m, 2H), 1.54 – 1.45 (m, 2H), 1.43 – 1.23 (m, 7H). For the *endo* diastereomer (characteristic signals): **$^1$H NMR** (600 MHz, $CDCl_3$) δ 5.08 (d, *J* = 1.4 Hz, 1H), 4.85 (d, *J* = 1.4 Hz, 1H), 4.01 (td, *J* = 9.8, 5.6 Hz, 1H), 2.65 (dd, *J* = 17.3, 2.2 Hz, 1H), 2.54 (td, *J* = 9.1, 2.3 Hz, 1H).

$^{13}$**C NMR** (126 MHz, $CDCl_3$) δ 154.5, 144.1, 141.1, 139.4, 137.4, 134.8, 129.6, 127.8, 127.63, 126.59, 114.9, 94.7, 86.7, 69.1, 62.8, 55.1, 52.8, 40.5, 32.6, 32.41, 31.42, 29.6, 29.4, 27.8, 25.5. **LRMS** (ESI, APCI) *m/z*: calc'd for $C_{29}H_{35}O_2$ [M-OCH$_3$]$^+$ 415.3, found 415.3.

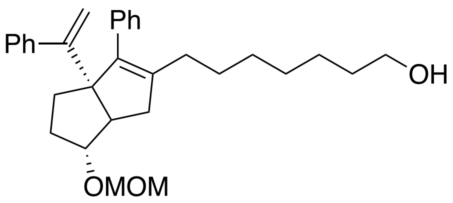

**7-(6-*exo*-(methoxymethoxy)-3-phenyl-3a-(1-phenylvinyl)-1,3a,4,5,6,6a-hexahydropentalen-2-yl)heptan-1-ol** (**S1d**): For the *exo* diastereomer: $^1$**H NMR** (600 MHz, $CDCl_3$) δ 7.46 – 7.02 (m, 10H), 5.05 (s, 1H), 5.00 (s, 1H), 4.62 – 4.56 (m, 2H), 3.79 (s, 1H), 3.66 – 3.60 (m, 2H), 3.30 (s, 3H), 2.41 (d, *J* = 9.4 Hz, 1H), 2.32 (dd, *J* = 17.0, 9.2 Hz, 1H), 2.08 – 1.92 (m, 4H), 1.78 – 1.67 (m, 2H), 1.59 – 1.49 (m, 2H), 1.44 – 1.19 (m, 9H). For the *endo* diastereomer (characteristic signals): $^1$**H NMR** (600 MHz, $CDCl_3$) δ 5.08 (d, *J* = 1.3 Hz, 1H), 4.85 (d, *J* = 1.2 Hz, 1H), 4.00 (td, *J* = 9.6, 5.6 Hz, 1H), 2.65 (dd, *J* = 17.4, 2.1 Hz, 1H), 2.53 (td, *J* = 8.9, 2.2 Hz, 1H).
$^{13}$**C NMR** (126 MHz, $CDCl_3$) δ 154.5, 144.1, 141.2, 139.3, 137.5, 129.6, 127.8, 127.63, 126.57, 114.9, 94.7, 86.7, 69.1, 63.0, 55.1, 52.8, 40.5, 32.7, 32.4, 31.4, 29.64, 29.58, 29.2, 27.7, 25.6. **LRMS** (ESI, APCI) *m/z*: calc'd for $C_{30}H_{37}O_2$ [M-OCH$_3$]$^+$ 429.3, found 428.8.

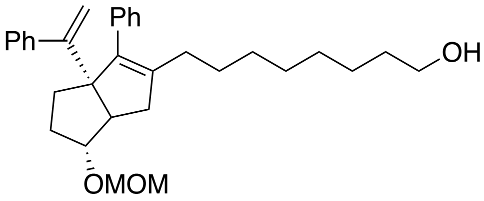

**8-(6-*exo*-(methoxymethoxy)-3-phenyl-3a-(1-phenylvinyl)-1,3a,4,5,6,6a-hexahydropentalen-2-yl)octan-1-ol (S1e)**: For the *exo* diastereomer: $^1$**H NMR** (500 MHz, $CDCl_3$) δ 7.41 – 7.18 (m, 10H), 5.05 (d, *J* = 1.5 Hz, 1H), 5.01 (d, *J* = 1.5 Hz, 1H), 4.63 – 4.53 (m, 2H), 3.79 (s, 1H), 3.62 (t, *J* = 6.7 Hz, 2H), 3.31 (s, 3H), 2.42 (d, *J* = 9.4, 1.8 Hz, 1H), 2.32 (dd, *J* = 16.9, 9.1 Hz, 1H), 2.11 – 1.97 (m, 4H), 1.83 – 1.54 (m, 4H), 1.59 – 1.50 (m, 2H), 1.38 – 1.17 (m, 9H). For the *endo* diastereomer (characteristic signals): $^1$**H NMR** (500 MHz, $CDCl_3$) δ 5.08 (d, *J* = 1.3 Hz, 1H), 4.85 (d, *J* = 1.3 Hz, 1H), 4.01 (td, *J* = 9.6, 5.8 Hz, 1H), 2.65 (dd, *J* = 17.4, 2.1 Hz, 1H), 2.53 (td, *J* = 8.9, 2.2 Hz, 1H). $^{13}$**C NMR** (126 MHz, $CDCl_3$) δ 154.5, 144.1, 141.3, 139.3, 137.5, 129.6, 127.8, 127.6, 126.59, 126.55, 114.8, 94.7, 86.7, 69.1, 63.0, 55.1, 52.8, 40.5, 32.8, 32.4, 31.4, 29.7, 29.6, 29.33, 29.27, 27.8, 25.7. **LRMS** (ESI, APCI) *m/z*: calc'd for $C_{31}H_{39}O_2$ [M-OCH$_3$]$^+$ 443.3, found 442.9.

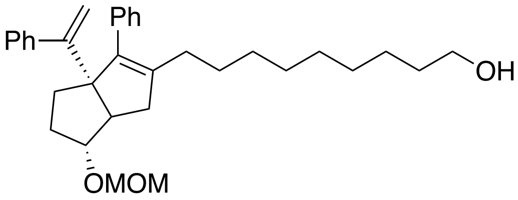

**9-(6-*exo*-(methoxymethoxy)-3-phenyl-3a-(1-phenylvinyl)-1,3a,4,5,6,6a-hexahydropentalen-2-yl)nonan-1-ol** (**S1f**): For the *exo* diastereomer: $^1$**H NMR** (500 MHz, $CDCl_3$) δ 7.39 – 7.15 (m, 10H), 5.06 (d, *J* = 1.4 Hz, 1H), 5.01 (d, *J* = 1.5 Hz, 1H), 4.62 – 4.56 (m, 2H), 3.80 (s, 1H), 3.62 (t, *J* = 6.7 Hz, 2H), 3.32 (s, 3H), 2.43 (d, *J* = 9.1, 1.7 Hz, 1H), 2.33 (dd, *J* = 16.9, 9.2 Hz, 1H), 2.11 – 1.99 (m, 4H), 1.79 – 1.60 (m, 4H), 1.60 – 1.51 (m, 2H), 1.43 – 1.17 (m, 11H).

For the *endo* diastereomer (characteristic signals): **$^1$H NMR** (500 MHz, $CDCl_3$) δ 5.09 (d, *J* = 1.4 Hz, 1H), 4.86 (d, *J* = 1.4 Hz, 1H), 4.01 (td, *J* = 9.6, 5.8 Hz, 1H), 2.66 (dd, *J* = 17.3, 2.4 Hz, 1H), 2.54 (td, *J* = 9.0, 2.3 Hz, 1H). **$^{13}$C NMR** (126 MHz, $CDCl_3$) δ 154.5, 144.1, 141.3, 139.2, 137.5, 129.6, 127.8, 127.6, 126.62, 126.56, 114.9, 94.7, 86.7, 69.1, 63.0, 55.2, 52.7, 40.5, 32.8, 32.4, 31.4, 29.7, 29.6, 29.5, 29.4, 29.3, 27.8, 25.7.**LRMS** (ESI, APCI) *m/z*: calc'd for $C_{32}H_{41}O_2$ $[M-CH_3O]^+$ 457.3, found 457.8.

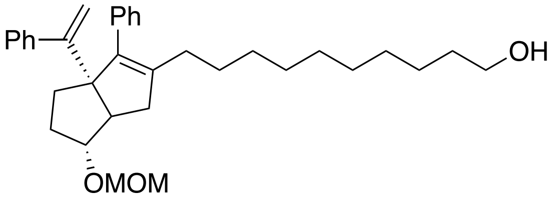

**10-(6-*exo*-(methoxymethoxy)-3-phenyl-3a-(1-phenylvinyl)-1,3a,4,5,6,6a-hexahydropentalen-2-yl)decan-1-ol** (**S1g**): For the *exo* diastereomer: **$^1$H NMR** (500 MHz, $CDCl_3$) δ 7.41 – 7.17 (m, 10H), 5.07 (d, *J* = 1.4 Hz, 1H), 5.02 (d, *J* = 1.5 Hz, 1H), 4.64 – 4.56 (m, 2H), 3.81 (p, *J* = 2.1 Hz, 1H), 3.62 (d, *J* = 6.6 Hz, 2H), 3.32 (s, 3H), 2.44 (dq, *J* = 9.2, 1.7 Hz, 1H), 2.35 (dd, *J* = 16.9, 9.2 Hz, 1H), 2.11 – 2.00 (m, 4H), 1.81 – 1.63 (m, 4H), 1.57 (m, 2H), 1.40 – 1.19 (m, 13H). For the *endo* diastereomer (characteristic signals): **$^1$H NMR** (500 MHz, $CDCl_3$) δ 5.10 (d, *J* = 1.3 Hz, 1H), 4.87 (d, *J* = 1.4 Hz, 1H), 4.03 (td, *J* = 9.7, 5.6 Hz, 1H), 2.68 (dd, *J* = 17.3, 2.3 Hz, 1H), 2.56 (td, *J* = 9.0, 2.3 Hz, 1H). **$^{13}$C NMR** (126 MHz, $CDCl_3$) δ 154.5, 144.1, 141.3, 139.2, 137.5, 129.6, 127.8, 127.6, 126.62, 126.56, 114.9, 94.7, 86.7, 69.1, 63.0, 55.2, 52.7, 40.5, 32.8, 32.4, 31.4, 29.7, 29.6, 29.5, 29.4, 29.3, 27.8, 25.7. **LRMS** (ESI, APCI) *m/z*: calc'd for $C_{33}H_{43}O_2$ $[M-CH_3O]^+$ 471.3, found 470.9.

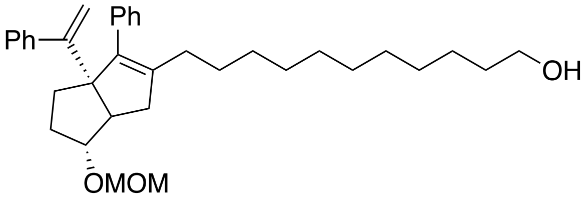

**11-(6-*exo*-(methoxymethoxy)-3-phenyl-3a-(1-phenylvinyl)-1,3a,4,5,6,6a-hexahydropentalen-2-yl)undecan-1-ol** (**S1h**): For the *exo* diastereomer: **$^1$H NMR** (500 MHz, $CDCl_3$) δ 7.42 – 7.16 (m, 10H), 5.08 (d, *J* = 1.5 Hz, 1H), 5.03 (d, *J* = 1.5 Hz, 1H), 4.65 – 4.56 (m, 3H), 3.82 (s, 1H), 3.64 (t, *J* = 6.7 Hz, 2H), 3.33 (s, 3H), 2.45 (d, *J* = 9.2, 1.7 Hz, 1H), 2.36 (dd, *J* = 16.9, 9.2 Hz, 1H), 2.13 – 2.00 (m, 4H), 1.89 – 1.61 (m, 4H), 1.62 – 1.51 (m, 2H), 1.41 – 1.19 (m, 15H).

For the *endo* diastereomer (characteristic signals): **$^1$H NMR** (500 MHz, $CDCl_3$) δ 5.11 (d, *J* = 1.4 Hz, 1H), 4.88 (d, *J* = 1.4 Hz, 1H), 4.03 (td, *J* = 9.8, 5.8 Hz, 1H), 2.68 (dd, *J* = 17.3, 2.3 Hz, 1H), 2.57 (td, *J* = 9.0, 2.3 Hz, 1H). **$^{13}$C NMR** (126 MHz, $CDCl_3$) δ 154.5, 144.1, 141.4, 139.2, 137.5, 129.6, 127.8, 127.7, 126.64, 126.58, 114.9, 94.7, 86.8, 69.1, 62.9, 55.2, 52.7, 40.6, 32.8, 32.5, 31.5, 29.7, 29.7, 29.61, 29.58, 29.54, 29.46, 29.4, 27.9, 25.8. **LRMS** (ESI, APCI) *m/z*: calc'd for $C_{34}H_{45}O_2$ $[M-CH_3O]^+$ 485.3, found 484.9.

**Phosphorylcholines: Compounds 1a–h**

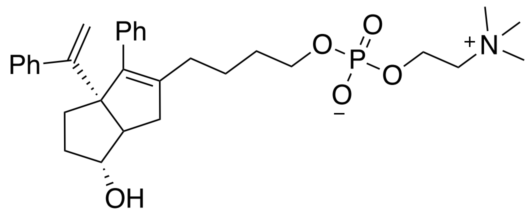

**4-(6-*exo*-hydroxy-3-phenyl-3a-(1-phenylvinyl)-1,3a,4,5,6,6a-hexahydropentalen-2-yl)butyl (2-(trimethylammonio)ethyl) phosphate** (**1a**): **$^1$H NMR** (500 MHz, $CDCl_3$) δ 7.28 – 7.13 (m, 10H), 5.02 (s, 1H), 4.99 (s, 1H), 4.19 (s, 2H), 3.81 (s, 1H), 3.76 (s, 2H), 3.64 (s, 2H), 3.21 (s, 9H), 2.26 – 2.05 (m, 3H), 2.01 – 1.87 (m, 1H), 1.68 – 1.36 (m, 9H). **$^{31}$P NMR (**121 MHz, $CDCl_3$) δ -0.76. **LRMS** (ESI, APCI) *m/z*: calc'd for $C_{31}H_{43}O_5NP$ $[M+H]^+$ : 540.3, found 540.3.

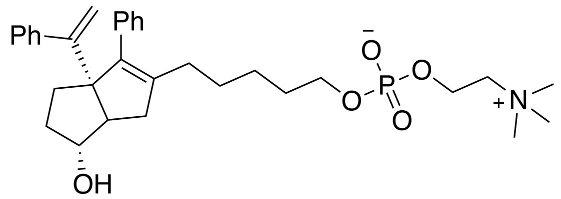

**5-(6-*exo*-hydroxy-3-phenyl-3a-(1-phenylvinyl)-1,3a,4,5,6,6a-hexahydropentalen-2-yl)pentyl (2-(trimethylammonio)ethyl) phosphate (1b): $^1$H NMR** (600 MHz, $CDCl_3$) δ 7.31 – 7.24 (m, 4H), 7.25 – 7.14 (m, 6H), 4.99 (s, 2H), 4.19 (s, 2H), 3.86 (s, 1H), 3.75 (s, 2H), 3.63 (s, 2H), 3.20 (s, 9H), 2.25 – 2.18 (m, 1H), 2.16 – 2.04 (m, 2H), 1.95 – 1.87 (m, 1H), 1.70 – 1.60 (m, 1H), 1.60 – 1.52 (m, 4H), 1.53 – 1.43 (m, 1H), 1.42 – 1.30 (m, 3H), 1.31 – 1.18 (m, 2H). **$^{31}$P NMR** (121 MHz, $CDCl_3$) δ -0.82. **LRMS** (ESI, APCI) *m/z*: calc'd for $C_{32}H_{45}O_5NP$ $[M+H]^+$ : 554.3, found 554.2.

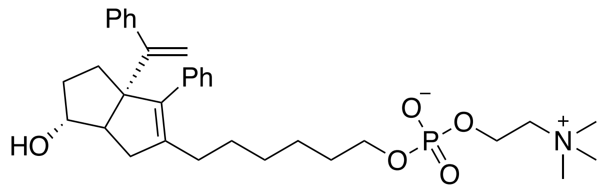

**6-(6-*exo*-hydroxy-3-phenyl-3a-(1-phenylvinyl)-1,3a,4,5,6,6a-hexahydropentalen-2-yl)hexyl (2-(trimethylammonio)ethyl) phosphate** (**1c**): **$^1$H NMR** (500 MHz, $CDCl_3$) δ 7.41 – 7.26 (m, 4H), 7.28 – 7.12 (m, 6H), 5.03 (s, 1H), 5.01 (s, 1H), 4.24 (s, 2H), 3.88 (s, 1H), 3.80 – 3.67 (m, 4H), 3.28 (s, 9H), 2.28 – 2.05 (m, 3H), 2.01 – 1.80 (m, 1H), 1.72 – 1.49 (m, 3H), 1.40 – 0.82 (m, 10H). **$^{31}$P NMR** (121 MHz, Chloroform-*d*) δ -0.59. **LRMS** (ESI, APCI) *m/z*: calc'd for $C_{33}H_{47}O_5NP$ $[M+H]^+$ 568.3, found 568.2.

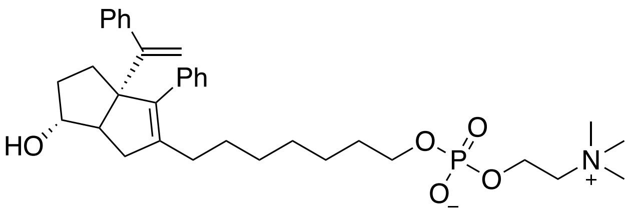

**7-(6-*exo*-hydroxy-3-phenyl-3a-(1-phenylvinyl)-1,3a,4,5,6,6a-hexahydropentalen-2-yl)heptyl (2-(trimethylammonio)ethyl) phosphate** (**1d**): **$^{1}$H NMR** (500 MHz, $CDCl_3$) δ 7.37 – 7.11 (m, 10H), 5.02 (s, 1H), 4.99 (s, 1H), 4.21 (s, 2H), 3.87 (s, 1H), 3.77 (s, 2H), 3.67 (s, 1H), 3.25 (s, 9H), 2.29 – 2.16 (m, 1H), 2.15 – 2.02 (m, 2H), 2.00 – 1.90 (m, 1H), 1.72 – 1.57 (m, 3H), 1.57 – 1.48 (m, 3H), 1.37 – 1.14 (m, 9H). **$^{13}$C NMR** (300 MHz, $CDCl_3$) δ 154.6, 144.1, 140.9, 139.6, 137.4, 129.6, 127.8, 127.7, 126.64, 114.7, 81.5, 69.1, 66.1, 65.7, 59.2, 55.7, 54.3, 40.0, 34.2, 32.0, 30.8, 29.5, 29.3, 29.2, 27.6, 25.6. **$^{31}$P NMR** (300 MHz, $CDCl_3$) δ -0.51. **HRMS** (ESI) *m/z*: calc'd for $C_{34}H_{49}O_5NP$ $[M+H]^+$ : 582.3343, found 582.3338.

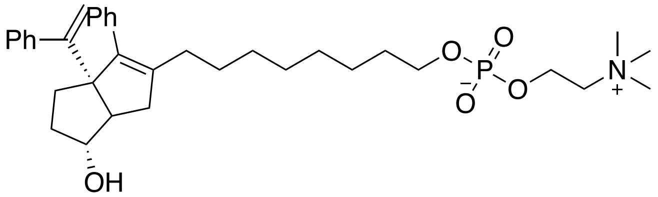

**8-(6-*exo*-hydroxy-3-phenyl-3a-(1-phenylvinyl)-1,3a,4,5,6,6a-hexahydropentalen-2-yl)octyl (2-(trimethylammonio)ethyl) phosphate (1e)**: **$^{1}$H NMR** (600 MHz, $CDCl_3$) δ 7.33 –7.15 (m, 10H), 5.00 (s, 1H), 4.96 (s, 1H), 4.22 (s, 2H), 3.89 (s, 1H), 3.81 – 3.75 (m, 2H), 3.74 – 3.66 (m, 2H), 3.27 (s, 9H), 2.28 – 2.17 (m, 2H), 2.10 – 2.04 (m, 1H), 1.96 – 1.87 (m, 1H), 1.71 – 1.58 (m, 3H), 1.57 – 1.48 (m, 2H), 1.36 – 1.13 (m, 12H). **$^{13}$C NMR** (151 MHz, $CDCl_3$) δ 154.6, 144.1, 141.0, 139.5, 137.4, 129.6, 127.8, 127.6, 126.61, 126.55, 114.8, 81.7, 69.2, 66.3, 59.1, 55.6, 54.4, 53.4, 40.1, 37.1, 34.3, 32.0, 30.8, 29.4, 29.2, 28.9, 28.8, 27.5, 25.6, 22.6. **$^{31}$P NMR** (300 MHz, $CDCl_3$) δ -0.43. **LRMS** (ESI, APCI) *m/z*: calc'd for $C_{35}H_{51}O_5NP$ $[M+H]^+$ 596.3, found 596.3. **HRMS** (ESI) *m/z*: calc'd for $C_{35}H_{51}O_5NP$ $[M+H]^+$ : 596.3499, found 596.3494.

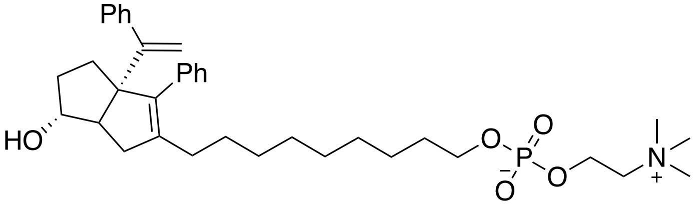

**9-(6-*exo*-hydroxy-3-phenyl-3a-(1-phenylvinyl)-1,3a,4,5,6,6a-hexahydropentalen-2-yl)nonyl (2-(trimethylammonio)ethyl) phosphate** (**1f**): **$^{1}$H NMR** (600 MHz, $CDCl_3$) δ 7.32 – 7.14 (m, 10H), 5.01 (d, *J* = 7.6 Hz, 1H), 4.95 (d, *J* = 7.9 Hz, 1H), 4.20 (s, 2H), 3.87 (s, 1H), 3.75 (s, 3H), 3.67 (s, 3H), 3.25 (s, 9H), 2.27 – 2.18 (m, 2H), 2.06 – 1.98 (m, 2H), 1.94 (p, *J* = 7.0 Hz, 1H), 1.70 – 1.58 (m, 4H), 1.58 – 1.48 (m, 3H), 1.36 – 1.10 (m, 12H). **$^{13}$C NMR** (126 MHz, $CDCl_3$) δ 154.7, 144.2, 141.1, 139.4, 137.4, 129.7, 127.8, 127.7, 127.6, 126.63, 126.57, 114.9, 81.7, 69.3, 66.2, 65.9, 59.3, 55.7, 54.4, 40.2, 34.2, 32.1, 30.9, 29.5, 29.3, 29.2, 27.7, 27.6, 25.8. **$^{31}$P NMR** (121 MHz, $CDCl_3$) δ -0.73. **LRMS** (ESI, APCI) *m/z*: calc'd for $C_{36}H_{53}O_5NP$ $[M+H]^+$ 610.4, found 609.8. **HRMS** (ESI) *m/z*: calc'd for $C_{36}H_{53}O_5NP$ $[M+H]^+$ : 610.3656, found 610.3655.

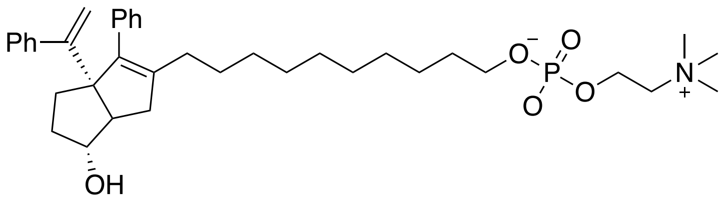

**10-(6-*exo*-hydroxy-3-phenyl-3a-(1-phenylvinyl)-1,3a,4,5,6,6a-hexahydropentalen-2-yl)decyl (2-(trimethylammonio)ethyl) phosphate** (**1g**): **¹H NMR** (600 MHz, $CDCl_3$) δ 7.34 – 7.14 (m, 10H), 5.03 (d, *J* = 4.9 Hz, 1H), 4.95 (d, *J* = 5.0 Hz, 1H), 4.24 (s, 2H), 3.90 (s, 1H), 3.79 (q, *J* = 6.4 Hz, 2H), 3.74 (s, 2H), 3.28 (s, 9H), 2.34 – 2.23 (m, 2H), 2.10 – 1.92 (m, 3H), 1.64 (q, *J* = 10.2, 6.5 Hz, 2H), 1.56 (t, *J* = 7.3 Hz, 2H), 1.37 – 1.10 (m, 16H). **¹³C NMR** (126 MHz, $CDCl_3$) δ 154.7, 144.2, 141.1, 139.4, 137.4, 129.7, 127.73, 127.70, 127.6, 126.63, 126.56, 81.7, 69.3, 66.2, 65.9, 59.3, 55.6, 54.3, 40.2, 34.1, 32.2, 30.91, 30.86, 29.6, 29.3, 29.22, 29.17, 27.6, 25.8. **³¹P NMR** (121 MHz, $CDCl_3$) δ -0.75. **LRMS** (ESI, APCI) *m/z*: calc'd for $C_{37}H_{55}O_5NP$ $[M+H]^+$ : 624.4, found 624.3.

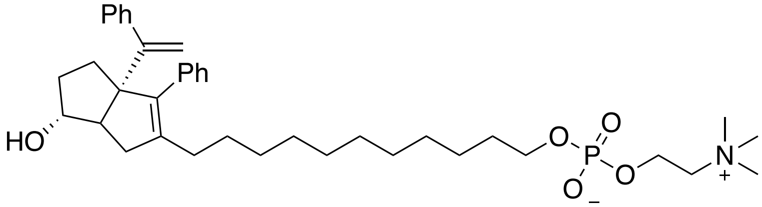

**11-(6-*exo*-hydroxy-3-phenyl-3a-(1-phenylvinyl)-1,3a,4,5,6,6a-hexahydropentalen-2-yl)undecyl (2-(trimethylammonio)ethyl) phosphate** (**1h**): **¹H NMR** (600 MHz, $CDCl_3$ ) δ 7.35 – 7.15 (m, 12H), 5.03 (s, 1H), 4.96 (s, 1H), 4.26 (s, 2H), 3.91 (s, 1H), 3.85 – 3.77 (m, 2H), 3.75 (s, 2H), 3.29 (s, 9H), 2.32 (dd, *J* = 16.7, 9.4 Hz, 1H), 2.26 (d, *J* = 9.6 Hz, 1H), 2.10 – 1.94 (m, 4H), 1.71 – 1.61 (m, 3H), 1.61 – 1.53 (m, 2H), 1.38 – 1.14 (m, 16H). **¹³C NMR** (126 MHz, $CDCl_3$) δ 154.7, 144.2, 141.1, 139.3, 137.4, 129.7, 114.9, 81.7, 69.3, 66.2, 65.9, 59.3, 55.7, 54.4, 40.3, 34.0, 32.2, 31.6, 30.91, 30.86, 29.6, 29.5, 29.43, 29.40, 29.36, 29.3, 29.2, 27.7, 25.8, 22.7. **³¹P NMR** (121 MHz, $CDCl_3$) δ -0.90.m**LRMS** (ESI, APCI) *m/z*: calc'd for $C_{38}H_{57}O_5NP$ $[M+H]^+$ 638.4, found 637.8 **HRMS** (ESI) *m/z*: calc'd for $C_{38}H_{57}O_5NP$ $[M+H]^+$ : 638.3969, found 638.3974.

### Carboxylate Precursors: Compounds S2a–S2h

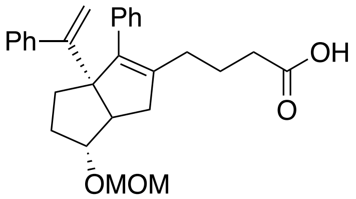

**4-(6-*exo*-(methoxymethoxy)-3-phenyl-3a-(1-phenylvinyl)-1,3a,4,5,6,6a-hexahydropentalen-2-yl)butanoic acid (S2a):** For the *exo* diastereomer: **¹H NMR** (500 MHz, $CDCl_3$) δ 7.47 – 7.05 (m, 10H), 5.07 (s, 1H), 5.01 (s, 1H), 4.66 – 4.52 (m, 2H), 3.80 (s, 1H), 3.31 (s, 3H), 2.45 (d, *J* = 8.9 Hz, 1H), 2.37 – 2.20 (m, 2H), 2.13 – 1.98 (m, 4H), 1.81 – 1.57 (m, 3H), 1.36 – 1.18 (m, 3H). For the *endo* diastereomer (characteristic signals): ¹H NMR (500 MHz, $CDCl_3$) δ 5.08 (s, 1H), 4.88 (s, 1H), 4.06 – 3.98 (m, 1H), 2.67 (d, *J* = 17.8 Hz, 1H). **¹³C NMR** (126 MHz, $cdcl_3$) δ 179.0, 154.3, 143.9, 140.6, 139.7, 137.1, 129.5, 127.80,

127.75, 127.7, 126.8, 115.0, 94.7, 86.6, 69.1, 55.2, 52.8, 40.3, 33.6, 32.4, 31.5, 29.1, 22.9. **LRMS** (ESI, APCI) *m/z*: calc'd for $C_{27}H_{29}O_3$ $[M\text{-}OCH_3]^+$ 401.2, found 400.8.

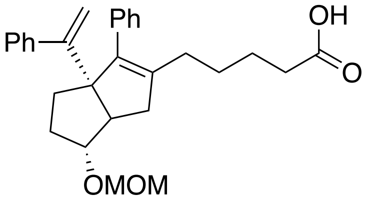

**5-(6-*exo*-(methoxymethoxy)-3-phenyl-3a-(1-phenylvinyl)-1,3a,4,5,6,6a-hexahydropentalen-2-yl)pentanoic acid (S2b):** For the *exo* diastereomer: **$^1$H NMR** (500 MHz, $CDCl_3$) δ 7.37 – 7.14 (m, 10H), 5.06 (d, *J* = 1.4 Hz, 1H), 5.02 (d, *J* = 1.4 Hz, 1H), 4.63 – 4.57 (m, 2H), 3.80 (s, 1H), 3.32 (s, 3H), 2.43 (d, *J* = 9.2 Hz, 1H), 2.35 – 2.26 (m, 3H), 2.10 – 2.00 (m, 4H), 1.79 – 1.53 (m, 4H), 1.46 – 1.20 (m, 3H). For the *endo* diastereomer (characteristic signals): **$^1$H NMR** (500 MHz, $CDCl_3$) δ 5.07 (d, *J* = 1.3 Hz, 1H), 4.85 (d, *J* = 1.3 Hz, 1H), 3.99 (dt, *J* = 9.6, 5.8 Hz, 1H), 2.63 (dd, *J* = 17.2, 1.8 Hz, 1H), 2.54 (dd, *J* = 9.5, 2.2 Hz, 1H). **$^{13}$C NMR** (151 MHz, $CDCl_3$) δ 179.5, 154.3, 144.0, 140.4, 139.9, 137.2, 129.5, 127.8, 127.7, 127.6, 126.7, 126.6, 114.9, 86.6, 69.1, 55.1, 52.7, 40.3, 32.4, 31.4, 29.9, 29.3, 27.2, 24.6. **LRMS** (ESI, APCI) *m/z*: calc'd for $C_{29}H_{33}O_4$ $[M\text{-}H]^-$ 445.2, found 445.1. Calc'd for $C_{28}H_{31}O_3$ $[M\text{-}OCH_3]^+$ 415.2, found 415.2.

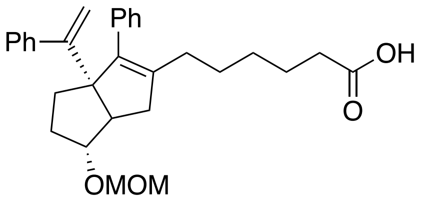

**6-(6-*exo*-(methoxymethoxy)-3-phenyl-3a-(1-phenylvinyl)-1,3a,4,5,6,6a-hexahydropentalen-2-yl)hexanoic acid (S2c):** For the *exo* diastereomer: **$^1$H NMR** (500 MHz, $CDCl_3$) δ 7.36 – 7.11 (m, 10H), 5.05 (d, *J* = 1.4 Hz, 1H), 5.02 (d, *J* = 1.5 Hz, 1H), 3.80 (s, 1H), 3.33 (s, 3H), 2.41 (d, *J* = 8.7 Hz, 1H), 2.38 – 2.29 (m, 2H), 2.24 (dd, *J* = 17.0, 8.9 Hz, 1H), 2.12 – 1.98 (m, 4H), 1.79 – 1.49 (m, 7H), 1.47 – 1.18 (m, 4H). For the *endo* diastereomer: **$^1$H NMR** (600 MHz, $CDCl_3$) δ 5.06 (d, *J* = 1.3 Hz, 1H), 4.84 (d, *J* = 1.3 Hz, 1H), 4.00 (td, *J* = 9.8, 5.8 Hz, 1H), 2.63 (dd, *J* = 17.4, 2.2 Hz, 1H), 2.53 (td, *J* = 9.0, 2.2 Hz, 1H). **$^{13}$C NMR** (151 MHz, $CDCl_3$) δ 179.5, 144.1, 140.7, 139.5, 137.2, 129.6, 127.72, 127.69, 127.66, 126.66, 126.65, 115.0, 82.0, 69.3, 60.4, 53.4, 40.1, 34.0, 32.0, 31.6, 29.4, 29.0, 27.4, 27.0, 22.6, 21.1. **LRMS** (ESI, APCI) *m/z*: calc'd for $C_{30}H_{35}O_4$ $[M\text{-}H]^-$ 459.3, found 459.3. Calc'd for $C_{29}H_{33}O_3$ $[M\text{-}OCH_3]^+$ 429.2, found 429.3.

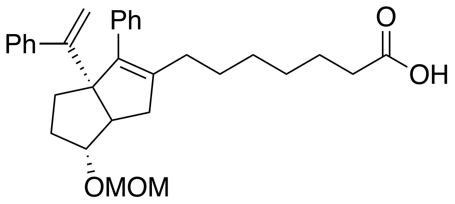

**7-(6-*exo*-(methoxymethoxy)-3-phenyl-3a-(1-phenylvinyl)-1,3a,4,5,6,6a-hexahydropentalen-2-yl)heptanoic acid (S2d):** **$^1$H NMR** (500 MHz, $CDCl_3$) δ 7.60 – 7.02 (m, 10H), 5.05 (d, *J* = 1.5 Hz, 1H), 5.00 (d, *J* = 1.5 Hz, 1H), 4.66 – 4.52 (m, 2H), 3.79 (s, 1H), 3.31 (s, 3H), 2.42 (d, *J* = 9.3, 1.7 Hz, 1H), 2.40 – 2.27 (m, 3H), 2.10 – 1.97 (m, 4H), 1.79 – 1.50 (m, 4H), 1.52 – 1.18 (m, 7H). For the *endo* diastereomer (characteristic signals): **$^1$H NMR** (500 MHz, Chloroform-*d*) δ 5.08 (d, *J* = 1.4 Hz, 1H), 4.85 (d, *J* = 1.4 Hz,

1H), 4.01 (td, *J* = 9.1, 5.8 Hz, 1H), 2.64 (d, *J* = 17.5 Hz, 1H), 2.54 (t, *J* = 9.1 Hz, 1H). **[13]C NMR** (126 MHz, $CDCl_3$) δ 179.5, 154.5, 144.1, 141.1, 139.4, 137.4, 134.6, 129.6, 127.8, 127.7, 126.6, 114.9, 94.7, 86.7, 69.1, 55.1, 52.8, 40.5, 33.9, 32.4, 31.4, 29.6, 29.2, 28.8, 27.6, 24.5. **LRMS** (ESI, APCI) *m/z*: calc'd for $C_{31}H_{37}O_4$ $[M-H]^-$ 473.3, found 473.4.

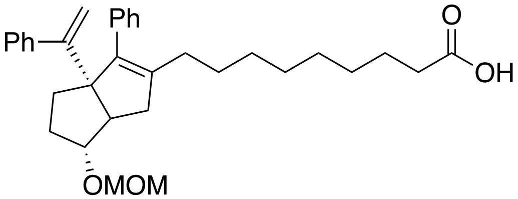

**9-(6-*exo*-(methoxymethoxy)-3-phenyl-3a-(1-phenylvinyl)-1,3a,4,5,6,6a-hexahydropentalen-2-yl)nonanoic acid** (**S2f**): For the *exo* diastereomer: **[1]H NMR** (500 MHz, $CDCl_3$) δ 7.39 – 7.18 (m, 10H), 5.05 (d, *J* = 1.4 Hz, 1H), 5.00 (d, *J* = 1.5 Hz, 1H), 4.65 – 4.57 (m, 2H), 3.80 (d, *J* = 2.1 Hz, 1H), 3.32 (d, *J* = 0.7 Hz, 3H), 2.42 (d, 1H), 2.37 – 2.29 (m, 3H), 2.10 – 1.98 (m, 4H), 1.69 – 1.58 (m, 5H), 1.44 – 1.08 (m, 11H). For the *endo* diastereomer (characteristic signals): **[1]H NMR** (500 MHz, $CDCl_3$) δ 5.09 (d, *J* = 1.3 Hz, 1H), 4.85 (d, *J* = 1.4 Hz, 1H), 4.02 (td, *J* = 9.8, 5.4 Hz, 1H), 2.65 (dd, *J* = 17.7, 2.1 Hz, 1H), 2.54 (td, *J* = 9.4, 2.1 Hz, 1H). **[13]C NMR** (126 MHz, $CDCl_3$) δ 179.2, 154.5, 144.1, 141.3, 139.3, 137.4, 129.6, 127.8, 127.6, 126.61, 126.57, 114.9, 94.6, 86.8, 69.1, 55.1, 52.8, 40.5, 33.9, 32.4, 31.4, 29.7, 29.5, 29.2, 29.1, 29.0, 27.7, 24.7. **LRMS** (ESI, APCI) *m/z*: calc'd for $C_{33}H_{41}O_4$ $[M-H]^-$ 501.3, found 501.4. Calc'd for $C_{32}H_{39}O_3$ $[M-CH_3O]^+$ 471.3, found 470.8.

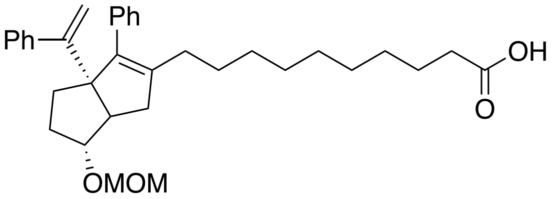

**10-(6-*exo*-(methoxymethoxy)-3-phenyl-3a-(1-phenylvinyl)-1,3a,4,5,6,6a-hexahydropentalen-2-yl)decanoic acid (S2g)**: For the *exo* diastereomer: **[1]H NMR** (500 MHz, $CDCl_3$) δ 7.38 – 7.18 (m, 10H), 5.05 (d, *J* = 1.4 Hz, 1H), 5.01 (d, *J* = 1.5 Hz, 1H), 4.61 – 4.59 (m, 2H), 3.80 (s, 1H), 3.32 (s, 3H), 2.42 (d, *J* = 9.2, 1.7 Hz, 1H), 2.38 – 2.29 (m, 3H), 2.11 – 1.97 (m, 4H), 1.70 – 1.59 (m, 5H), 1.37 – 1.18 (m, 13H). For the *endo* diastereomer (characteristic signals): **[1]H NMR** (500 MHz, $CDCl_3$) δ 5.09 (d, *J* = 1.3 Hz, 1H), 4.86 (d, *J* = 1.3 Hz, 1H), 4.02 (td, *J* = 9.7, 5.5 Hz, 1H), 2.65 (dd, *J* = 17.7, 2.2 Hz, 1H), 2.54 (td, *J* = 9.3, 2.2 Hz, 1H). **[13]C NMR** (126 MHz, $CDCl_3$) δ 179.4, 154.5, 144.1, 141.3, 139.2, 137.5, 129.6, 127.8, 127.6, 126.61, 126.56, 114.9, 94.7, 86.7, 69.1, 55.1, 52.7, 40.5, 34.0, 32.4, 31.4, 29.7, 29.6, 29.30, 29.26, 29.2, 29.0, 27.8, 24.7. **LRMS** (ESI, APCI) *m/z*: calc'd for $C_{34}H_{43}O_4$ $[M-H]^-$ 515.3, found 515.1. Calc'd for $C_{33}H_{41}O_3$ $[M-CH_3O]^+$ 485.3, found 484.9.

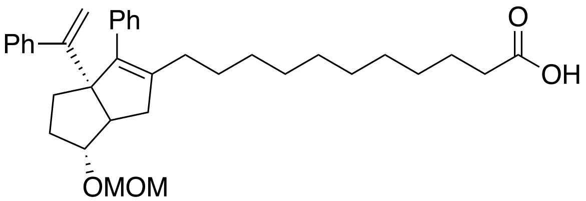

**11-(6-*exo*-(methoxymethoxy)-3-phenyl-3a-(1-phenylvinyl)-1,3a,4,5,6,6a-hexahydropentalen-2-yl)undecanoic acid** (**S2h**): For the *exo* diastereomer: **$^{1}$H NMR** (500 MHz, $CDCl_3$) δ 7.39 – 7.18 (m, 10H), 5.05 (d, *J* = 1.2 Hz, 1H), 5.00 (d, *J* = 1.2 Hz, 1H), 4.64 – 4.58 (m, 2H), 3.80 (s, 1H), 3.32 (s, 3H), 2.42 (d, *J* = 8.8 Hz, 1H), 2.38 – 2.30 (m, 3H), 2.11 – 1.98 (m, 4H), 1.72 – 1.59 (m, 5H), 1.44 – 1.17 (m, 16H). For the *endo* diastereomer (characteristic signals): **$^{1}$H NMR** (500 MHz, Chloroform-*d*) δ 5.09 (d, *J* = 1.2 Hz, 1H), 4.86 (d, *J* = 1.2 Hz, 1H), 4.02 (td, *J* = 9.4, 5.7 Hz, 1H), 2.66 (dd, *J* = 17.6, 2.2 Hz, 1H), 2.54 (td, *J* = 9.9, 2.1 Hz, 1H). **$^{13}$C NMR** (126 MHz, $CDCl_3$) δ 179.2, 154.5, 144.1, 141.4, 139.2, 137.5, 129.6, 127.8, 127.63, 126.60, 126.55, 114.9, 94.7, 86.7, 69.1, 55.1, 52.7, 40.5, 33.9, 32.4, 31.4, 29.7, 29.6, 29.38, 29.36, 29.3, 29.2, 29.0, 27.8, 24.7. **LRMS** (ESI, APCI) *m/z*: calc'd for $C_{35}H_{46}O_4$ $[M-H]^-$ 529.3, found 529.5. Calc'd for $C_{34}H_{43}O_3$ $[M-CH_3O]^+$ 499.3, found 498.9.

**Carboxylates: Compounds 2a–2h**

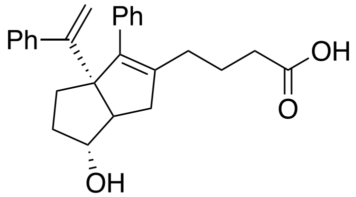

**4-(6-*exo*-hydroxy-3-phenyl-3a-(1-phenylvinyl)-1,3a,4,5,6,6a-hexahydropentalen-2-yl)butanoic acid** (**2a**): **$^{1}$H NMR** (500 MHz, $CDCl_3$) δ 7.38 – 7.16 (m, 10H), 5.08 (d, *J* = 1.4 Hz, 1H), 5.00 (d, *J* = 1.4 Hz, 1H), 3.97 (s, 1H), 2.42 – 2.36 (m, 1H), 2.37 – 2.31 (m, 2H), 2.30 – 2.23 (m, 2H), 2.14 – 2.06 (m, 4H), 1.75 – 1.65 (m, 5H). **$^{13}$C NMR** (126 MHz, $CDCl_3$) δ 178.7, 154.4, 144.0, 140.4, 139.6, 137.0, 129.6, 127.8, 127.7, 126.81, 126.75, 115.2, 82.0, 69.4, 55.7, 40.1, 34.0, 33.8, 32.0, 29.1, 22.9. **LRMS** (ESI, APCI) *m/z*: calc'd for $C_{26}H_{27}O_3$ $[M-H]^-$ 387.2, found 387.1. **HRMS** calc'd for $C_{26}H_{27}O_3$ $[M-H]^-$ : 387.1966, found 387.1969.

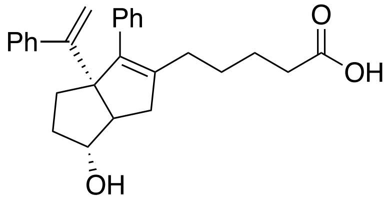

**5-(6-hydroxy-3-phenyl-3a-(1-phenylvinyl)-1,3a,4,5,6,6a-hexahydropentalen-2-yl)pentanoic acid (2b):** For the *exo* diastereomer: **$^{1}$H NMR** (600 MHz, $CDCl_3$) δ 7.56 – 6.96 (m, 10H), 5.05 (d, *J* = 1.4 Hz, 1H), 4.97 (d, *J* = 1.4 Hz, 1H), 3.93 (s, 1H), 2.38 – 2.23 (m, 4H), 2.13 – 1.97 (m, 5H), 1.74 – 1.62 (m, 3H), 1.53 (p, *J* = 7.4 Hz, 2H), 1.44 – 1.32 (m, 2H).

For the *endo* diastereomer (characteristic signals): $^{1}$H NMR (600 MHz, $CDCl_3$) δ 5.07 (d, *J* = 1.6 Hz, 1H), 4.94 (d, *J* = 1.4 Hz, 1H), 4.20 (td, *J* = 9.0, 5.7 Hz, 1H), 2.64 (dd, *J* = 17.3, 2.0 Hz, 1H), 2.50 (td, *J* = 8.6, 1.9 Hz, 1H). **$^{13}$C NMR** (126 MHz, $CDCl_3$) δ 177.8, 154.5, 144.1, 140.3, 139.8, 137.2, 129.7, 127.8, 127.7, 126.8, 126.7, 115.1, 82.0, 69.4, 55.8, 40.1, 34.0, 33.5, 32.1, 29.3, 27.2, 24.6. **LRMS** (ESI, APCI) *m/z*: calc'd for $C_{27}H_{30}O_3$ $[M-H]^-$ 401.2, found 401.5. Calc'd for $C_{27}H_{29}O_2$ $[M-OH]^+$ 385.2, found 385.2 **HRMS** calc'd for $C_{27}H_{31}O_3$ $[M+H]^+$ : 403.2268, found 403.2266.

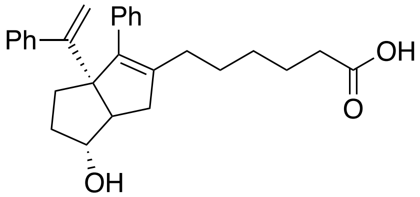

**6-(6-*exo*-hydroxy-3-phenyl-3a-(1-phenylvinyl)-1,3a,4,5,6,6a-hexahydropentalen-2-yl)hexanoic acid** (**2c**): **$^{1}$H NMR** (600 MHz, $CDCl_3$) δ 7.49 – 7.05 (m, 10H), 5.05 (d, *J* = 1.5 Hz, 1H), 4.97 (d, *J* = 1.4 Hz, 1H), 3.93 (s, 1H), 2.36 – 2.21 (m, 4H), 2.10 – 1.97 (m, 5H), 1.73 – 1.61 (m, 3H), 1.59 – 1.51 (m, 2H), 1.34 (p, *J* = 7.6 Hz, 2H), 1.28 – 1.20 (m, 2H). **$^{13}$C NMR** (126 MHz, $cdcl_3$) δ 179.4, 154.5, 144.1, 140.7, 139.5, 137.3, 129.6, 127.8, 127.70, 127.68, 126.7, 115.0, 82.1, 69.3, 55.7, 40.2, 34.0, 33.8, 32.0, 29.5, 29.0, 27.4, 24.5. **LRMS** (ESI, APCI) *m/z*: calc'd for $C_{28}H_{31}O_3$ [M-H]$^-$ 415.2, found 414.9. Calc'd for $C_{28}H_{31}O_2$ [M-OH]$^+$ 399.2, found 399.2 **HRMS** calc'd for $C_{28}H_{32}O_3Na$ [M+Na]$^+$ : 439.2235, found 439.2237.

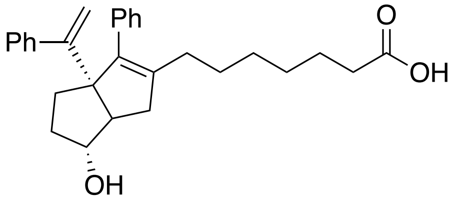

**7-(6-*exo*-hydroxy-3-phenyl-3a-(1-phenylvinyl)-1,3a,4,5,6,6a-hexahydropentalen-2-yl)heptanoic acid** (**2d**): **$^{1}$H NMR** (600 MHz, $CDCl_3$) δ 7.37 – 7.14 (m, 9H), 5.05 (d, *J* = 1.5 Hz, 1H), 4.97 (d, *J* = 1.4 Hz, 1H), 3.93 (s, 1H), 2.38 – 2.25 (m, 4H), 2.11 – 1.97 (m, 4H), 1.76 – 1.47 (m, 4H), 1.33 (p, *J* = 7.6 Hz, 2H), 1.29 – 1.14 (m, 6H). **$^{13}$C NMR** (126 MHz, $CDCl_3$) δ 179.9, 154.5, 144.1, 140.9, 139.3, 137.3, 129.7, 127.74, 127.70, 127.66, 126.7, 115.0, 82.1, 69.3, 55.6, 40.2, 34.0, 33.9, 32.0, 29.6, 29.2, 28.8, 27.6, 24.6. **LRMS** (ESI, APCI) *m/z*: calc'd for $C_{29}H_{33}O_3$ [M-H]$^-$ 429.3, found 429.3. Calc'd for $C_{29}H_{33}O_2$ [M-OH]$^+$: 413.2, found 413.28.

**HRMS** calc'd for $C_{29}H_{33}O_3$ [M-H]$^-$ : 429.2435, found 429.2378.

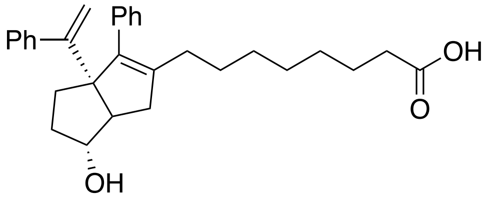

**8-(6-*exo*-hydroxy-3-phenyl-3a-(1-phenylvinyl)-1,3a,4,5,6,6a-hexahydropentalen-2-yl)octanoic acid** (**2e**): **$^{1}$H NMR** (600 MHz, $CDCl_3$) δ 7.81 – 7.13 (m, 10H), 5.05 (s, 1H), 4.97 (s, 1H), 3.93 (s, 1H), 2.37 – 2.22 (m, 5H), 2.09 – 1.94 (m, 5H), 1.79 – 1.58 (m, 6H), 1.38 – 1.16 (m, 5H). **$^{13}$C NMR** (126 MHz, $CDCl_3$) δ 180.0, 154.5, 144.1, 141.0, 139.2, 137.3, 129.7, 127.74, 127.68, 127.6, 126.7, 115.0, 82.1, 69.3, 55.6, 40.2, 34.0, 33.8, 32.0, 29.4, 28.94, 28.88, 27.7, 24.6, 21.0. **LRMS** (ESI, APCI) *m/z*: calc'd for $C_{30}H_{35}O_3$ [M-H]$^-$ 443.3, found 443.2. Calc'd for $C_{30}H_{35}O_2$ [M-OH]$^+$ 427.3, found 427.4. **HRMS** calc'd for $C_{30}H_{35}O_3$ [M-H]$^-$ : 443.2592, found 443.2593.

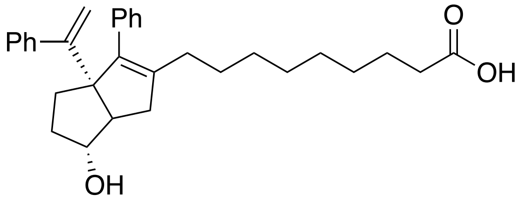

**9-(6-*exo*-hydroxy-3-phenyl-3a-(1-phenylvinyl)-1,3a,4,5,6,6a-hexahydropentalen-2-yl)nonanoic acid** (**2f**): **$^{1}$H NMR** (500 MHz, $CDCl_3$) δ 7.41 – 7.15 (m, 10H), 5.07 (d, *J* = 1.4 Hz, 1H), 4.99 (d, *J* = 1.4 Hz, 1H), 3.95 (s, 1H), 2.39 – 2.27 (m, 5H), 2.12 – 1.97 (m, 5H), 1.76 – 1.56 (m, 6H), 1.40 – 1.15 (m, 7H). **$^{13}$C NMR** (126 MHz, $CDCl_3$) δ 179.5, 154.6, 144.1, 141.1, 139.2, 137.4, 129.7, 127.74, 127.71, 127.6, 126.7, 126.6, 115.0, 82.1, 69.3, 55.7, 40.2, 34.0, 32.1, 29.6, 29.5, 29.2, 29.1, 29.0, 24.7. **LRMS** (ESI, APCI) *m/z*: calc'd for $C_{31}H_{37}O_3$ [M-H]$^-$ 457.3, found 457.3. Calc'd for $C_{31}H_{37}O_2$ [M-OH]$^+$ 441.3, found 440.8. **HRMS** (ESI) *m/z* : calc'd for $C_{31}H_{37}O_3$ [M-H]$^-$ : 457.2748, found 457.2749.

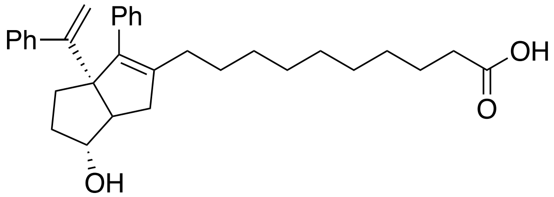

**10-(6-*exo*-hydroxy-3-phenyl-3a-(1-phenylvinyl)-1,3a,4,5,6,6a-hexahydropentalen-2-yl)decanoic acid (6HP-CA 2g)**: **$^{1}$H NMR** (500 MHz, $CDCl_3$) δ 7.39 – 7.15 (m, 10H), 5.07 (d, *J* = 1.6 Hz, 1H), 4.99 (d, *J* = 1.6 Hz, 1H), 3.95 (s, 1H), 2.41 – 2.24 (m, 5H), 2.15 – 1.94 (m, 5H), 1.75 – 1.50 (m, 6H), 1.42 – 1.09 (m, 9H). **$^{13}$C NMR** (126 MHz, $CDCl_3$) δ 179.4, 154.6, 144.2, 141.2, 139.1, 137.4, 129.7, 127.74, 127.71, 127.6, 126.7, 126.6, 115.0, 82.1, 69.3, 55.7, 40.3, 34.0, 33.9, 32.1, 29.7, 29.6, 29.30, 29.28, 29.2, 29.0, 27.8, 24.7. **LRMS** (ESI, APCI) *m/z*: calc'd for $C_{32}H_{39}O_3$ [M-H]$^-$ 471.3, found 471.3. Calc'd for $C_{32}H_{39}O_2$ [M-OH]$^+$: 455.3, found 454.8. **HRMS** (ESI) *m/z*: calc'd for $C_{32}H_{39}O_3$ [M-H]$^-$ : 471.2882, found 471.2905. **FT-IR** (neat): 3360(b), 1708 (s) cm$^{-1}$.

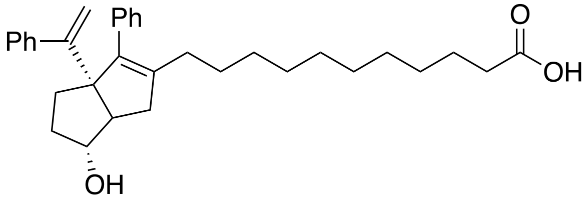

**11-(6-*exo*-hydroxy-3-phenyl-3a-(1-phenylvinyl)-1,3a,4,5,6,6a-hexahydropentalen-2-yl)undecanoic acid** (**2h**): **$^{1}$H NMR** (500 MHz, $CDCl_3$) δ 7.41 – 7.12 (m, 10H), 5.07 (d, *J* = 1.4 Hz, 1H), 4.99 (d, *J* = 1.5 Hz, 1H), 3.99 – 3.92 (s, 1H), 2.38 – 2.26 (m, 5H), 2.11 – 1.98 (m, 5H), 1.75 – 1.58 (m, 6H), 1.36 – 1.17 (m, 11H). **$^{13}$C NMR** (126 MHz, $CDCl_3$) δ 179.4, 154.6, 144.2, 141.2, 139.1, 137.4, 129.7, 127.74, 127.71, 127.6, 126.7, 126.6, 115.0, 82.1, 69.3, 55.7, 40.2, 34.0, 33.9, 32.1, 29.7, 29.6, 29.4, 29.3, 29.2, 29.0, 27.8, 24.7. **LRMS** (ESI, APCI) *m/z*: calc'd for $C_{33}H_{41}O_3$ [M-H]$^-$ 485.3, found 485.3. Calc'd for $C_{33}H_{41}O_2$ [M-OH]$^+$: 469.3, found 468.9. **HRMS** (ESI) *m/z* : calc'd for $C_{33}H_{41}O_3$ [M-H]$^-$ : 485.3061, found 485.3065.

### Diols: Compounds 3a–3h

**General Procedure for Diol MOM Deprotection**

To a solution of **(S1a-h)** in MeCN was added concentrated HCl in excess (5-20 equiv.). The solution was stirred for 5 minutes or until the reaction was completed by TLC/LCMS. The resulting solution as concentrated *in vacuo* and subjected to preparatory HPLC to isolate the desired product as the major *exo* diastereomer.

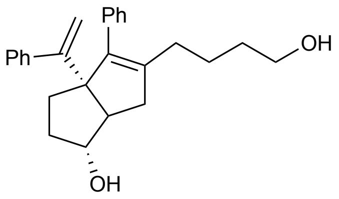

***Exo*-5-(4-hydroxybutyl)-4-phenyl-3a-(1-phenylvinyl)-1,2,3,3a,6,6a-hexahydropentalen-1-ol (3a): S1a** (18.3 mg, 0.04 mmol) was reacted and purified according to the general procedure to give the title compound as a clear, colorless oil (13.0 mg, 79% yield). **$^1$H NMR** (600 MHz, Chloroform-*d*) δ 7.33 – 7.24 (m, 6H), 7.23 (s, 2H), 7.20 – 7.16 (m, 2H), 5.05 (d, *J* = 1.4 Hz, 1H), 4.97 (d, *J* = 1.5 Hz, 1H), 3.93 (s, 1H), 3.55 (t, *J* = 6.2 Hz, 2H), 2.36 (dd, *J* = 16.9, 9.3 Hz, 1H), 2.29 (d, *J* = 6.9 Hz, 1H), 2.10 – 2.02 (m, 4H), 1.74 – 1.61 (m, 3H), 1.50 – 1.37 (m, 4H). **$^{13}$C NMR** (500 MHz, Chloroform-*d*) δ 154.6, 144.1, 140.8, 139.4, 137.3, 129.6, 127.7, 127.7, 127.6, 126.6, 116.6, 114.8, 81.8, 69.3, 62.4, 55.7, 40.1, 33.9, 32.7, 32.0, 29.4, 24.0, 22.5, 10.6. **HRMS** calcd for $C_{26}H_{31}O_2$ $[M+H]^+$ : 375.23186, found 375.23145.

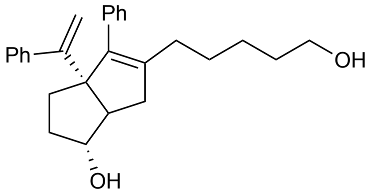

***Exo*-5-(5-hydroxypentyl)-4-phenyl-3a-(1-phenylvinyl)-1,2,3,3a,6,6a-hexahydropentalen-1-ol (3b): S1b** (21.2 mg, 0.05 mmol) was reacted and purified according to the general procedure to give the title compound as a clear, colorless oil (19.0 mg, >99% yield). **$^1$H NMR** (500 MHz, Chloroform-*d*) δ 7.43 – 7.28 (m, 7H), 7.28 – 7.19 (m, 3H), 5.09 (d, *J* = 1.4 Hz, 1H), 5.01 (d, *J* = 1.4 Hz, 1H), 3.97 (s, 1H), 3.61 (t, *J* = 6.6 Hz, 2H), 2.38 (dd, *J* = 16.8, 9.4 Hz, 1H), 2.31 (d, *J* = 9.3, 1.5 Hz, 1H), 2.16 – 2.03 (m, 4H), 1.77 – 1.65 (m, 3H), 1.56 – 1.48 (m, 3H), 1.43 – 1.25 (m, 3H). **$^{13}$C NMR** (500 MHz, Chloroform-*d*) δ 154.5, 144.1, 140.8, 139.3, 137.3, 131.5, 129.6, 128.2, 127.7, 127.6, 126.7, 126.6, 115.0, 82.0, 69.3, 62.9, 61.9, 55.8, 40.3, 34.0, 32.5, 32.1, 31.3, 29.7, 27.6, 25.8, 16.0. **HRMS** calc'd for $C_{27}H_{33}O_2$ $[M+H]^+$ : 389.24751, found 398.24762.

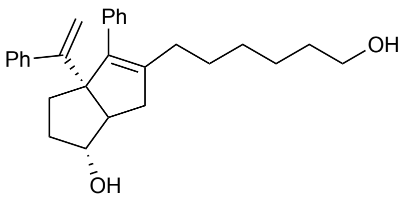

***Exo*-5-(6-hydroxyhexyl)-4-phenyl-3a-(1-phenylvinyl)-1,2,3,3a,6,6a-hexahydropentalen-1-ol (3c): S1c (**9.5 mg, 0.02 mmol) was reacted and purified according to the general procedure to give the title compound as a clear, colorless oil (6.8 mg, 79% yield). **$^1$H NMR** (600 MHz, Chloroform-*d*) δ 7.36 – 7.25 (m, 6H), 7.24 – 7.15 (m, 4H), 5.05 (s, 1H), 4.97 (s, 1H), 3.93 (s, 1H), 3.58 (t, *J* = 6.6 Hz, 2H), 2.35 (dd, *J* =

16.9, 9.4 Hz, 1H), 2.28 (d, *J* = 9.4 Hz, 1H), 2.11 – 1.97 (m, 4H), 1.75 – 1.62 (m, 3H), 1.53 – 1.46 (m, 2H), 1.38 – 1.15 (m, 6H). **$^{13}$C NMR** (600 MHz, Chloroform-*d*) δ 154.5, 144.1, 140.9, 139.2, 137.3, 129.7, 127.7, 127.6, 126.7, 126.6, 115.0, 110.0, 82.0, 69.3, 63.0, 55.8, 40.3, 34.0, 32.6, 32.1, 29.6, 29.4, 27.7, 25.5. **HRMS** calc'd for $C_{28}H_{35}O_2$ $[M+H]^+$ : 403.26316, found 403.26338.

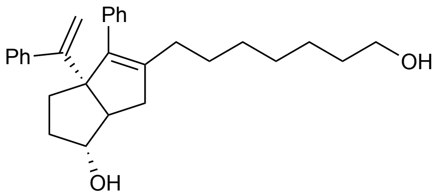

***Exo*-5-(7-hydroxyheptyl)-4-phenyl-3a-(1-phenylvinyl)-1,2,3,3a,6,6a-hexahydropentalen-1-ol (3d): S1d** (11.6 mg, 0.03 mmol) was reacted and purified according to the general procedure to give the title compound as a clear, colorless oil (8.5 mg, 81% yield). **$^{1}$H NMR** (600 MHz, Chloroform-*d*) δ 7.35 – 7.25 (m, 5H), 7.24 – 7.15 (m, 5H), 5.05 (d, *J* = 1.4 Hz, 1H), 4.97 (d, *J* = 1.4 Hz, 1H), 3.93 (s, 1H), 3.60 (t, *J* = 6.7 Hz, 2H), 2.35 (dd, *J* = 16.9, 9.3 Hz, 1H), 2.28 (d, *J* = 9.4 Hz, 1H), 2.09 – 1.98 (m, 5H), 1.73 – 1.61 (m, 3H), 1.36 – 1.19 (m, 10H). **$^{13}$C NMR** (600 MHz, Chloroform-*d*) δ 154.5, 144.1, 141.0, 139.2, 137.3, 129.7, 127.7, 127.6, 126.64, 126.59, 115.0, 82.0, 69.3, 63.0, 58.5, 55.8, 50.9, 40.2, 34.0, 32.7, 32.1, 29.63, 29.58, 29.2, 27.7, 25.6, 18.4. **HRMS** calc'd for $C_{29}H_{36}O_2Cl$ $[M+Cl]^-$ : 451.24093, found 451.24179.

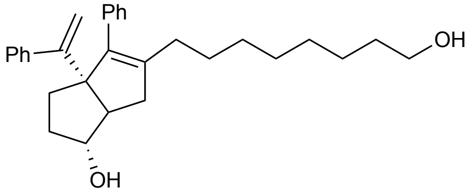

***Exo*-5-(8-hydroxyoctyl)-4-phenyl-3a-(1-phenylvinyl)-1,2,3,3a,6,6a-hexahydropentalen-1-ol (3e): S1e** (4.6 mg, 0.01 mmol) was reacted and purified according to the general procedure to give the title compound as a clear, colorless oil (2.3 mg, 55% yield) **$^{1}$H NMR** (600 MHz, Chloroform-*d*) δ 7.35 – 7.25 (m, 6H), 7.24 – 7.16 (m, 4H), 5.05 (d, *J* = 1.4 Hz, 1H), 4.97 (d, *J* = 1.4 Hz, 1H), 3.93 (s, 1H), 3.61 (t, *J* = 6.7 Hz, 2H), 2.35 (dd, *J* = 16.9, 9.3 Hz, 1H), 2.28 (d, *J* = 9.2 Hz, 1H), 2.10 – 2.01 (m, 2H), 2.00 – 1.97 (m, 2H), 1.74 – 1.61 (m, 3H), 1.35 – 1.16 (m, 13H). **HRMS** calc'd for $C_{30}H_{38}O_2Cl$ $[M+Cl]^-$ : 465.25658, found 465.25703.

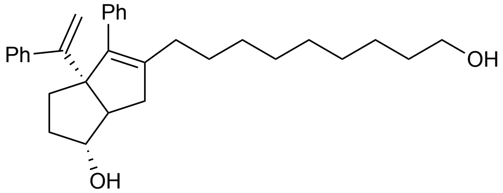

***Exo*-5-(9-hydroxynonyl)-4-phenyl-3a-(1-phenylvinyl)-1,2,3,3a,6,6a-hexahydropentalen-1-ol (3f): S1f** (9.4 mg, 0.02 mmol) was reacted and purified according to the general procedure to give the title compound as a clear, colorless oil (8.2 mg, 96% yield.) **$^{1}$H NMR** (600 MHz, Chloroform-*d*) δ 7.39 – 7.26 (m, 5H), 7.24 – 7.14 (m, 5H), 5.05 (d, *J* = 1.4 Hz, 1H), 4.97 (d, *J* = 1.4 Hz, 1H), 3.93 (s, 1H), 3.62 (t, *J* = 9.4 Hz, 2H), 2.34 (dd, *J* = 16.7, 9.4 Hz, 1H), 2.27 (d, *J* = 9.3 Hz, 1H), 2.12 – 2.03 (m, 2H), 2.01 – 1.96 (m, 1H), 1.74 – 1.58 (m, 3H), 1.34 – 1.17 (m, 18H). **$^{13}$C NMR** (600 MHz, Chloroform-*d*) δ 154.6, 144.2, 141.1, 139.1, 137.4, 129.7, 127.71, 127.70, 127.59, 126.64, 126.57, 115.0, 82.1, 69.3, 63.1, 55.8, 40.2, 34.0, 32.8, 32.1, 29.7, 29.6, 29.5, 29.4, 29.3, 27.8, 25.7. **HRMS** calc'd for $C_{31}H_{40}O_2Cl$ $[M+Cl]^-$ : 479.27223, found 479.27260.

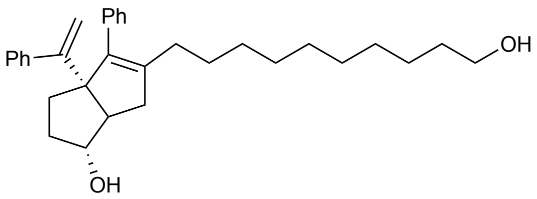

***Exo*-5-(10-hydroxydecyl)-4-phenyl-3a-(1-phenylvinyl)-1,2,3,3a,6,6a-hexahydropentalen-1-ol (3g): S1g** (81.4 mg, 0.16 mmol) was reacted and purified according to the general procedure to give the title compound as a clear, colorless oil (62.0 mg, 83% yield). **$^1$H NMR** (600 MHz, Chloroform-*d*) δ 7.35 – 7.25 (m, 5H), 7.24 – 7.16 (m, 5H), 5.05 (d, *J* = 1.4 Hz, 1H), 4.97 (d, *J* = 1.4 Hz, 1H), 3.93 (s, 1H), 3.61 (t, *J* = 6.7 Hz, 2H), 2.34 (dd, *J* = 16.9, 9.4 Hz, 1H), 2.27 (d, *J* = 9.2 Hz, 1H), 2.11 – 2.00 (m, 3H), 1.73 – 1.62 (m, 3H), 1.54 (dq, *J* = 8.2, 6.7 Hz, 2H), 1.34 – 1.17 (m, 17H). **LRMS [APCI]** calc'd for $C_{32}H_{41}O_2$ $[M-H]^-$ : 457.3, found 457.2.

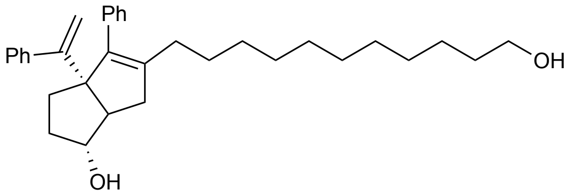

***Exo*-5-(11-hydroxyundecyl)-4-phenyl-3a-(1-phenylvinyl)-1,2,3,3a,6,6a-hexahydropentalen-1-ol (3h): S1h** (7.7 mg, 0.015 mmol) was reacted and purified according to the general procedure to give the title compound as a clear, colorless oil (5.4 mg, 78% yield). **$^1$H NMR** (600 MHz, Chloroform-*d*) δ 7.34 – 7.25 (m, 4H), 7.24 – 7.16 (m, 5H), 5.05 (d, *J* = 1.4 Hz, 1H), 4.96 (d, *J* = 1.3 Hz, 1H), 3.93 (s, 1H), 3.61 (t, *J* = 6.7 Hz, 2H), 2.33 (dd, *J* = 17.2, 9.8 Hz, 1H), 2.26 (d, *J* = 9.3 Hz, 1H), 2.09 – 1.99 (m, 1H), 2.02 – 1.99 (m, 5H), 1.72 – 1.61 (m, 3H), 1.57 – 1.51 (m, 2H), 1.40 – 1.11 (m, 10H). **LRMS [APCI]** calc'd for $C_{33}H_{43}O_2$ $[M-H]^-$ : 471.3, found 471.0.

### II. Supplemental Figures S1-S9

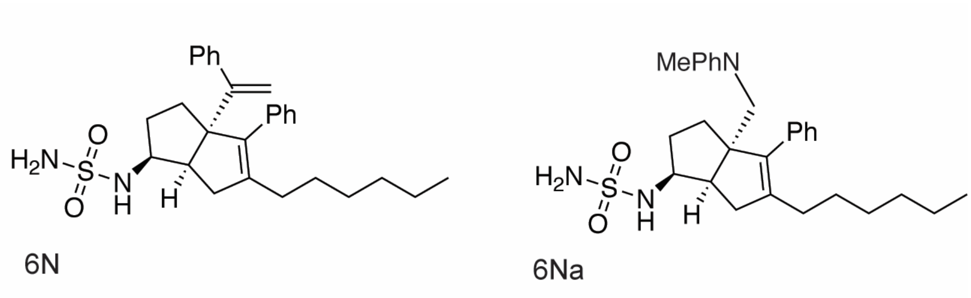

**Figure S1. Chemical structures of compounds 6N and 6Na**.

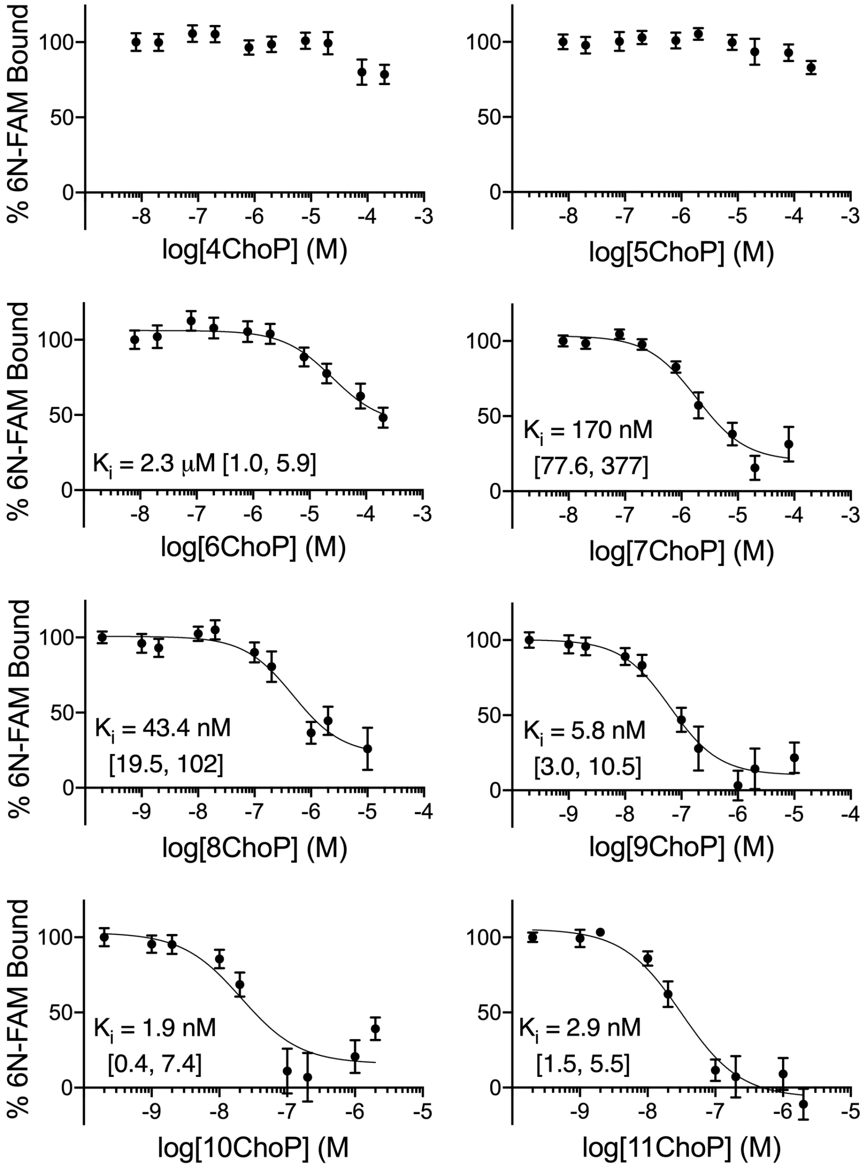

**Figure S2. Fluorescence polarization competition binding curves: phosphorylcholines.** Each point is the mean +/- SEM from two experiments, each with four technical replicates. *Insets* are calculated $K_i$ values with 95% confidence intervals in square brackets.

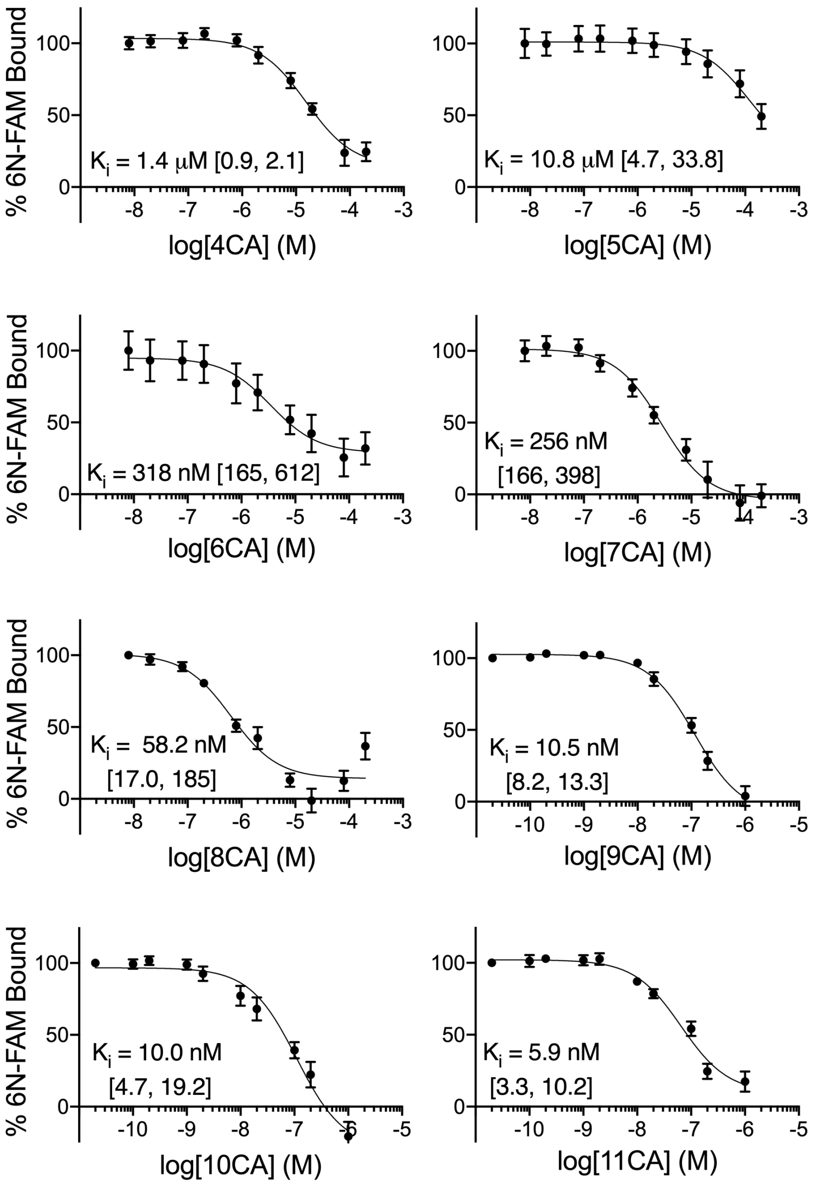

**Figure S3. Fluorescence polarization competition binding curves: carboxylic acids**. Each point is the mean +/- SEM from two experiments, each with four technical replicates. *Insets* are calculated $K_i$ values with 95% confidence intervals in square brackets.

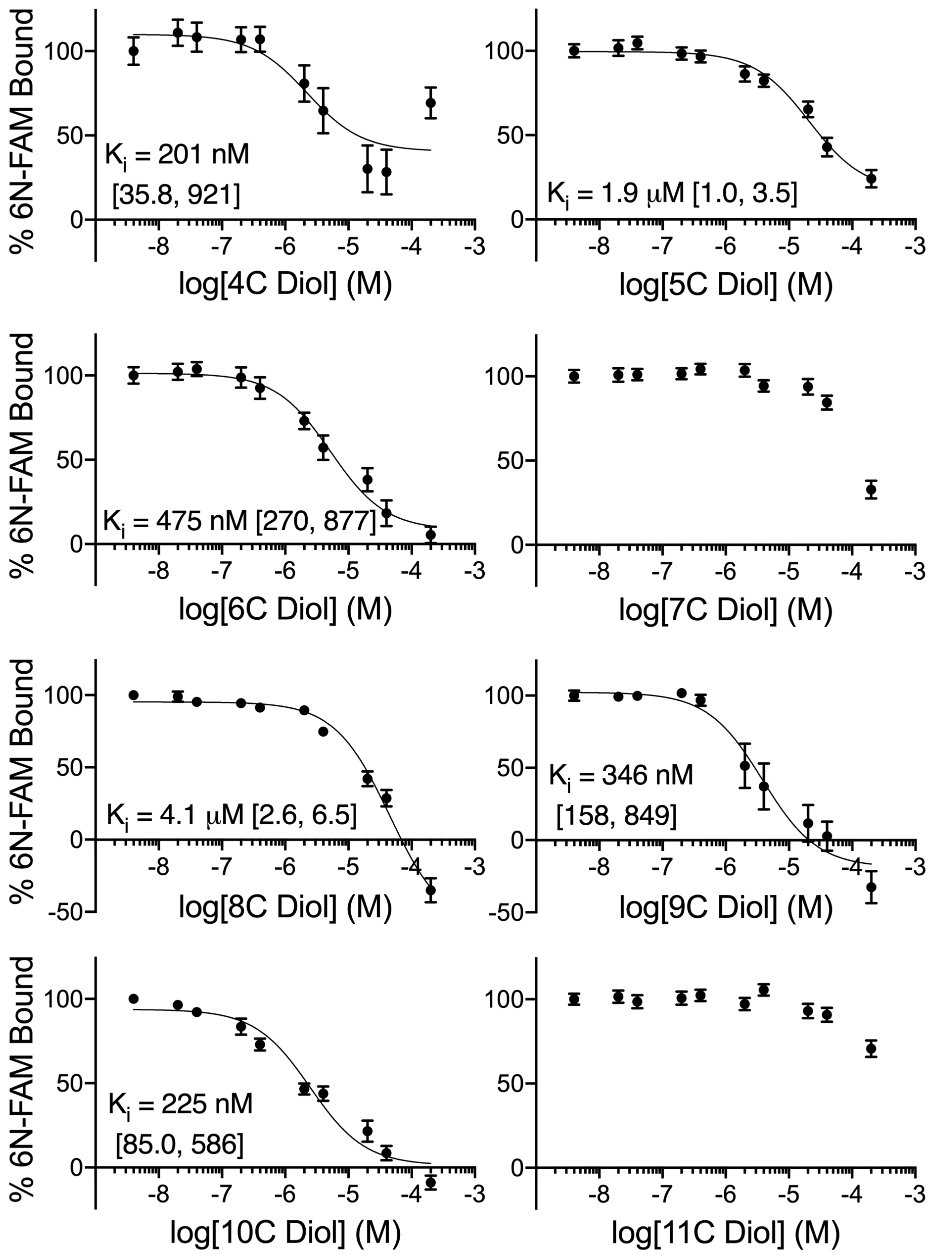

**Figure S4. Fluorescence polarization competition binding curves: diols**. Each point is the mean +/- SEM from two experiments, each with four technical replicates. *Insets* are calculated $K_i$ values with 95% confidence intervals in square brackets.

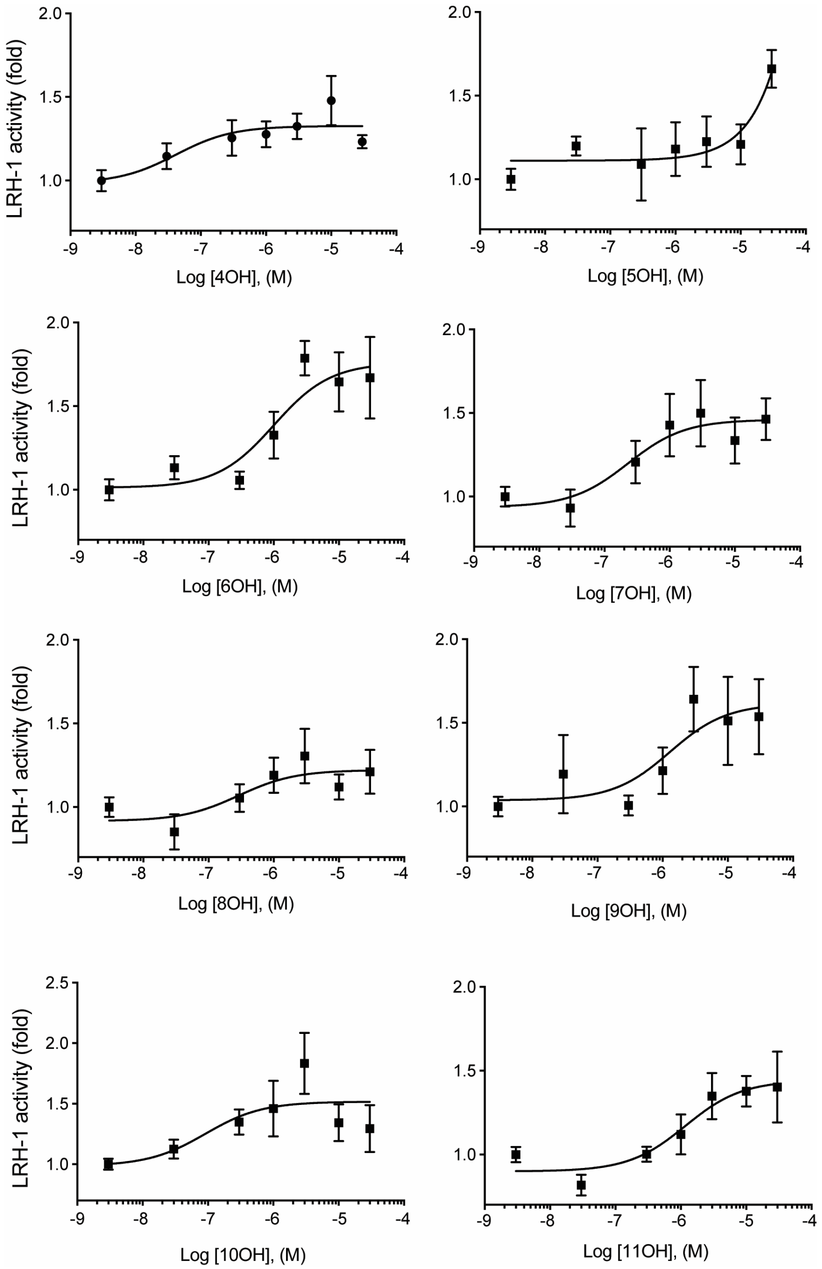

**Figure S5. Luciferase reporter assays with diols.** Hela cells were treated with the indicated concentration of each compound. Each point represents the mean ± SEM from two experiments conducted in triplicate.

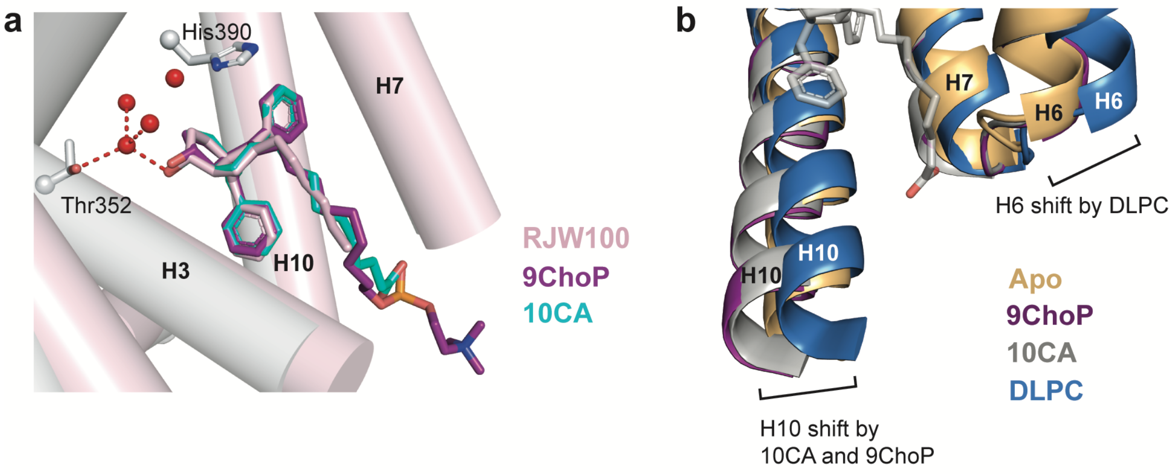

**Figure S6. Structural analyses. a,** Superposition of RJW100 (pink), 10CA (cyan), and 9ChoP (purple) ligands shows the 6HP cores of all three ligands adopt nearly identical positions. 10CA backbone is shown in grey; RJW100 backbone shown in pink. **b**, Superposition of LRH-1 structures in the apo state (PBD 4PLD, pale orange), LRH-1-DLPC (4DOS, blue), LRH-1-10CA (light grey), and LRH-1-9ChoP (purple). This is a close view of the mouth of the pocket, with helices 2 and 3 omitted to view conformational changes at helices 6 and 10. DLPC displaces helix 6 relative to apo-LRH-1, whereas 10CA and 9ChoP displace helix 10. The 10CA ligand (grey sticks) is shown for perspective. *H,* helix.

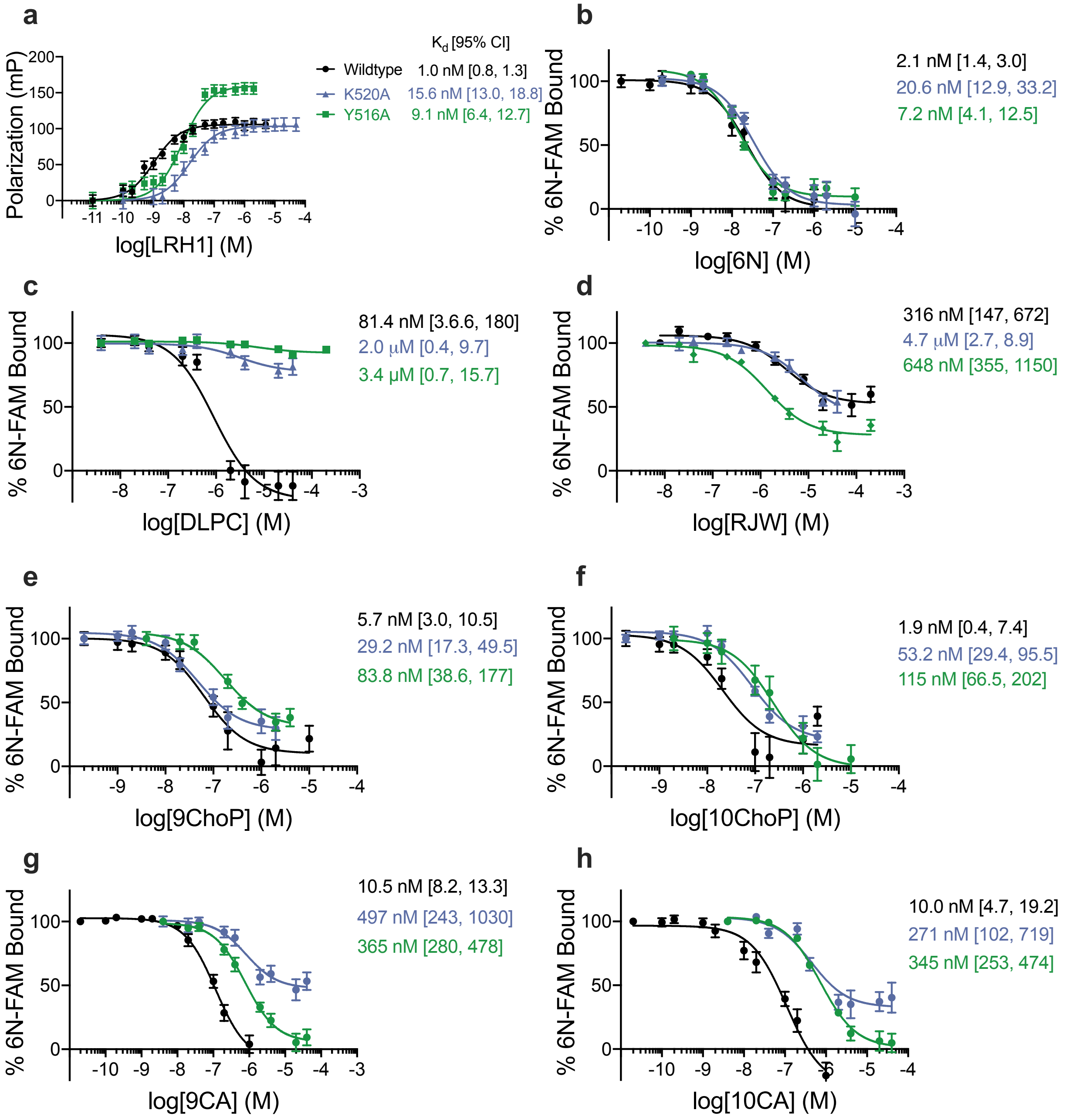

**Figure S7. Fluorescence polarization competition binding curves: LRH-1 mutants. a**, Forward binding curve for the 6N-FAM probe for wildtype *versus* mutant LRH-1. Each point is the mean +/- SEM from three experiments, each conducted in triplicate. *Insets* show $K_d$ values, with 95% CI in square brackets. **b-h**, Competition curves. Each point is the mean +/- SEM for two experiments conducted in quadruplicate. *Insets* are the calculated $K_i$ values, with 95% confidence intervals in square brackets.

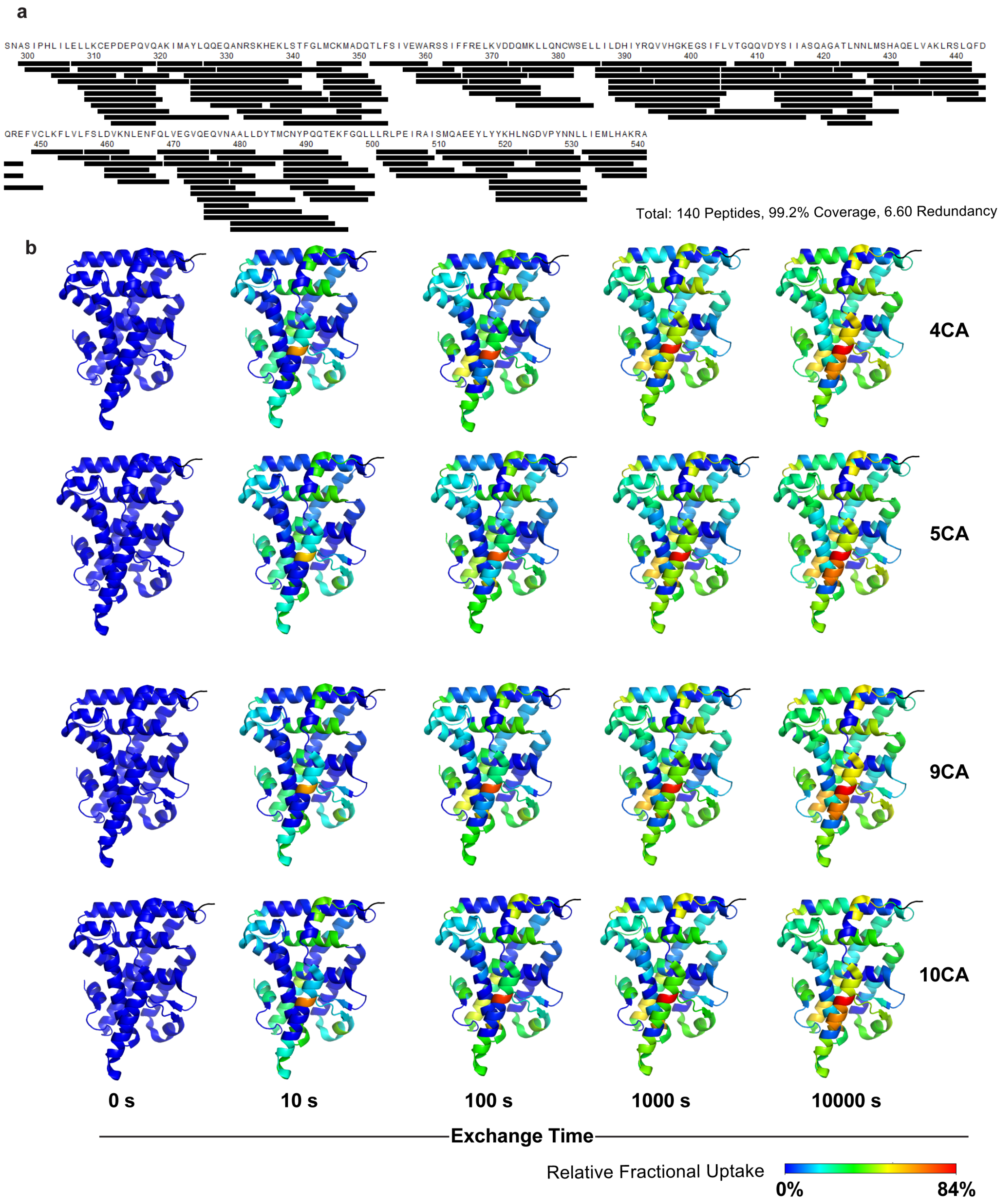
a
SNASIPHLILELLKCEPDEPQVQAKIMAYLQQEQANRSKHEKLSTFGLMCKMADQTLFSIVEWARSSIFFRELKVDDQMKLLQNCWSELLILDHIYRQVVHGKEGSIFLVTGQQVDYSIIASQAGATLNNLMSHAQELVAKLRSLQFD
300 310 320 330 340 350 360 370 380 390 400 410 420 430 440
QREFVCLKFLVLFSLDVKNLENFQLVEGVQEQVNAALLDYTMCNYPQQTEKFGQLLLRLPEIRAISMQAEEYLYYKHLNGDVPYNNLLIEMLHAKRA
450 460 470 480 490 500 510 520 530 540
Total: 140 Peptides, 99.2% Coverage, 6.60 Redundancy
b
4CA
5CA
9CA
10CA
0 s
10 s
100 s
1000 s
10000 s
Exchange Time
Relative Fractional Uptake
0%
84%

**Figure S8. Peptide coverage and deuterium uptake in HDX-MS experiments.** **a**, Sequence map of peptides identified in the experiments show 99% coverage of LRH-1 LBD. **b**, Relative fractional deuterium uptake over time for each HDX experiment, mapped onto LRH-1 LBD structure (PDB: 4DOS). The relative uptake is colored along the continuous gradient from blue to red, with red indicating the greatest degree of $D_2O$ exchange.

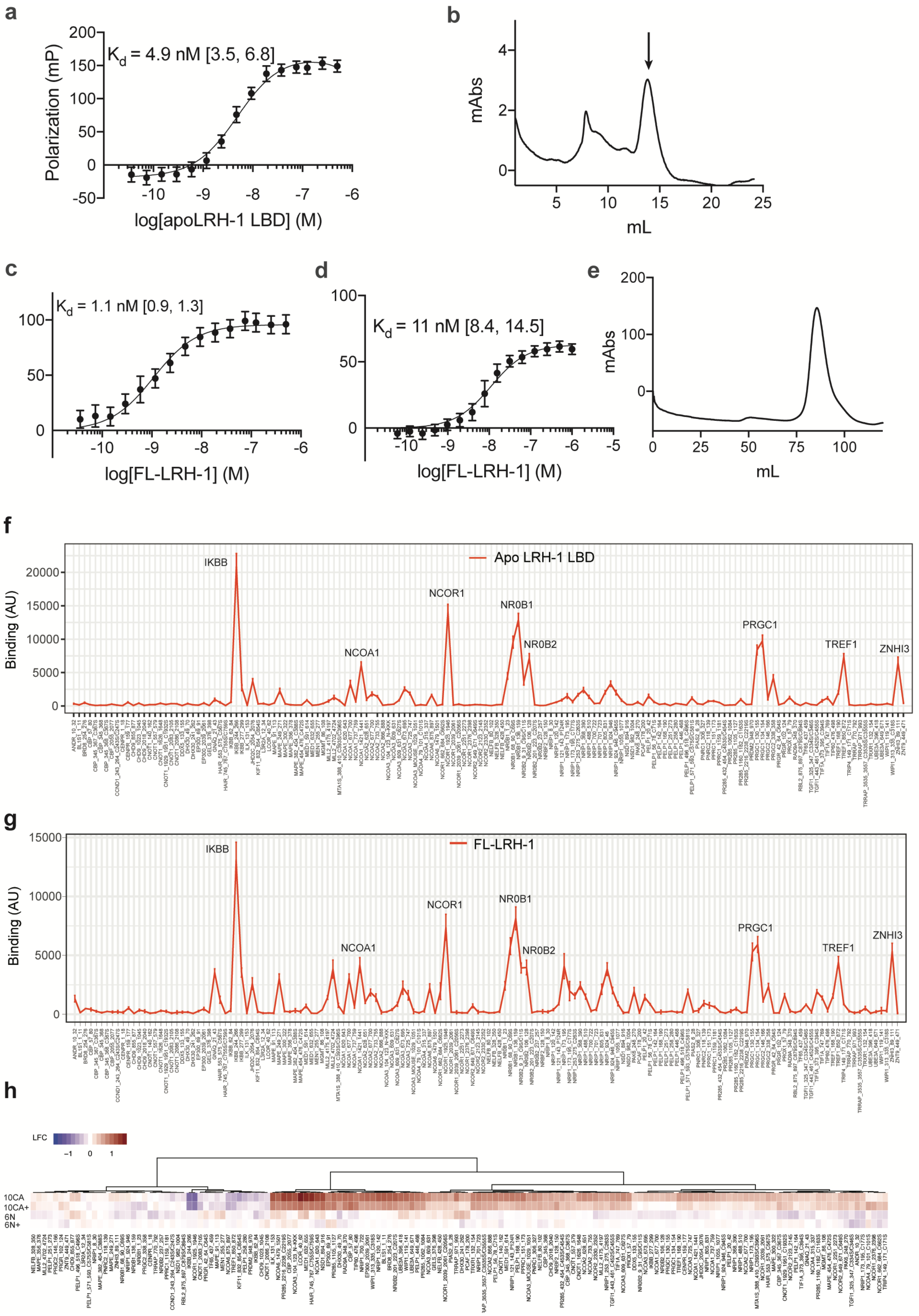
a
Kd = 4.9 nM [3.5, 6.8]
Polarization (mP)
log[apoLRH-1 LBD] (M)
b
mAbs
mL
c
Kd = 1.1 nM [0.9, 1.3]
log[FL-LRH-1] (M)
d
Kd = 11 nM [8.4, 14.5]
log[FL-LRH-1] (M)
e
mAbs
mL
f
Apo LRH-1 LBD
Binding (AU)
IKBB
NCOA1
NCOR1
NR0B1
NR0B2
PRGC1
TREF1
ZNHI3
g
FL-LRH-1
Binding (AU)
IKBB
NCOA1
NCOR1
NR0B1
NR0B2
PRGC1
TREF1
ZNHI3
h
LFC
10CA
10CA+
6N
6N+

**Figure S9. Protein purification and MARCoNI. a**, Binding curve of the 6N-FAM probe to apo LRH-1 LBD. Each point represents the mean ± SEM from one experiment conducted in duplicate. *Inset* indicates the $K_d$, with the 95% confidence interval in square brackets. **b**, Profile from size exclusion chromatography showing the pure protein eluting at ~14 ml (arrow). **c**, Binding curve of the 6N-FAM probe to FL-LRH-1. Each point represents the mean ± SEM from one experiment conducted in duplicate. *Inset* indicates the $K_d$, with the 95% confidence interval in square brackets. **d**, Fluorescence polarization binding curve showing the association between FL-LRH-1 and the LRH-1 response element on the CYP7A1 promoter. *Inset* indicates the $K_d$, with the 95% confidence interval in square brackets. **e**, Size exclusion chromatogram for FL-LRH-1, with clear elution peak at ~85 ml. **f,** Coregulator recruitment by apo LRH-1 LBD, showing recruitment of coregulator peptides, including previously identified LRH-1-interacting motifs from NR0B1, NR0B2, and NCOR1, as well as the novel interactor, IκBβ. **g**, FL-LRH-1 demonstrates a similar MARCoNI trace to apo LRH-1 LBD when not complexed with agonist. **h**, Heatmap comparing coregulator recruitment by 10CA and 6N to FL-LRH-1. "10CA+" and "6N+" refer to the addition of excess ligand immediately before the MARCoNI versus only complexing with ligand before size exclusion chromatography (see Methods).

### III. Supplemental Tables S1-S4

**Table S1. Summary of linker lengths and key biological parameters for LRH-1 agonists.**

**Phosphorylcholines**

| Linker length (# Carbons) | Linker length (Å) | log $K_i$ (M) Mean, [95% CI] | $EC_{50}$ (µM)* Mean +/- SEM | $E_{max}$* (Mean Fold vs DMSO +/- SEM) |
|---|---|---|---|---|
| 4 | 10.9 | cnc | cnc | cnc |
| 5 | 12.4 | cnc | cnc | cnc |
| 6 | 13.9 | -5.6 [-6.0, -5.2] | cnc | cnc |
| 7 | 15.5 | -6.8 [-7.1, -6.4] | cnc | cnc |
| 8 | 17.0 | -7.4 [-7.7, -7.0] | >30 | cnc |
| 9 | 18.6 | -8.2 [ -8.5, -7.9] | 7 +/- 2 | 2.3 +/- 0.2 |
| 10 | 20.1 | -8.7 [-9.4, -8.1] | 5 +/- 2 | 2.1 +/-0.1 |

| | | | | |
|---|---|---|---|---|
| 11 | 21.6 | -8.5 [-8.8, -8.3] | 5.1 +/- 0.5 | 1.4 +/- 0.06 |

**Carboxylic acids**

| Linker length (# Carbons) | Linker length (Å) | log $K_i$ (M) Mean, [95% CI] | $EC_{50}$ (μM)* Mean +/- SEM | $E_{max}$* (Mean Fold vs DMSO +/- SEM) |
|---|---|---|---|---|
| 4 | 7.6 | -5.8 [-6.0, -5.7] | cnc | cnc |
| 5 | 9.1 | -5.0 [-5.3, -4.4] | cnc | cnc |
| 6 | 10.7 | -6.5 [-6.8, -6.2] | 9 +/- 4 | 2 +/- 0.1 |
| 7 | 12.2 | -6.6 [-6.8, -6.4] | 4 +/- 3 | 1.6 +/- 0.1 |
| 8 | 13.8 | -7.2 [-7.7, -6.7] | 4 +/- 3 | 2.1 +/- 0.1 |
| 9 | 15.3 | -8.0 [-8.1, -7.9] | 1.8 +/- 0.7 | 2.5 +/- 0.1 |
| 10 | 16.8 | -8.0 [-8.3, -7.7] | 0.4 +/- 0.2 | 2.3 +/- 0.2 |
| 11 | 18.4 | -8.2 [-8.5, -8.0] | 0.3 +/- 0.2 | 1.9 +/- 0.1 |

**Diols**

| Linker length (# Carbons) | Linker length (Å) | log $K_i$ (M) Mean, [95% CI] | $EC_{50}$ (μM) Mean +/- SEM | $E_{max}$ (Mean Fold vs DMSO +/- SEM) |
|---|---|---|---|---|
| 4 | 7.6 | -6.7 [-7.4, -6.0] | 0.4 +/-0.5 | 1.44 +/- 0.07 |
| 5 | 9.1 | -5.7 [-6.0, -5.5] | cnc | cnc |
| 6 | 10.7 | -6.3 [-6.6, -6.1] | 1.0 +/- 0.8 | 1.8 +/- 0.1 |
| 7 | 12.2 | cnc | 0.2 +/- 0.3 | 1.46 +/-0.09 |
| 8 | 13.8 | -5.4 [-5.6, -5.2] | 0.7 +/- 1 | 1.4 +/- 0.1 |
| 9 | 15.3 | -6.5 [-6.8, -6.1] | 1 +/- 2 | 1.6 +/-0.2 |
| 10 | 16.8 | -6.6 [-7.1, -6.2] | 0.1+/-0.2 | 1.6 +/- 0.1 |
| 11 | 18.4 | cnc | 1 +/-1 | 1.4 +/- 0.1 |

* Values previously reported (Flynn *et al*, 2018)

**Table S2. Sequences of primers used for qRT-PCR.**

| Gene | Forward Primer (5' → 3') | Reverse Primer (5' → 3') |
|---|---|---|
| hLRH-1 | CTTTGTCCCGTGTGTGGAGAT | GTCGGCCCTTACAGCTTCTA |
| Tbp | TGCACAGGAGCCAAGAGTGAA | CACATCACAGCTCCCCACCA |
| m/hLRH-1 | GTGTCTCAAT TTAAAATGGTGAATTACTCC TATGATGAAG | AATAAGTTTGGGC CAATGTACAA GAGAGACAGG |
| Cyp11a1 | GCTGGAAGGTGTAGCTCAGG | CACTGGTGTGGAACATCTGG |
| Cyp11b1 | primers purchased from QuantiTect, Qiagen (NM_001033229, catalog # QT01198575) | |
| Cyp8b1 | TTGCAAATGCTGCCTCAACC | TAACAGTCGCACACATGGCT |
| IL-10 | GCCTTATCGGAAATGATCCAGT | GCTCCACTGCCTTGCTCTTATT |
| Rplp0 (36B4) | GAAACTGCTGCCTCACATCCG | GCTGGCACAGTGACCTCACAC |

**Table S3. Modified colitis disease activity score.**

| Scores | 0 | 1 | 2 | 3 | 4 |
|---|---|---|---|---|---|
| Weight loss (%) | 0 | >0% | ≥5% | ≥10% | ≥15% |
| Diarrhea | Normal | Mild | Loose | Moderate | Liquid |
| Hematochezia | None | FOBT± | FOBT+ | Blood++ | Blood+++ |
| Sick appearance | Normal | Mild uncleanness | Moderate uncleanness | Hunched, Slow moving | Lethargic |

**Table S4. Gene expression changes in the livers of mice expressing hLRH-1.** Mice were injected with AAV8-hLRH-1 and subsequently treated with three concentrations of 10CA or vehicle (Veh). In the table, the treatment groups are referred to as "Low" (0.1 mg/ kg), "Med" (1 mg/ kg) and "High" (10 mg/ kg). Gene expression changes in the liver were measured by Nanostring, and statistical significance was measured using two-way ANOVA followed by Benjamini-Yekutieli False Discovery Rate (FDR) method, using an FDR threshold of 0.05. The "P adj" is the q-value determined from this post-hoc analysis. The cohort of control mice (not given AAV8-hLRH-1) did not exhibit a significant effect of 10CA treatment by two-way ANOVA (p = 0.31).

| Comparison | Mean Diff. | Discovery? | P adj |
|---|---|---|---|
| **Acacb** | | | |
| Veh vs. Low | 0.4984 | No | 0.1009 |
| Veh vs. Med | 1.072 | Yes | 0.0038 |
| Veh vs. High | 0.5244 | No | 0.1009 |
| **Cyp7a1** | | | |
| Veh vs. Low | -0.6877 | No | 0.0893 |
| Veh vs. Med | 0.6512 | No | 0.0893 |
| Veh vs. High | -0.3018 | No | 0.3948 |
| **Cyp8b1** | | | |
| Veh vs. Low | -0.7386 | No | 0.0971 |
| Veh vs. Med | 0.2694 | No | 0.4508 |
| Veh vs. High | 0.5699 | No | 0.1498 |
| **Esr** | | | |
| Veh vs. Low | -0.4735 | No | 0.2603 |
| Veh vs. Med | -0.09738 | No | 0.8138 |
| Veh vs. High | -0.6548 | No | 0.1744 |
| **Fasn** | | | |
| Veh vs. Low | 1.038 | Yes | 0.0026 |
| Veh vs. Med | 1.545 | Yes | <0.0001 |
| Veh vs. High | 1.559 | Yes | <0.0001 |

| | | | |
|---|---|---|---|
| **Gck** | | | |
| Veh vs. Low | -0.01708 | No | >0.9999 |
| Veh vs. Med | 0.2463 | No | 0.7401 |
| Veh vs. High | 0.5528 | No | 0.3322 |
| **Gls2** | | | |
| Veh vs. Low | -0.3198 | No | 0.9132 |
| Veh vs. Med | 0.04026 | No | 0.9512 |
| Veh vs. High | 0.1887 | No | 0.9132 |
| **Gnmt** | | | |
| Veh vs. Low | -0.2454 | No | 0.5452 |
| Veh vs. Med | 0.2197 | No | 0.5452 |
| Veh vs. High | 0.367 | No | 0.5452 |
| **Hnf-3beta** | | | |
| Veh vs. Low | -0.2915 | No | 0.6182 |
| Veh vs. Med | 0.3803 | No | 0.6182 |
| Veh vs. High | -0.1574 | No | 0.6763 |
| **Hnf-4alpha** | | | |
| Veh vs. Low | -0.1407 | No | >0.9999 |
| Veh vs. Med | 0.3478 | No | 0.9693 |
| Veh vs. High | -0.01357 | No | >0.9999 |
| **Mdr2** | | | |
| Veh vs. Low | -0.2176 | No | 0.7822 |
| Veh vs. Med | 0.1109 | No | 0.7822 |
| Veh vs. High | 0.3831 | No | 0.7822 |
| **Nr5a2** | | | |
| Veh vs. Low | -0.2664 | No | 0.7992 |
| Veh vs. Med | -0.1036 | No | 0.7992 |
| Veh vs. High | 0.1318 | No | 0.7992 |
| **Plk3** | | | |
| Veh vs. Low | 0.7596 | Yes | 0.0139 |
| Veh vs. Med | 0.8558 | Yes | 0.0131 |
| Veh vs. High | 0.1494 | No | 0.2314 |

| | | | |
|---|---|---|---|
| **Prox1** | | | |
| Veh vs. Low | -0.5284 | No | 0.3742 |
| Veh vs. Med | 0.003255 | No | >0.9999 |
| Veh vs. High | -0.4028 | No | 0.3742 |
| **Scarb1** | | | |
| Veh vs. Low | -0.2971 | No | 0.6041 |
| Veh vs. Med | -0.1651 | No | 0.6595 |
| Veh vs. High | -0.3624 | No | 0.6041 |
| **Scd1** | 0.7405 | Yes | 0.0319 |
| Veh vs. Med | 1.157 | Yes | 0.0015 |
| Veh vs. High | 1.133 | Yes | 0.0015 |
| **Shp** | | | |
| Veh vs. Low | -0.1944 | No | 0.7165 |
| Veh vs. Med | 0.1395 | No | 0.7165 |
| Veh vs. High | -0.4466 | No | 0.6003 |
| **Srebpf** | | | |
| Veh vs. Low | 0.5762 | No | 0.0961 |
| Veh vs. Med | 0.653 | No | 0.0883 |
| Veh vs. High | 0.6995 | No | 0.0883 |
| **TGFbeta** | | | |
| Veh vs. Low | 0.1297 | No | 0.7387 |
| Veh vs. Med | 0.1945 | No | 0.7387 |
| Veh vs. High | -0.1505 | No | 0.7387 |
